# Chimeric infective particles expand species boundaries in phage inducible chromosomal island mobilization

**DOI:** 10.1101/2025.02.11.637232

**Authors:** Lingchen He, Jonasz B. Patkowski, Laura Miguel-Romero, Christopher H S Aylett, Alfred Fillol-Salom, Tiago R. D. Costa, José R Penadés

## Abstract

Some mobile genetic elements spread among unrelated bacterial species through unknown mechanisms. Recently, we discovered that identical capsid-forming phage-inducible chromosomal islands (cf-PICIs), a new family of phage satellites, are present across multiple species and genera, raising questions about their widespread dissemination. Here we have identified and characterized a new biological entity enabling this transfer. Unlike other satellites, cf-PICIs produce their own capsids and package their DNA, relying solely on phage tails for transfer. Remarkably, cf-PICIs release non-infective, tail-less capsids containing their DNA into the environment. These subcellular entities then interact with phage tails from various species, forming chimeric particles that inject DNA into different bacterial species depending on the tail present. Additionally, we elucidated the structure of the tail-less cf-PICIs and the mechanism behind their unique capsid formation. Our findings illuminate novel mechanisms used by satellites to spread in nature, contributing to bacterial evolution and the emergence of new pathogens.

## INTRODUCTION

Horizontal gene transfer (HGT) between bacterial species is crucial for bacterial evolution, adaptation to different environments, the spread of antibiotic resistance, and the emergence of new pathogens. Understanding these processes is essential for developing strategies to combat the antibiotic resistance crisis and mitigate the impact of bacterial pathogens on human health and the environment.

In recent years, we have delved into the biology of a new family of mobile genetic elements (MGEs), the phage-inducible chromosomal islands (PICIs).^1^ These are small (∼10-15 kb), chromosomally integrated elements that are extremely widespread in nature, being present in more than 200 species, with many strains containing two or more of these elements.^2^ PICIs are intimately related to certain temperate (helper) phages, whose life cycles they parasitize. In the presence of active helper phages, and once induced, PICIs replicate extensively, then excise from the bacterial chromosome and are efficiently packaged into infectious particles composed of helper phage virion proteins, which mediate the high intra-species transfer of these genetic elements.^1^

While PICIs possess captivating biology that facilitates their dissemination in nature, they also hold great importance in the biology of their bacterial hosts. Clinically, PICIs carry and disseminate an impressive array of virulence and resistance genes, including superantigens, genes involved in host adaptation, and antimicrobial resistance determinants that can ultimately transform a non-pathogenic bacterial strain into a pathogenic one.^1^ Evolutionarily, PICIs engage in two highly versatile and potent mechanisms of gene transfer called PICI lateral transduction and PICI lateral cotransduction, which promote astonishing chromosomal mobility.^3^ In conjunction with helper phage-mediated lateral transduction,^4^ they can mobilize up to 70% of the bacterial chromosome in a single event.^5^ Additionally, PICIs participate in generalized transduction (GT).^6^ Ecologically, since PICIs also carry an important repertoire of immune systems,^7^ they exert control over horizontal gene transfer, either facilitating or limiting it. They also have a significant impact on the biology of other MGEs, including phages^8^ and plasmids.^9^

Recently, one question that piqued the curiosity of our lab was how these elements emerge in different species. One possibility is that some species obtained PICIs from other species through HGT. Although some reports have demonstrated that the prototypical members of the PICIs, the Staphylococcal pathogenicity islands (SaPIs), can be mobilized in the lab to other Staphylococcal species or even to *Listeria monocytogenes*,^10,11^ the relevance of these processes *in vivo* remains unclear. In fact, genomic analyses of clinical isolates from Staphylococcal species or other genera have not revealed the presence of identical SaPIs in non-*S. aureus* species. Therefore, it is currently assumed that PICIs have a low probability (if any) of being naturally mobilized between different species.

The situation changed dramatically with the discovery of the capsid-forming PICIs (cf-PICIs). Unlike other PICIs and other phage satellites, including P4, PLE, or PICMIs, whose packaging and production of infective satellite particles depend entirely on the helper phage, cf-PICIs have the unique ability to produce small PICI-sized capsids and package PICI DNA exclusively into these capsids.^12^ Therefore, once induced, these elements only require the tails from the phage for their transfer. Interestingly, the cf-PICI-encoded genes that produce specific PICI capsids show sequence similarity to phage genes. Intriguingly, to make the cf-PICI capsids, the proteins encoded by cf-PICI genes interact exclusively with other cf-PICI-encoded proteins, not phage counterparts.^12^ How this specificity is accomplished remains a mystery.

In a previous study analyzing the distribution of cf-PICIs, we discovered that these satellites were not only the most abundant in nature, but also that a significant proportion of identical cf-PICIs could be detected in various host species.^2^ Since this does not occur with the classical PICIs or other satellites, we speculated that the fact the cf-PICIs encoded multiple genes to produce the cf-PICI capsids, instead of one or two carried by the rest of the satellites (including the classical PICIs, P4, or PLE elements), should provide an advantage in their wider distribution in nature. In other words, we hypothesized that, compared to other phage satellites, cf-PICIs had the potential to disseminate across more phylogenetically distant hosts because of their ability to produce cf-PICI capsids, which uniquely contain the cf-PICI DNA. Therefore, we initiated this study to identify the unprecedented mechanisms enabling not just their intra-but also inter-species dissemination in nature.

Here, we report that, even in the absence of a helper phage, these islands naturally produce tail-less small capsids, which contain the packaged cf-PICI genome. Once released into nature, these tail-less small capsids would interact with phage tails from different species, which are produced in excess during the phage lytic cycle. In turn, this creates chimeric infective particles capable of injecting the cf-PICI DNA into different hosts, depending on the hijacked tail. We have denominated this biological phenomenon as tail piracy. Additionally, given the significance of this novel HGT strategy, we have elucidated the molecular details behind the cf-PICI capsids, providing novel insights into the structure of these entities that drive new trajectories in bacterial evolution.

## RESULTS

### Helper phages can promote cf-PICI induction but not their transfer

During previous analyses, we realized the existence of almost identical cf-PICIs in different species. As a clear and unexpected example, we observed that one cf-PICI, EcCIGN02175, was present in five different bacterial genera and seven different bacterial species, including *Escherichia coli*, *Klebsiella pneumoniae*, *Shigella flexneri*, *Citrobacter freundii*, *Citrobacter amalonaticus*, *Enterobacter asburiae*, and *Enterobacter hormaechei* (Fig. S1; Table S1). In addition to the *E. coli* strain GN02175 carrying this element, we included in our studies *Klebsiella pneumoniae* DSM30104, which carries an island (KpCIDSM30104) very similar to EcCIGN02175 (Fig. 1A; Table S2). When these islands were compared, four variable regions were identified, two located before and after the operon encoding the genes required for the formation of mature cf-PICI capsids (packaging module), which are expected to encode immune systems.^7^ Another variable region between the islands corresponds to the gene located downstream of *alpA*, of unknown function, while the last divergent region was present in the middle of the packaging module, corresponding to the genes encoding the tail connector and tail adaptor proteins (Fig. 1A). While we did not find identical KpCIDSM30104 islands in other species, it was present in other *K. pneumoniae* strains (Table S2).

**Figure 1.**
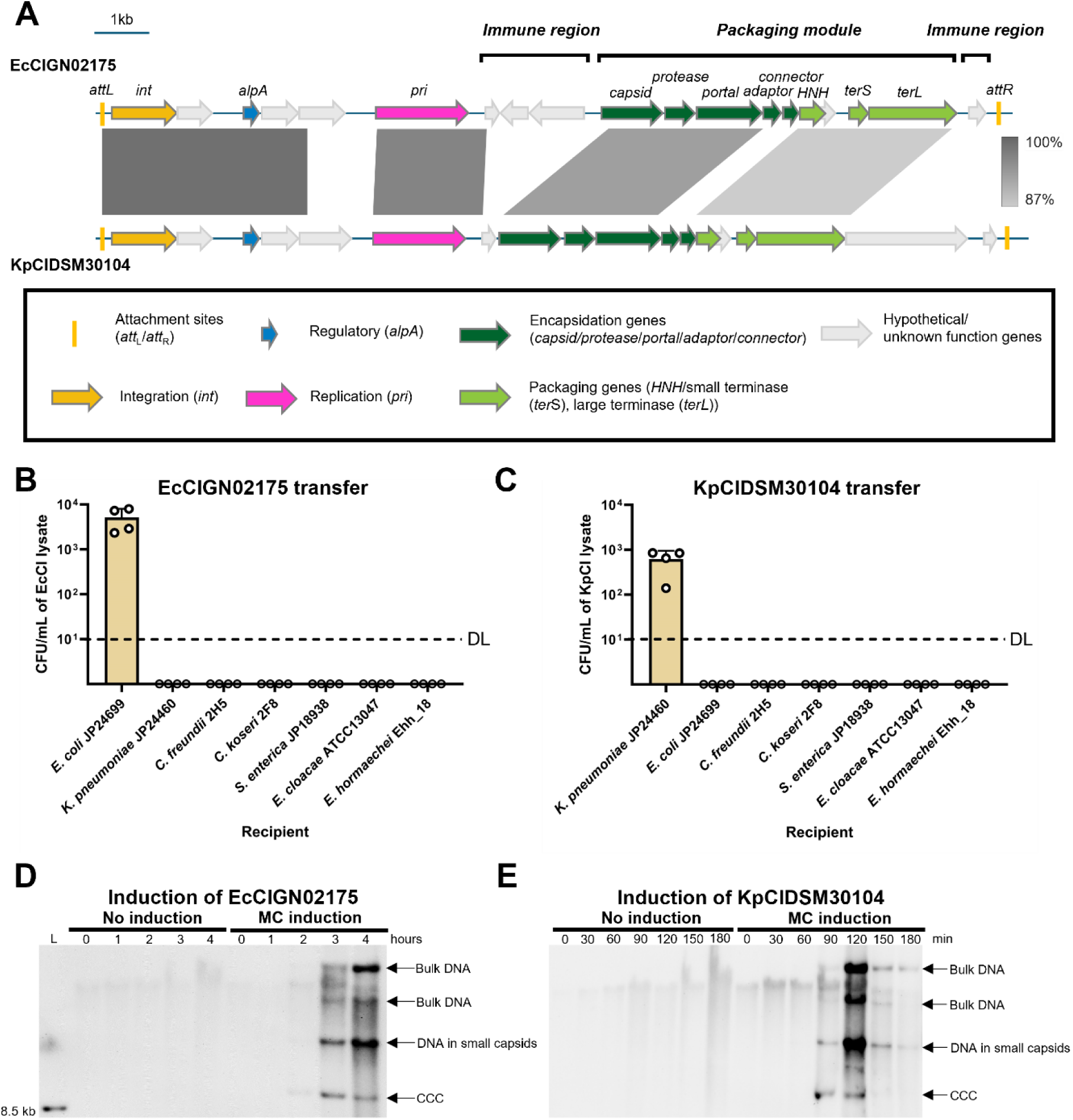
Induced cf-PICls have low intra-species transfer. **(A) A comparative map between cf-PICls EcCIGN02175 and KpCIDSM30104.** Genes are colored based on their function. Grey scales between cf-PICls indicate the regions that share similarity, identified by BLASTn. **(B, C) Transfer of EcCIGN02175 (B) or KpCIDSM30104 (C) to different bacterial species.** *E. coli* GN02175 or *K. pneumoniae* DSM30104 strains were MC-induced, and the resulting lysates were tested for transduction. The recipient strains were *E. coli* JP24699, *K. pneumoniae* JP24460, *Citrobacter freundii* 2H5, *Citrobacter koseri* 2F8, *Salmonella enterica* JP18938, *Enterobacter cloacae* ATCC13047, and *Enterobacter hormaechei* Ehh_18. Values are presented as means of colony-forming units (CFU) per milliliter of cf-PICI donor lysates. Error bars represent the standard deviation. n = 4 independent samples. DL: Detection limits. **(D, E) Induction of EcCIGN02175 or KpCIDSM30104 by resident prophages.** *E. coli* GN02175 or *K. pneumoniae* DSM30104 strains were MC induced, and samples were taken at the indicated time points (hours) for DNA analyses. DNA was separated on a 0.7% agarose gel, followed by Southern blotting analysis using specific EcCIGN02175 (D) or KpCIDSM30104 (E) probes. L: Southern blot molecular marker (DNA molecular weight marker VII; Roche). CCC: Covalently closed circular.

The presence of identical elements not only in different strains but also in different species suggested that they were horizontally mobilized. While the basis of intra-species mobilization seems clear – the helper phage provides tails after induction that allow the mobilization of the cf-PICIs to a new recipient strain – the mechanism of inter-species transfer remains unclear. One possibility is that the infective particles acquired the ability to inject their DNA into different species from the helper phages that induced these islands. To test this idea, we utilized the two natural strains carrying our model islands, *K. pneumoniae* DSM30104 and *E. coli* GN02175, which, in addition to the cf-PICIs, carried two and seven additional resident prophages, respectively. We inserted an antibiotic marker in each of the cf-PICIs, induced the resident prophages with mitomycin C (MC), and tested the transfer of the islands into different bacterial species used as recipients. Note that we included the same strains, with scarless deletions of the cf-PICI islands, as recipients to minimize the impact of the bacterial immune system (especially restriction-modification systems) in blocking HGT. In these experiments, we expected that one or some of the resident prophages would be able to induce the islands and provide the tails required for the formation of the cf-PICI infective particles. However, our initial results suggested this was not the case, as we observed low intra-species transfer of both islands (Fig. 1B and 1C). No interspecies transfer was detected, which we hypothesized was due to the lack of cf-PICI induction by the resident prophages.

To confirm that neither the *E. coli* nor the *K. pneumoniae* cf-PICIs were induced, we performed screening lysate analyses. The strains carrying the islands were induced with MC, and samples for DNA analysis were taken at different time points after induction, run on an agarose gel, and subjected to Southern blot analyses using specific probes against the cf-PICI DNAs (Fig. 1D and 1E). Unexpectedly, and contrary to our initial hypothesis, the samples obtained from the *E. coli* strain showed clear signs of cf-PICI induction and packaging, with the presence of bulk DNA (indicative of replication) and a small-sized cf-PICI band (representative of packaging into small capsids; Fig. 1D). A much clearer pattern of induction was observed when the *K. pneumoniae* cf-PICI was analyzed (Fig. 1E), suggesting that both strains contained a helper phage for the satellites. Since the strain carrying the *E. coli* cf-PICI contains multiple prophages and an additional PICI, we focused our analyses on the KpCIDSM30104 element. As previously mentioned, the *K. pneumoniae* DSM30104 strain only contained two prophages and the cf-PICI KpCIDSM30104. To identify its helper phage, we obtained derivative mutants in which prophages P1 or P2 were removed. These strains were MC-induced and analyzed as previously indicated. Deletion of phage P1 abolished cf-PICI induction (Fig. S2A and S2B) and transfer (Fig. S2C), confirming its identity as the helper. Interestingly, the analysis of a *K. pneumoniae* DSM30104 derivative strain uniquely carrying the helper phage revealed that this phage is severely affected by the island (more than 100-fold reduction), as shown by the low amount of phage particles present in the lysates containing the island compared to the lysates containing only the phage (Fig. S2D).

### cf-PICIs naturally produce tail-less particles

The previous results with KpCIDSM30104 suggested that while the island was induced, the helper phage was unable to provide (at least at very high frequency) the tails required to complete the formation of the infective particles. In support of this idea, the helper phage belongs to the *Podoviridae* family of phages, which have short tails, whereas cf-PICIs use non-contractile large tails from lambdoid phages belonging to the *Siphoviridae* family.^12^ Due to this apparent incompatibility, and to gain more insight into how the KpCIDSM30104 island was transferred, we obtained derivative mutants of the strain carrying both the island and the helper phage P1, in which either the capsid gene from the island or the capsid or tail genes from the helper phage were deleted. These strains were MC-induced, and the transfer of the island was analyzed. Importantly, the deletion of the capsid gene from the cf-PICI did not affect its transfer, while the deletion of either the capsid or the tail genes from the phage eliminated the formation of both phage and KpCIDSM30104 infective particles (Fig. S3). These results indicate that after induction, the helper phage mobilized KpCIDSM30104 via generalized transduction, a mechanism independent of the production of cf-PICI capsids.

The previous results also suggested that because KpCIDSM30104 is induced by a phage (P1) that is unable to provide tails to complete the formation of infective KpCIDSM30104 particles, tail-less KpCIDSM30104 capsids should be present in the lysate of the induced cells. To test this, the particles present in the lysates from MC-induced cultures of WT *K. pneumoniae* DSM30104, or derivative mutants in helper phage 1 (Δphage P1), in non-helper phage 2 (Δphage P2), or in KpCIDSM30104 and in phage 1 (Δcf-PICI Δphage P1), were precipitated. The packaged DNA was extracted, run on an agarose gel, and analyzed by Southern blot. As shown in Fig. 2, all lysates from strains containing both P1 and KpCIDSM30104 exhibited DNA packaged in small capsids. Importantly, the most intense bands observed in the DNA analysis from the WT strain corresponded to the packaged cf-PICI (PICI-sized DNA; Fig. 2A) and to a high molecular weight band, which represents renatured cf-PICI DNA forming concatemers due to interactions between the cohesive ends of the *cos* sites (Fig. 2B). Together, these results indicate that: i) this element blocks helper phage P1 reproduction, as previously indicated, and ii) after induction, packaging of this element is extremely efficient and occurs at high frequency. Additionally, no DNA band was observed in the strain containing only the non-helper phage P2 (Δcf-PICI Δphage P1), indicating that phage P2 is either defective or not SOS-inducible. In support of the latter option, it is similar to *E. coli* phage P2^13^, which is not induced after treatment of the P2 lysogen with MC.

**Figure 2.**
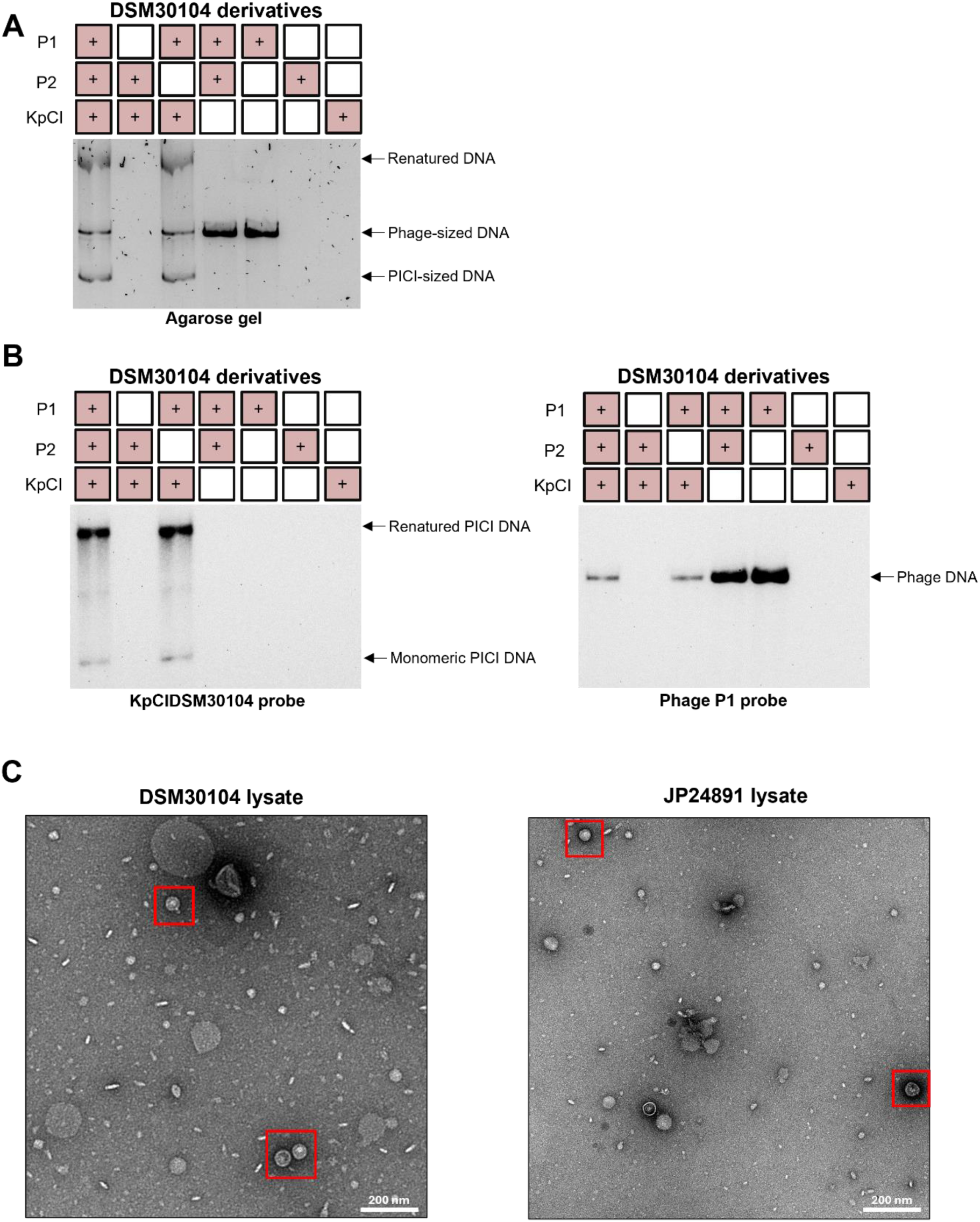
cf-PICls naturally produce tail-less particles. **(A)** Packaged DNA extracted from lysates after induction of different DSM30104 derivatives was separated on a 0.7% agarose gel. KpCI: KpCIDSM30104. ‘+’ indicates the presence of P1, P2, or KpCIDSM30104 in the donor strain. **(B)** Southern blot analyses of the samples obtained in panel A, using KpCIDSM30104 or P1-specific probes. **(C)** Electron microscopy images of tail-less cf-PICls. Left: Image showing capsids present in the lysate of the induced WT DSM30104 strain. Right: Image showing the KpCIDSM30104 capsids obtained after induction of the DSM30104 derivative mutant in P2 and in the capsid gene of P1. Capsid particles are highlighted with a red frame.

To confirm that the packaged cf-PICIs did not contain attached phage tails, we analyzed the particles obtained after induction of the WT strain, and the strain carrying the helper phage P1 with a capsid gene deletion and KpCIDSM30104 (JP24891), showing that all the observed small cf-PICI-sized particles were tail-less (Fig. 2C).

In the satellite world, this represents a scenario where the helper phage can induce the island and lyse the cells but is unable to provide the tails required for the completion of the cf-PICI infective particles. Our results reveal a novel type of interaction between satellites and phages, with some cf-PICIs potentially requiring two helper phages: one for induction and the other for completing the formation of the infective particles by providing the tails.

### The formation of hybrid infective cf-PICIs promotes intra- and inter-generic transfer

The previous results also raised an interesting question: are the tail-less cf-PICIs defective particles unable to inject their DNA into new recipient cells, or, on the contrary, are they biologically relevant? If the latter, this would represent the discovery of a new type of biological entity whose function remains to be determined.

It is important to remark here that DNA packaging of phage or satellite DNAs into large or small capsids, respectively, and phage tail formation, are two processes that occur simultaneously but independently in the bacterial cells.^14^ Once the DNAs are packaged and the capsids matured, the formed tails attach to them, creating the infective phage or satellite particles. It is also well known that the host range of a phage is primarily determined by phage tail fibers (or spikes), which initially mediate reversible and specific recognition and adsorption by susceptible bacteria.^15^ Since tails receptors are usually species (and even sometimes strains) specific, it is assumed that the phage host range is narrow. However, the fact that induced cf-PICI can create tail-less capsids opened an interesting possibility that would explain the biological relevance of these new entities: we hypothesized that the released cf-PICI capsids have the ability to interact with tail fibers from phages infecting different species, creating chimeric particles with a different host range. This would explain the inter-species transfer of the cf-PICIs.

To test this, we MC-induced the clinical isolate *K. pneumoniae* DSM30104 Δphage P2 (JP24853), which lysate mainly contains tail-less KpCIDSM30104 islands (Fig. 2B). In parallel, and since our collection of *E. coli* phages is more extensive, we induced the lysogenic strains carrying capsid mutant versions of the HK106 or HK022 prophages, which have different tails (Fig. S4). These phages were used in the previous study characterizing the cf-PICIs in *E. coli*.*^12^* The lysates obtained from these mutant prophages uniquely contain tails. Next, we mixed the *K. pneumoniae* JP24853 lysate containing cf-PICI capsids with the *E. coli* ones containing tails, in a 1:1 ratio, and tested the transfer of the KpCIDSM30104 cf-PICI to various species.

Remarkably, while no increased intra-species transfer was detected, the presence of the tails from *E. coli* phage HK022 allowed the massive inter-generic transfer of KpCIDSM30104 to *E. coli* strain C1a, with frequencies of transfer rarely seen before for other satellites (Fig. 3A). No transfer to *E. coli* was observed in absence of the extra tails, or in presence of the HK106 tails. This suggested that not all the *E. coli* phage tails are compatible with the KpCIDSM30104 capsids. While transfer to other *E. coli* strains also occurred at high frequency, no transfer was observed either when other species were used as recipient (Fig. 3B), confirming the idea that tails determine the tropism of the infective particles. However, in the presence of compatible tails, our results suggest that KpCIDSM30104 capsids create functional chimeric particles with the *E. coli* phage HK022 tails with the ability to mobilize their DNA to *E. coli* (Fig. 3B). The presence of these infective particles was confirmed by EM (Fig. 3C).

**Figure 3.**
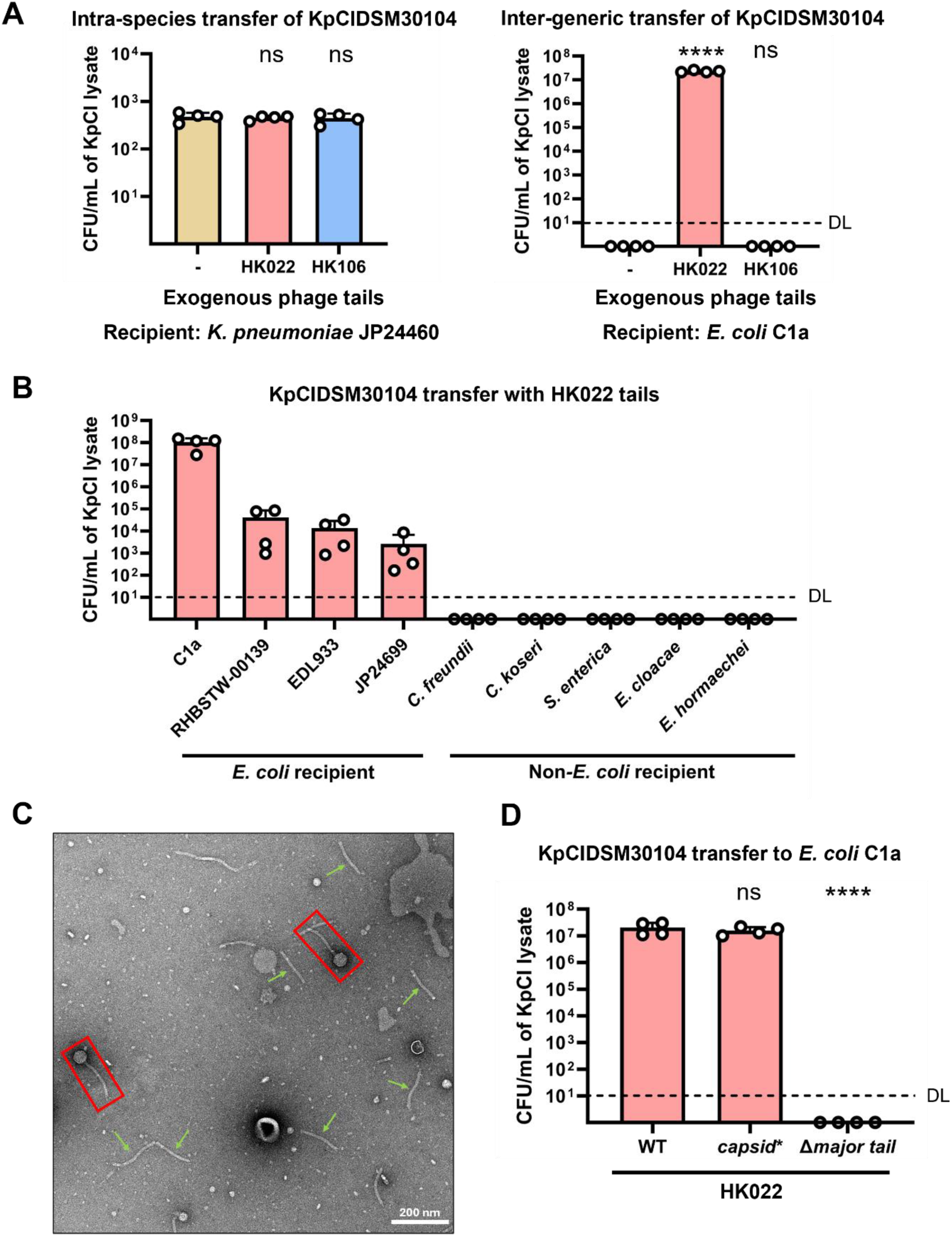
Formation of chimeric cf-PICIs promotes inter-generic transfer. **(A)** Intra- or inter-species transfer of KpCIDSM30104 by the formation of chimeric infective particles. The lysate obtained after induction of the *K. pneumoniae* DSM30104 Aphage P2 strain (JP24853) was mixed with the lysate obtained after induction of HK022 or KK106 prophage mutants in their respective capsid genes, and then the transfer of **the** KpCIDSM30104 island to *K. pneumoniae* JP24460 (left) or *E. coli* C1a (right) was evaluated. Values are presented as means of colony-forming units (CFU) per milliliter of cf-PICI donor lysates. Error bars indicate the standard deviation. A t-test was used to compare the means of control and samples with additional phage tails after Iog10 transformation, ns: P > 0.05. ****: P ≤ 0.0001 n = 4 independent samples. ‘-’ represents the control where cf-PICIs were mixed with LB. DL: Detection limits. **(B)** The chimeric particles have a host range that depends on the hijacked tails. The lysate containing the tail-less KpCIDSM30104 particles was mixed with HK022 tails, and the transfer of KpCIDSM30104 to different species was evaluated. Recipients were *E. coli* (C1a, RHBSTW-00139, EDL933, and JP24699), *Citrobacter freundii* 2H5, *Citrobacter koseri* 2F8, *Salmonella enterica* JP18938, *Enterobacter cloacae* ATCC13047, and *Enterobacter hormaechei* Ehh_18. Values are presented as means of colony-forming units (CFU) per milliliter of cf-PICI donor lysates. Error bars indicate the standard deviation, n = 4 independent samples. DL: Detection limits. **(C)** Negative stain electron microscopy image of KpCIDSM30104 after incubation with HK022 phage tails, yielding assembled virion particles (red frames). Excess free HK022 tails are also shown (green arrows). **(D)** Induction of WT phage HK022 produces tails in excess for the inter-generic transfer of KpCIDSM30104. The lysates obtained after induction of the *K pneumoniae* JP24853 strain were mixed with the lysates obtained after induction of the WT *E. coli* HK022 lysogen, or with lysates obtained after induction of the HK022 capsid and major tail mutant prophages, and the transfer of KpCIDSM30104 to *E. coli* strain C1a was evaluated. Values are presented as means of colony-forming units (CFU) per milliliter of cf-PICI donor lysates. Error bars indicate the standard deviation. After Iog10 transformation, a One-way ANOVA was conducted, followed by a Dunnett’s multiple comparisons test to compare the sample with exogenous WT HK022 lysate to other samples, n = 4 independent samples, ns: P > 0.05. ****: P ≤ 0.0001. KpCI: KpCIDSM30104. WT: wild-type.

Since in the previous experiment we used *E. coli* lysates uniquely containing tails, we tested whether these chimeric particles were also produced in the presence of functional *E. coli* phages (used again as tail donors). To do that, we repeated the same experiment, but now mixing the *K. pneumoniae* JP24853 lysate with the lysates obtained after induction of the *E. coli* strain carrying the WT version of the HK022 prophage. Remarkably, the inter-species transfer of the KpCIDSM30104 occurred at the same level than when the phage capsid mutants, but not the phage tail mutants, were used (Fig. 3D), confirming previous studies of the lab that suggested that phage tails are produced in excess during phage lytic cycle.^16^

To test if cf-PICIs can hijack tails from intact phages, phage HK022 lysate was diluted to 10^6^ PFU/mL, mixed with KpCIDSM30104 (present at 10^8^ particles/mL), and phage titration was subsequently performed. The phage titration results remained unchanged (Fig. S5A and B), indicating that cf-PICIs utilize excess tails from the phage lysates rather than appropriating tails from intact phages (Fig. 5SC). Taking together with previous experiments, these suggest using tails excessively produced by phages is a common strategy and natural scenario for cf-PICI tail-less particles to produce infective particles.

**Figure 4.**
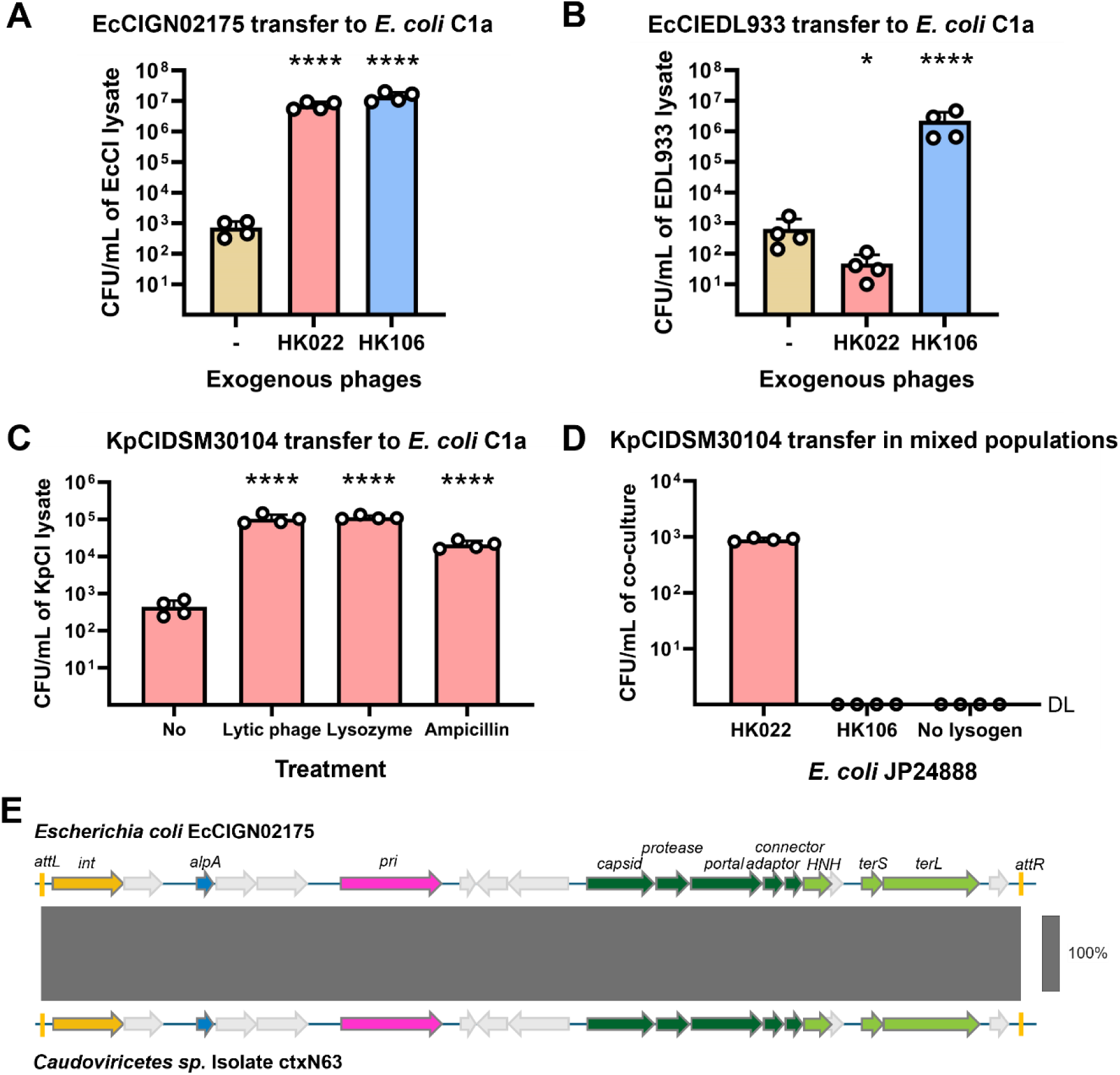
Tail-less cf-PICIs are frequently released into natural environments and transmit within microbiota. Intra-species transfer of EcCIGN02175 **(A)** or EcCIEDL933 **(B)** to C1a. Natural *E. coli* GN02175 and EDL933 strains, carrying EcCIGN02175 or EcCIEDL933 respectively, were MC-induced, and the resulting lysates were tested for transduction in the presence or absence of lysates obtained after induction of WT HK106 or HK022 lysogens. Values are presented as means of colony-forming units (CFU) per milliliter of cf-PICI donor lysates. Error bars indicate the standard deviation. ‘-’ represents the control where cf-PICIs were mixed with LB. A t-test was used to compare the data between samples with no exogenous phage (control) and other samples after Iog10 transformation. *: P ≤ 0.05. ****: P ≤ 0.0001. n = 4 independent samples. **(C)** cf-PICIs are produced in the absence of an inducing helper prophage. The non-lysogenic *K. pneumoniae* JP24871 strain, carrying KpCIDSM30104, was grown until OD600 = 0.2. Then, the strain was cultured in the presence of lytic phage K68 QB (MOI = 0.01), lysozyme (500 pg/mL), or ampicillin (50 pg/mL). As a control, non-treated cells were also grown. The resulting filtered lysates were mixed with HK022 lysates and tested for transduction to *E. coli* C1a. Values are presented as means of colony­forming units (CFU) per milliliter of cf-PICI donor lysates. Error bars indicate the standard deviation. A t-test was used to compare the data between samples with no treatment and the other samples after Iog10 transformation. ****: P ≤ 0.0001. n = 4 independent samples. **(D)** Inter-species transfer occurs naturally in mixed populations. *K. pneumoniae* JP24853, carrying both P1 and KpCIDSM30104, was co-cultured with either the *E. coli* HK022 or HK106 lysogens in the presence of the *E. coli* strain JP24888, used here as the recipient for KpCIDSM30104. Strains were mixed at a ratio of 1:1:1 for 5 hours and then plated in the presence of both kanamycin (for KpCIDSM30104) and tetracycline (for JP24888). Values are presented as means of colony-forming units (CFU) per milliliter of co-culture. Error bars indicate the standard deviation, n = 4 independent samples. DL: Detection limits. **(E)** Presence of cf-PICI EcCIGN02175 in human metagenome studies. Genes are colored according to their sequence and function. Grey scales between cf-PICI sequences indicate regions of similarity identified by BLASTn. The overall identity between both islands is 99.95%.

**Figure 5.**
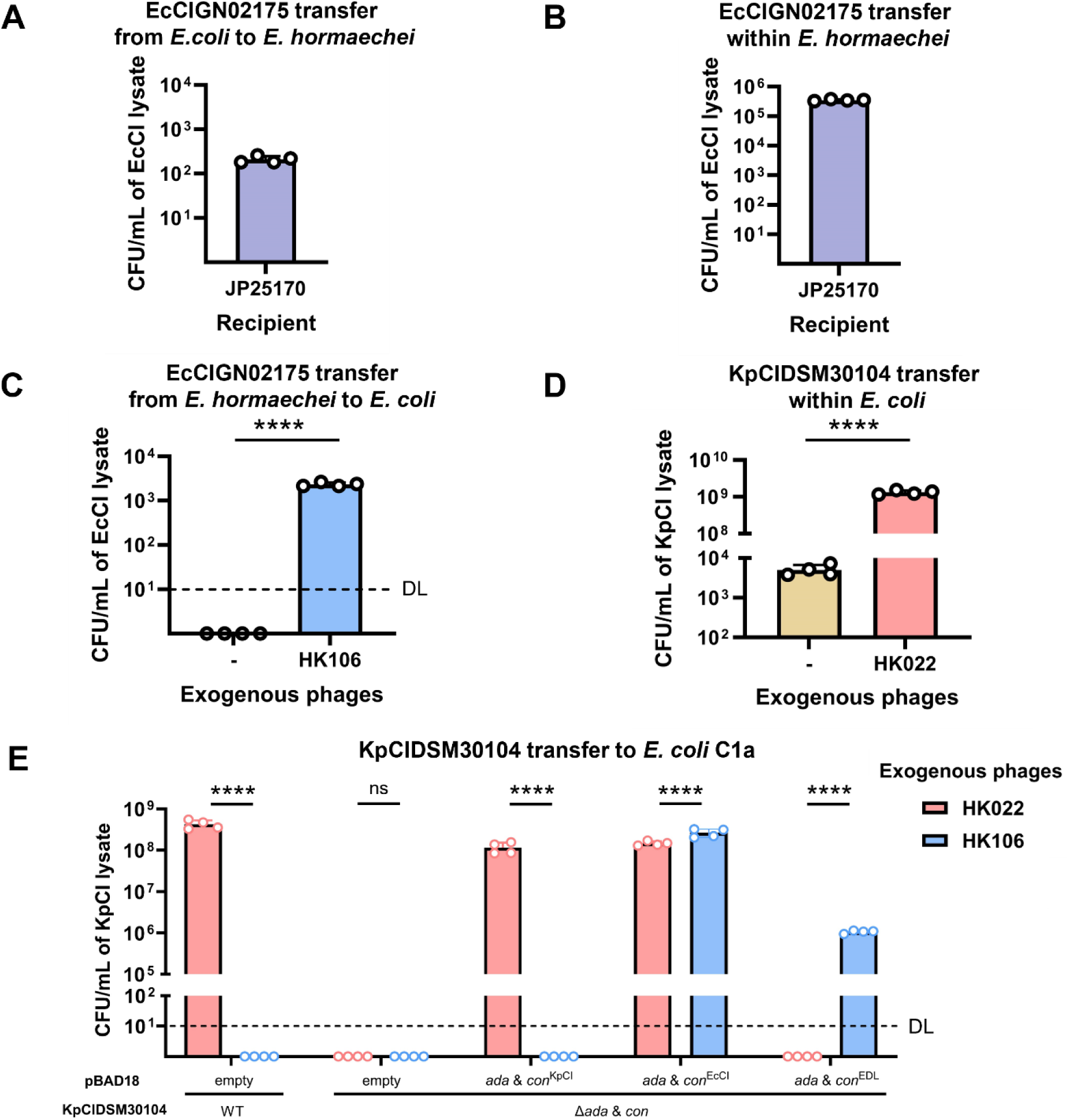
The cf-PICI-encoded tail adaptor and connector proteins determine tail specificity. **(A)** Inter-species transfer of EcCIGN02175 from *E. coli* GN02175 to *E. hormaechei* Ehh_18 JP25170 (Ehh_18 AEhCIEhh_18). The *E. coli* strain GN02175 was induced, and the lysate mixed with that obtained after MC induction of *E. hormaechei* Ehh_18 JP25170. The transfer of EcCIGN02175 is quantified. No transfer was observed in the absence of the *E. hormaechei* Ehh_18 JP25170 lysate. **(B)** Intra-species transfer of EcCIGN02175. The *E. hormaechei* Ehh_18 JP25149 strain carrying GN02175 was induced, and the transfer of the island to *E. hormaechei* Ehh_18 JP25170 (Ehh_18 AEhCIEhh_18) was analyzed. **(C)** Inter-species transfer of EcCIGN02175 from the *E. hormaechei* Ehh_18 derivative JP25149 to *E. coli* C1a was analyzed in the presence of the lysate obtained after induction of the HK106 prophage. **(D)** Intra-species transfer of KpCIDSM30104 in *E. coli.* The lysate obtained after induction of the *E. coli* GN02175 derivative JP25235 carrying KpCIDSM30104 was mixed with the lysate obtained after induction of the HIK022 prophage, and the transfer of the island was analyzed. ‘-’ represents the cf-PICI sample with no phage lysate added. In **(A), (B), (C),** and **(D),** values are presented as means of colony-forming units (CFU) per milliliter of cf-PICI donor lysates. Error bars indicate standard deviation, n = 4 independent samples. In (C) and (D), a t-test was used after a Iog10 transformation. ****: P ≤ 0.0001. **(E)** cf-PICIs evolve their adaptor and connector proteins to interact with different phage tails. The strain carrying KpCIDSM30104 mutated in its adaptor and connector genes was complemented with different adaptor and connector genes from other cf-PICIs. These strains were MC- and L-arabinose-induced. The resulting lysates were tested for KpCIDSM30104 transfer in the presence of the lysates obtained after induction of the HK022 or HK106 prophages. *E. coli* C1a was used as the recipient strain in these experiments. Values are presented as means of colony-forming units (CFU) per milliliter of cf-PICI donor lysates. Error bars indicate standard deviation. A two-way ANOVA with Sidak’s multiple comparisons test was performed after a Iog10 transformation. ****: P ≤ 0.0001. n = 4 independent samples, ada & con: adaptor and connector gene. EcCI: EcCIGN02175. KpCI: KpCIDSM30104. EDL: EcCIEDL933. DL: Detection limits.

### Formation of tail-less cf-PICI capsids occurs widely in nature

Are the results obtained with KpCIDSM30104 an exception, or conversely, do they represent the biology of the cf-PICI elements? To answer this question, we made use of two clinical *E. coli* strains naturally carrying cf-PICIs. One corresponds to the aforementioned *E. coli* GN02175. The other corresponds to the *E. coli* EDL933, which in addition to the prototypical EcCIEDL933 used in the original study characterizing the cf-PICI elements,^12^ carries 17 additional prophages.^17^ Importantly, none of these resident prophages induced the island nor provided compatible tails to the EcCIEDL933 capsids (assuming they were formed in the absence of induction). Note that just a very few EcCIEDL933 transductants were obtained using the lysate obtained after MC induction of the EDL933 strain.^12^ The *E. coli* GN02175 and EDL933 strains were MC induced, the obtained lysates were mixed with the lysates obtained after induction of the WT HK106 or HK022 prophages, and then the transfer to *E. coli* C1a was measured. Remarkably, the transfer of the EcCIGN02175 island significantly increased (nearly 4 logs) in the presence of the tails from any phages (Fig. 4A), which indicates that as occurred with KpCIDSM30104, EcCIGN02175 is able to produce tail-less particles after induction. Interestingly, and contrary to what occurred with KpCIDSM30104, which only interacted with tails from HK022, EcCIGN02175 was also able to interact with the tails provided by phage HK106, suggesting that this island has an increased ability to interact with different phages (see below).

The results with EcCIEDL933 were even more interesting. As indicated, none of the resident phages induce the island.^12^ However, upon adding tails to the lysate obtained after MC induction of the EDL933 strain, there was a significant increase in the transfer of this island (Fig. 4B). Specifically, it was observed that the tails from phage HK106, rather than those from HK022, were capable of generating infective particles. This implies that tail-less EcCIEDL933 capsids are produced at a significant level even in the absence of helper phage induction. These capsids will be liberated after the lysis of the culture by the non-inducing phages.

To test the possibility that cf-PICIs were produced in absence of an inducing phage, we made use of the non-lysogenic *K. pneumoniae* JP24871 strain, which uniquely carries the KpCIDSM30104 island. We then challenged this strain with different biological or natural compounds that naturally disrupt the cell wall: a *K. pneumoniae* lytic phage that does not induce the island (K68 QB), lysozyme, or with beta-lactam antibiotics. Note that these external stressors are found regularly in natural scenarios, like the gut, which contain lytic phages, produce lysozyme secreted by intestinal Paneth cells,^18^ and some competitors or treatments provide ampicillin. Importantly, after these challenges, in the presence of the compatible tails, the island was highly transduced to *E. coli* C1a (Fig. 4C), confirming the hypothesis that the tail-less cf-PICIs are spontaneously produced at a high level even in the absence of helper phages. These scenarios present a unique phenomenon never observed with other satellites. These also highlight the biological relevance of tail-less cf-PICIs: they represent incomplete delivery particles which have evolved to be produced without tails as a strategy which allows them later, when liberated in nature, to hijack free tails from phages of different species. In the end, this will increase the repertoire of bacterial species that can be used as recipients of the cf-PICI DNA packaged, a strategy that will promote the promiscuous transfer of the cf-PICIs in nature.

### Inter-species cf-PICI transfer occurs naturally in mixed population

The observation that cf-PICIs are massively transferred in the presence of the correct tails prompted our investigation into whether such transfer would manifest in more natural scenarios, where various species are simply mixed and allowed to grow without artificially inducing resident prophages. It is noteworthy that most species carrying identical cf-PICIs coexist harmoniously, as they are part of the gut microbiota. To explore this possibility, we mixed the *K. pneumoniae* JP24853 strain, carrying the helper phage P1, along with the KpCIDSM30104 island, with the *E. coli* strain JP24850 lysogenic for phage HK022 as the tail provider for KpCIDSM30104 transfer. Additionally, we included the *E. coli* strain JP24888, which carries a non-transmissible plasmid pBAD18-*kmR*, as the recipient in these experiments. Our rationale was that during their growth, the *K. pneumoniae* JP24853 strain would release some tail-less KpCIDSM30104 islands at population level, which would interact with tails released from the *E. coli* HK022 lysogen to generate infective particles that would be transduced into the *E. coli* JP24888 strain. In support of our hypothesis, this was indeed the case, and we observed transfer of the island in this natural scenario (Fig. 4D). To confirm the proposed model of transference, we repeated these experiments, this time using the *E. coli* strain lysogenic for phage HK106 as the tail donor. In this experiment, no transfer of the island was observed (Fig. 4D).

### Packaged cf-PICIs in gut microbiota

Taking into account the previous experiments, along with the fact that EcCIGN02175 was present in different species colonizing the same niche, the human gut, we were prompted to test the possibility that these islands were not only identified in the resident bacteria but also packaged as viral particles, as a precursor to inter-generic transfer. Indeed, this was the case: an identical cf-PICI to EcCIGN02175 (Fig. 4E) was reported in a study that identified tens of thousands of viruses from human metagenomes associated with chronic diseases.^19^ Moreover, we identified another 79 cf-PICI elements (Table S3) from different virome studies, confirming that the presence of packaged cf-PICI DNA occurs frequently in nature. Whether these cf-PICI particles are tail-less or chimeric remains to be determined.

### Expanding cf-PICI intra- and inter-species transfer to other bacterial systems

To confirm the relevance of the previous results showing the abundance of cf-PICIs in species present in the gut microbiome, and to expand our previous studies in which we mobilized the KpCIDSM30104 element from *K. pneumoniae* to *E. coli*, we tested whether the cf-PICI present in the *E. coli* strain GN02175 could be mobilized to *Enterobacter hormaechei* Ehh_18,^20^ an species which also is frequently found in the human gut. This strain was selected because it carries 5 prophages, which will be used as potential tail donors, and a cf-PICI (EhCIEhh_18), which is very similar to EcCIGN02175 (Fig. S6), suggesting that this strain could be a good recipient for EcCIGN02175. The *E. coli* GN02175 and *Enterobacter hormaechei* Ehh_18 strains were induced with MC separately, and then the obtained lysates were used to transfer the EcCIGN02175 island into *E. hormaechei* Ehh_18 mutant in EhCIEhh_18 (*E. hormaechei* Ehh_18 ΔEhCIEhh_18). We eliminated the resident cf-PICI to avoid interaction between the different cf-PICI elements. Importantly, we were able to mobilize EcCIGN02175 into *E. hormaechei* Ehh_18 derivative JP25170 (Fig. 5A).

Next, we hypothesized that once in a new species, EcCIGN02175 could be mobilized intra-species after induction of *E. hormaechei* Ehh_18, as previously shown this strain contains prophages that provide compatible tails for EcCIGN02175. To test this, we MC induced *E. hormaechei* Ehh_18 derivative JP25149, which carries EcCIGN02175 and all the resident prophages present in this strain. Next, we tested the transfer of the EcCIGN02175 island to *E. hormaechei* Ehh_18 ΔEhCIEhh_18 (JP25170). We also tested the transfer of the island back to *E. coli*. Importantly, while we obtained extremely high transfer frequencies when the *E. hormaechei* Ehh_18 derivative JP25170 was used as recipient (Fig. 5B), we did not observe any transfer when *E. coli* was used as a recipient (Fig. 5C). However, if the lysate obtained after induction of *E. hormaechei* Ehh_18 carrying EcCIGN02175 was complemented with tails from *E. coli* phage HK106, then the transfer from *E. hormaechei* to *E. coli* occurred (Fig. 5C).

Finally, to generalize our results, we confirmed that the KpCIDSM30104 island, when present in *E. coli* GN02175, can also be mobilized at astonishingly high frequencies to *E. coli* C1a, especially when compatible HK022 tails are present (Fig. 5D). Altogether, these results support the idea that cf-PICIs can be mobilized back and forth between species when compatible tails are available. They also highlight the potential of these new biological entities (the tail-less cf-PICIs) in mobilizing DNA in nature.

### cf-PICI capsids evolve to interact with different tails

KpCIDSM30104 and EcCIGN02175 share high identity in their packaging modules (Fig. 1A). However, they require different tails to complete the formation of infective particles. In trying to identify the molecular basis for this discrepancy, we realized that although both islands carry head-tail adaptor and head-tail connector genes, the proteins encoded by these genes in both islands were substantially different (50% and 83% identity for adaptors and connectors, respectively; Fig. S7). In view of this, we postulated that these proteins were essential to determine tail compatibility between cf-PICI capsids and tails. To test this, we cloned these two genes from EcCIGN02175 and KpCIDSM30104 into an expression vector. We additionally cloned the head-tail adaptor and head-tail connector genes from EcCIEDL933, which encode completely proteins from those encoded by either KpCIDSM30104 (35% and 63% identity for adaptors and connectors, respectively) or EcCIGN02175 (40% and 61% identity for adaptors and connectors, respectively; Fig. S7). Note that EcCIEDL933 exclusively interacted with the tails from HK106 rather than HK022 (Fig. 4B). These constructed plasmids were introduced into the *K. pneumoniae* DSM30104 mutant in the KpCIDSM30104 adaptor and connector genes (JP24930). Then, the strains were induced with both MC and L-arabinose, and the lysates were mixed 1:1 with lysates from phages HK022 and HK106, respectively, as previously performed. When different head-tail adaptor and head-tail connector genes were expressed in JP24930, the cf-PICI showed different abilities to produce infective particles by interacting with different phage tails (Fig. 5E). These results were consistent with the previous ones in the way that the mutant KpCIDSM30104 specifically interacted with the HK022 tails when the adaptor and connector proteins were from the KpCIDSM30104 or EcCIGN02175, or interacted with the HK106 tails in presence of the EcCIEDL933- or EcCIGN02175-encoded proteins (Fig. 3A, 4A and 4B).

Next, to generalize our results, we performed genomic analyses and searched for almost identical cf-PICIs carrying different head-tail adaptor and head-tail connector genes. As shown in Fig. S1 and Fig. S8, there were examples of cf-PICIs carrying identical packaging modules but different head-tail adaptor and head-tail connector genes. These results confirm the evolutionary adaptation of cf-PICIs to engage with different types of phage tails.

### Overall architecture of the tail-less cf-PICI

The ability of cf-PICIs to expand their host repertoire lies in their capacity to generate tail-less small cf-PICI capsids, which uniquely package cf-PICI DNA. Importantly, the packaging operon within cf-PICI originates from phages that use the prototypical *E. coli* phage HK97 packaging system.^12^ Note that phage HK97 has been used for decades as one of the best models to study phage capsid assembly and maturation.^21,22^ In fact, the cf-PICI proteins encoded in the packaging operon exhibit sequence similarity to homologous proteins encoded by phage HK97.^12^ The evolution of cf-PICI proteins to produce exclusively small capsids, and more significantly, their ability to avoid interaction with their phage counterparts – a crucial process for distinguishing these two distinct entities – remain a mystery. To address these pivotal questions, central to the unprecedented mechanism of gene transfer reported here, we endeavored to determine the structure of cf-PICIs using cryo-EM.

To elucidate the very first structure of a cf-PICI, we MC induced *E. coli* strain JP24598, which carries the prototypical cf-PICI, EcCIEDL933, and its helper prophage HK106. To avoid the formation of the phage capsid, we used a HK106 derivative mutant in its capsid gene (mutation *gp05*\*). Induction of this strain only generated infective EcCIEDL933 particles, but not phage particles.^12^ The resultant lysate was precipitated, and its contents resolved using density gradient ultracentrifugation with CsCl. Due to the substantial excess of HK106 tails present in the purified fraction compared to the assembled EcCIEDL933 particles—a challenge likely inherent in preparations of all cf-PICI, we further purified virions by anion-exchange chromatography (Fig. S9A). This method effectively separated free helper phage tails, which exhibit weak binding to the anionic exchange resin, from more strongly bound, fully assembled cf-PICI virions (Fig. S9B). Mass spectrometry analysis of pure EcCIEDL933 reveals that its capsid is composed of the protein product of gene *1786*.^12^ Our findings demonstrate that this protein undergoes N-terminal cleavage, producing a 32.6 kDa mature capsid protein (MCP) that spans residues 93–327 (Fig. 6A). Initial EcCIEDL933 particles integrity was accessed by negative-staining TEM (Fig. 6B) and upon successful inspection, the same particles were frozen on grids, followed by cryo-electron microscopy (cryo-EM) imaging (Fig. S10A) using the collection parameters listed in Table S4. After initial beam-induced motion and CTF-correction, particles were extracted from images and subjected to several rounds of 2D classification (Fig. S10B) until 2D classes with high-resolution details were obtained. This subset of particles was then used to generate the final 3D reconstruction, with icosahedral symmetry applied. The resulting 3D map of the EcCIEDL933 capsid achieved an overall resolution of 3.24 Å (Fig. 6C, Fig. S10C, Table S4), with local resolutions across most of the capsid shell reaching sub-4 Å resolution (Fig. S10D). An atomic model of the asymmetric unit (ASU), consisting of four MCPs in total—three from the hexon and one from the penton—was constructed (Fig. 6D and E). The model statistics are provided in Table S4.

**Figure 6.**
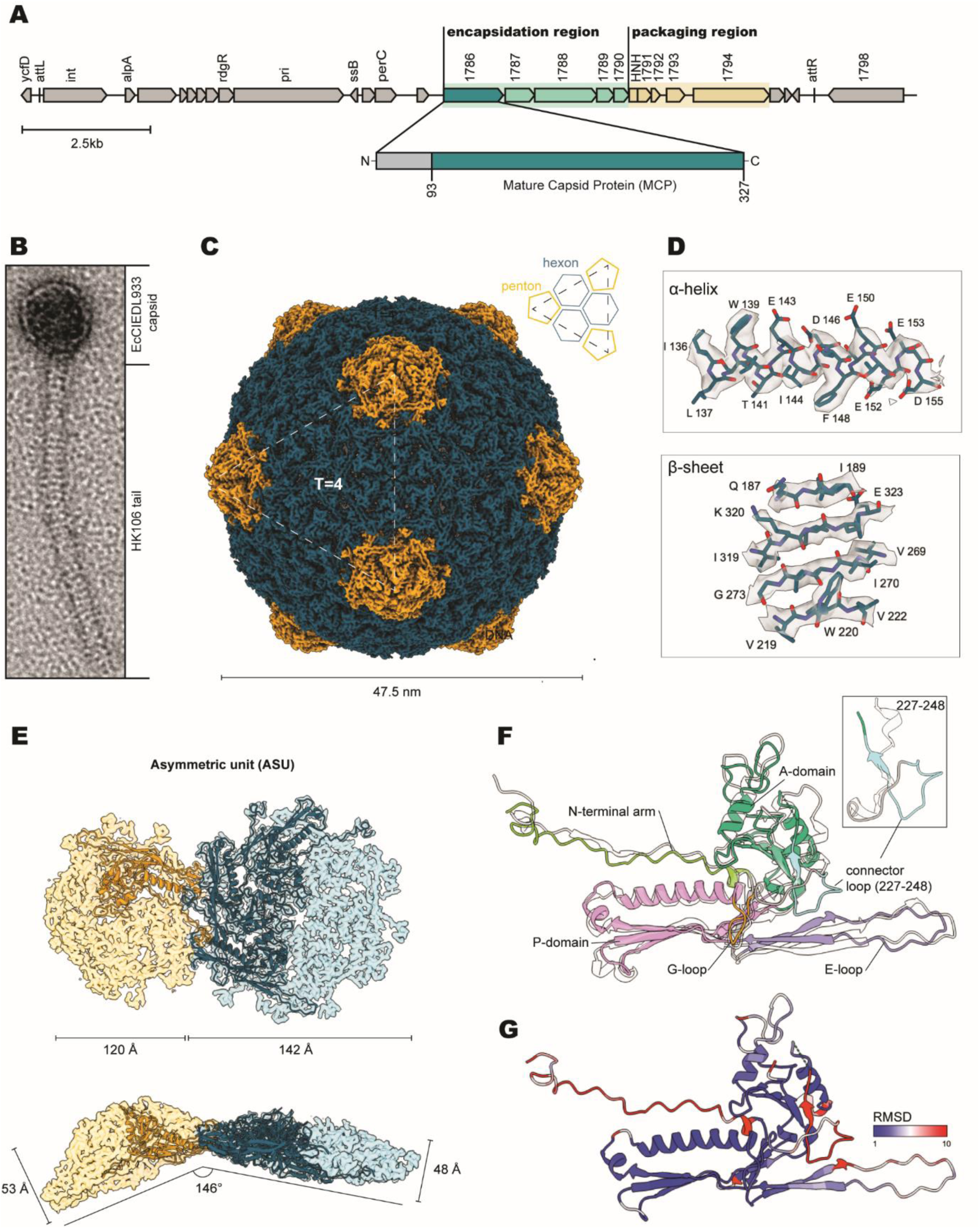
Structure of the EcCIEDL933 capsid. **(A)** Organization of the EcCIEDL933 genome, highlighting the maturation of the protein product of gene *1786,* yielding the 93-327 stretch of the mature capsid protein (MCP). **(B)** Negative stain electron microscopy image of the infective particle of EcCIEDL933, comprised of the island-encoded capsid and the hijacked tail of the co-residing HK106 tail. **(C)** 3.24 Å resolution map of the EcCIEDL933 capsid with diameter of 47.5 nm and T=4 icosahedral symmetry made up of pentameric and hexameric capsomers (pentons and hexons, shown in orange and green, respectively). **(D)** Side chain features of the atomic model (green) and the corresponding map density section (grey) of a representative a-helix (top) and P-sheet (bottom). **(E)** Atomic model of the asymmetric unit and the corresponding density maps of the hexon (green) and penton (orange), showing their respective dimensions and the angle between them. **(F)** Structure of the EcCIEDL933 MCP overlaid on the HK97 MCP (PDH ID: 10HG), showing a nearly perfect match between the structure, with the main divergence coming from the 227-248 connector loop. **(G)** Root mean square deviation (RMSD) between the EcCIEDL933 and HK97 MCP.

The assembled infective particle of cf-PICI EcCIEDL933 exhibits a *Siphoviridae* morphology (Fig. 6B), consisting of the EcCIEDL933 capsid and a long tail hijacked from the co-residing helper HK106 phage. The capsid has a T=4 icosahedral symmetry with a diameter of 47.5 nm (Fig. 6B), similar to that of the classical (non-capsid forming) PICI, SaPI1 with T=4 icosahedral symmetry and 50 nm in diameter (PDB ID: 6C22),^23^ but significantly smaller than the HK97 phage capsid, which has a T=7 icosahedral symmetry and a diameter of 60 nm (PDB ID: 1OHG).^23,24^ Due to its size, the EcCIEDL933 capsid is well-suited to accommodate its 15.5 kbp genome, with DNA forming evenly spaced layers at an approximately 23 Å interlayer distance (Fig. S10D). Larger genomes, such as that of the co-residing HK106 phage at 41.5 kbp (NCBI ID: NC_019768), would exceed the capsid’s internal volume capacity. Thus, the capsid’s diameter and corresponding internal volume size are critical physical barriers that prevent interference of EcCIEDL933 with the propagation of the co-residing HK106 phage.

Topologically, the EcCIEDL933 capsid consists of 240 copies of MCP organized into 12 pentameric capsomers (referred to as pentons, each consisting of five MCPs) and 30 hexameric capsomers (referred to as hexons, each consisting of six MCPs) (Fig. 6B). In nature, one penton of the capsid is replaced by the portal assembly and tail connector, which serve as the DNA packaging and tail attachment site. Our structural data shows that a single MCP of EcCIEDL933 results from proteolytic cleavage, akin to HK97, with both MCPs sharing the N-terminal motif [SxGxxAD], where x denotes any residue (Fig. S11). This suggests a similar maturation mechanism between the EcCIEDL933 and HK97 capsids. Moreover, the MCP of EcCIEDL933 exhibits a clear HK97-like fold, showing a nearly identical structural overlay with the HK97 phage (Fig. 6F), with the root mean square deviation (RMSD) – a quantitative measure of distance between atoms of superimposed structures – nears 1 Å across almost the entire MCP structure (Fig. 6G). The organization of EcCIEDL933 MPC includes the canonical domains characteristic of the HK97-like fold, namely: N-terminal arm (N-arm), axis domain (A-domain), glycine-rich loop (G-loop), periphery domain (P-domain), and extended loop (E-loop) (Fig. 6F). A notable structural difference between EcCIEDL933 and HK97 MCPs is the additional loop, here designated the “connector loop”, formed by residues 227-248 of EcCIEDL933 (Fig. 6F and G), which connects the G-loop and A-domain. Although the functional significance of this loop remains uncertain, a similar motif identified as the S-loop in the atomic model of the bacteriophage P22 MCP (PDB ID: 8I1T)^25^ has been shown to modulate capsid size and symmetry (Fig. S12).^26,27^ Given that the connector loop represents the primary structural divergence between the T=7 capsid of HK97 and the T=4 capsid of EcCIEDL933, it is plausible to propose that this loop functions similarly to the S-loop in P22, granting a change in symmetry, and hence playing a role in establishing the helper phage interference barrier mentioned earlier.

When directly compared, the MCP within the ASU (Fig. S13A) exhibit some expected variation, particularly in the N-arm, A-domain, and E-loop regions (Fig. S13B). Specifically, the penton MCP shows the narrowest conformation, with a 146° angle between its N-arm and E-loop, whereas the hexon MCPs gradually flatten, reaching approximately 180° of the broadest MCP (Fig. S13B). Although these local differences might appear minor, they have a pronounced effect on the topology of the entire capsomers, resulting in a penton that is narrower but taller (120 Å width, 53 Å height) when compared to the wider, shorter hexon (142 Å width, 48 Å height) (Fig. 6E).

### Interactions network between cf-PICI capsomers

In the T=4 icosahedral capsid of EcCIEDL933, two unique intercapsomer interaction sites are observed: between the penton-hexon (Fig. 7A and B) and between hexon-hexon (Fig. 7C and D). These interactions occur on two distinct faces of the MCP: the N-arm face (Fig. 7A and C) and the P-domain face (Fig. 7B and D). The N-arm face interactions are formed along a quasi-two-fold symmetrical axis between anti-parallel N-arms of adjacent capsomers, either between a hexon and a penton (Fig. 7A) or between two hexons (Fig. 7C). In contrast, the P-domain face interactions occur at quasi-three-fold symmetrical axes, involving penton-hexon-hexon (Fig. 7B) or hexon-hexon-hexon assemblies (Fig. 7D). This organization is common to phages sharing the HK97-like fold, such as R4C (PDB ID: 8GTA), TW1 (PDB ID: 5WK1) or RcGTA (PDB ID: 6TSU) phage capsids.^28^ The three-fold interaction sites form an extensive network composed of nine MCP originating from three capsomers: three from the penton and three each from the two neighboring hexons (Fig. 7B), or three from each of three adjacent hexons (Fig. 7D). The three-fold vertex interaction sites can be further divided into two distinct layers: an inner layer, located closest to the center of the vertex, formed by P-domains; and an outer layer, composed of E-loops and N-arms. This organization is visualized in Fig. 7D, where the nine interacting MCPs contributing the P-domains (green), E-loops (orange), and N-arms (purple) are color-coded according to three-fold symmetry. Unique interactions established within this network are shown in Fig. 7E, highlighting a complex array of salt bridges. These include interactions between adjacent P-domains (residues D186 and R345), between E-loops and P-domains (R156 and E183; N148 and Y175; E151 and K219), and between N-arms and E-loops (D99 and Q163) (Fig. 7E). The overall interactions between the inner and the outer layers of the threefold vertex are stabilized by complementary electrostatic potentials, where negatively charged grooves on the surface of P-domains (Fig. 7F) align with positively charged protrusions of the E-loop (Fig. 7G). This groove-protrusion organization is similar to what is observed in HK97, which exhibits a comparable electrostatic complementation pattern (Fig. S14A) (PDB ID: 1OHG).^24^ Notably, no direct physical crosslink between the E-loop and P-domain of neighboring capsomers, such as the interaction observed between residues K169 and N356 in HK97 capsid (Fig. S14B) (PDB ID: 1OHG),^24^ was detected in EcCIEDL933 capsid. This observation is further supported by the fact that no higher molecular weight bands were detected during SDS-PAGE, (Fig. S9A), which are present in HK97 preparations,^29^ indicating that the EcCIEDL933 capsid is stabilized solely by non-covalent interactions.

**Figure 7.**
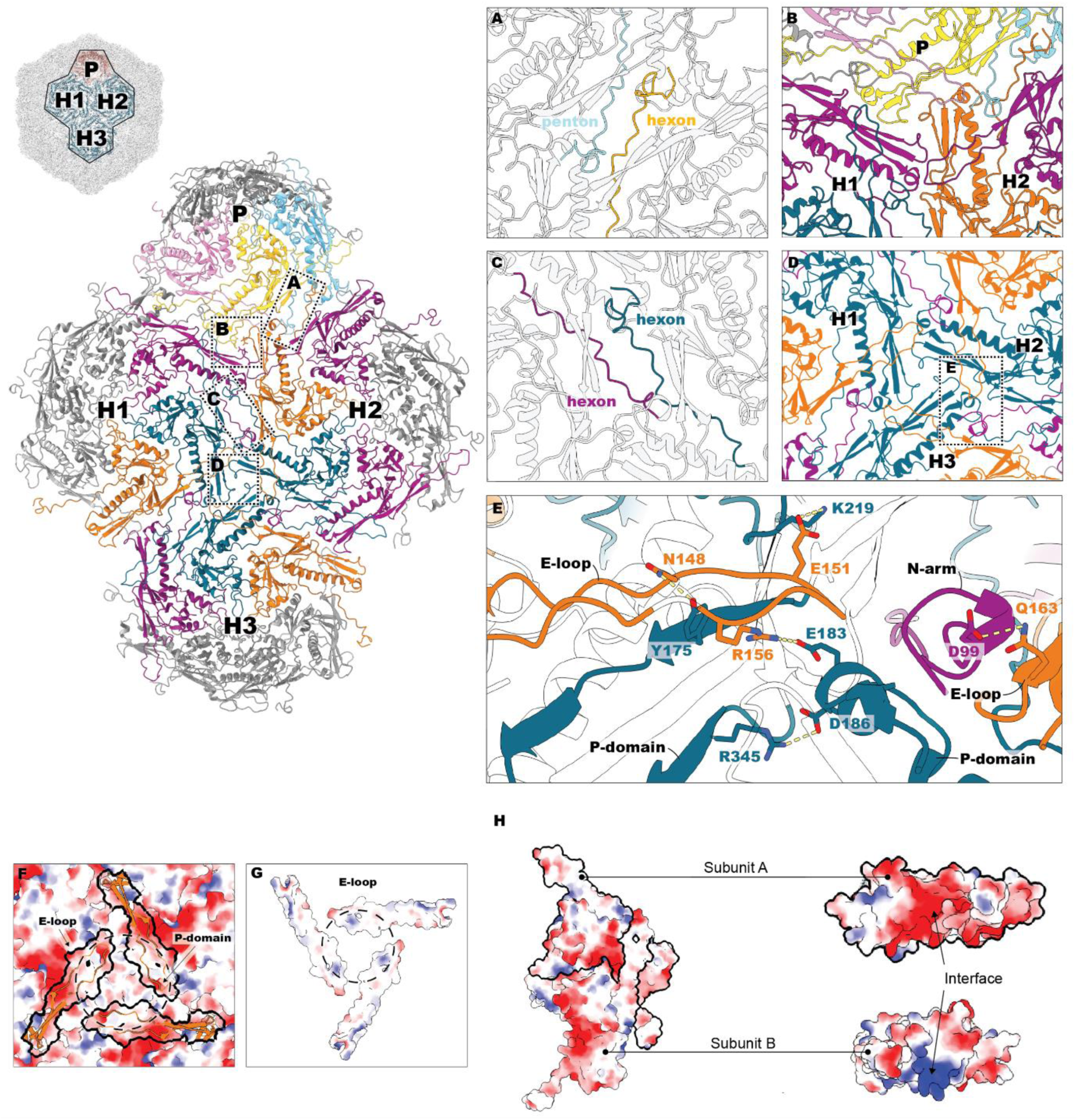
Interaction details between EcCIEDL933 capsomers and within them. To represent intercapsomere interactions, a penton (P) and three hexons (H1-H3) were used used in the analysis. **(A,B)** Penton-hexon interactions are established by two anti-parallel N-arms around a twofold axis **(A)** and between a pentamer and two hexamers around a threefold axis **(B). (C,D)** Similarly, hexon-hexon interactions are formed around the twofold axis **(C)** and threefold axis **(D). (E)** The detailed interaction network of the asymmetric unit of the threefold axis interaction shown in **(E),** salt bridges are marked with yellow dashed lines. **(F)** Electrostatic surface potential of the interaction shown in (D) without the E-Ioops (shown in orange cartoons). (G) The electrostatic surface potential of the bottom side of the E-Ioop. The dashed lines between **(F)** and **(G)** show an electrostatically complementary patch for interaction. **(H)** Electrostatic surface potential of two neighboring MCP within a capsomer showing complementarity between the two interacting faces of subunits A and B.

The fundamental interactions essential for the assembly of any capsid are those between the adjacent MCP subunits. In EcCIEDL933, each MCP subunit exhibits intrinsic polarity, featuring one locally positively charged face and one negatively charged face, enabling stable assembly into capsomers through front-to-back interactions (Fig. 7H). Notably, the presence of a complementary electrostatic interaction between MCP subunits in EcCIEDL933 is a trait clearly present within HK97 as well (Fig. S14C) (PDB ID: 1OHG).^24^ Details of the interactions within capsomers, including specific residues forming salt bridges between MCP in a penton and hexon are shown in Fig. S15A and S15B, respectively.

### Structural validation of the cf-PICI capsid model

As previously mentioned, the interaction network of the EcCIEDL933 MCP is entirely non-covalent, relying on a complex array of hydrogen bonds and salt bridges, with no involvement of additional decoration proteins. Each MCP uniquely interacts with five neighboring MCP (Fig. S16A), with the unique bonds and their respective lengths detailed in Table S5. To identify key residues for structural validation of our model, we compared the interacting residues of EcCIEDL933 (Fig. S16A, Table S5) to those documented in the HK97 phage (Fig. S16B, Table S6) (PDB ID: 1OHG),^24^ providing a comprehensive overview of the interaction networks in both capsids. From these comparisons, we selected residues for mutagenesis that were similarly positioned in both the EcCIEDL933 and HK97 MCP structures and showed a significant effect on the local surface charge. These residues include D256, K289, R294, and R317 in HK97 MCP, and their corresponding residues D257, K290, D295, and E319 in EcCIEDL933 MCP (Fig. 8A). In EcCIEDL933, we also targeted residue R345, which plays a critical role in maintaining the quasi three-fold symmetrical vertex between interacting hexons (Fig. 7E and 8A), along with the deletion of the bulk of the unique connector loop – the knockout spanning residues 232-247 out of the original loop (227-248) (Fig. 6F and 8A). The introduced mutations aimed to reverse the local charge of the selected residues from positive to negative and *vice versa*. The strains carrying WT or the capsid mutant versions of the HK97 prophage, or carrying the helper phage for EcCIEDL933 in presence of the WT or the capsid mutant versions of this island, were MC induced, and the formation of the infective particles analyzed. Overall, the resulting mutants revealed that the EcCIEDL933 capsid is more sensitive to disruption by single-point mutations than the HK97 capsid, highlighting the fragility of its non-covalent inter-subunit interactions. Of the four mutations introduced in HK97, two affected phage reproduction: K289D and D256K, leading to a reduction in infectivity by 0.5 and 3 logs, respectively (Fig. 8B). In contrast, all four corresponding mutations in EcCIEDL933 had a detrimental impact on cf-PICI transfer. Specifically, mutations K290D and E319K completely abolished detectable transfer, while D295K and D257K reduced transfer by 2 and 4 logs, respectively. Furthermore, the additional EcCIEDL933 mutants, including R345D and the deletion of the connector loop, were also found to abolish EcCIEDL933 transfer, underscoring the critical roles of these elements in forming and/or maintaining a functional EcCIEDL933 capsid. This analysis validates the biological relevance of our atomic model of the EcCIEDL933 capsid and emphasizes the delicate balance of non-covalent interactions required for its structural integrity and infectivity compared to HK97.

**Figure 8.**
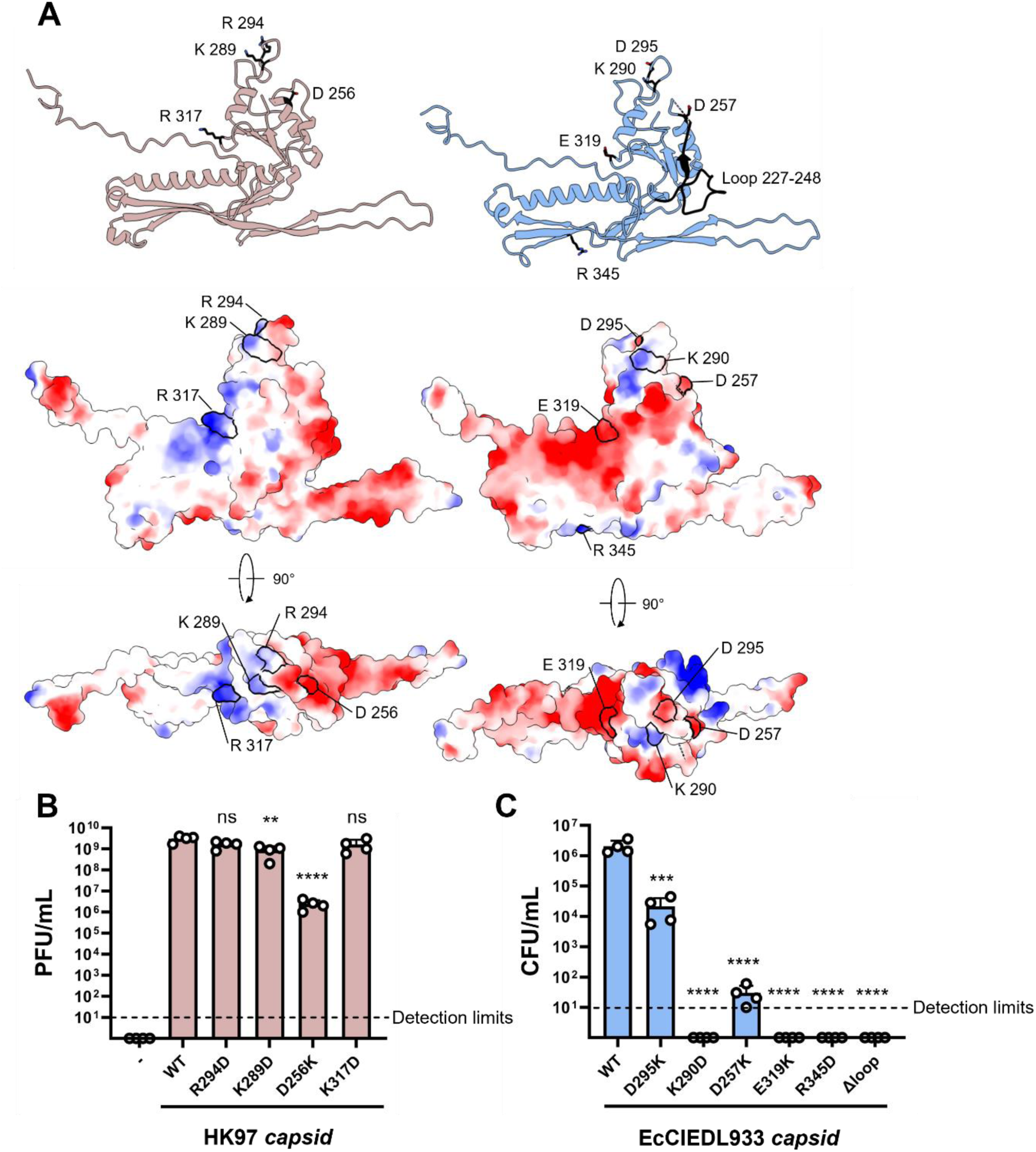
Genetic validation of the structure of the EcCIEDL933 capsid. **(A)** Representation of the residues involved in the formation of salt bridges in EcCIEDL933 (right) and the corresponding residues in HK97, together with the representation of their electrostatic potential. **(B)** HK97 capsid mutagenesis affected phage capsid formation. The strain carrying an HK106 prophage mutant in its capsid gene was complemented with the pBAD18 empty vector (-) or different versions of the HK97 capsid gene. Samples were induced with 2 pg/mL MC and 0.02% arabinose. The resulting filtered lysates were tested for phage titration in *E. coli* strain 594 expressing the WT HK97 capsid gene. Values are presented as means for the plaque-forming units (PFU) per milliliter of lysates. Error bars indicate the standard deviation. A one-way ANOVA with Dunnett’s multiple comparisons test was used to compare the data between samples with the WT HK97 capsid and the other samples after Iog10 transformation, ns: P > 0.05. **: P ≤ 0.01. ****: P ≤ 0.0001. n = 4 independent samples. **(C)** EcCIEDL933 mutagenesis abolished EcCIEDL933 transfer. Derivative strains lysogenic for HK106 and carrying different versions of the EcCIEDL933 capsid gene were MC-induced, and the transfer of the WT and mutant islands was analyzed. Values are presented as means for the colony-forming units (CFU) per milliliter of lysates. Error bars indicate the standard deviation. A one-way ANOVA with Dunnett’s multiple comparisons test was used to compare the data between samples with the WT EcCIEDL933 capsid and the other samples after Iog10 transformation. **** P ≤ 0.0001. n = 4 independent samples.

## DISCUSSION

Since the discovery of the cf-PICIs, a recurrent question has arisen in our minds: what are the evolutionary advantages that the cf-PICIs obtain, compared to other satellites, by encoding a specific packaging module composed of 9 genes in their small genomes? So far, all satellites described apart from cf-PICIs, no more than 1 or 2 genes are encoded in these elements to promote the formation of the small capsids and the preferential packaging of these elements in these capsids. Why do these elements use an important part of their genomes to encode the packaging module instead of using this precious space to encode auxiliary genes that could provide benefits to their host to promote their persistence in different environments? Here we have solved this mystery.

With the classical satellites, since both their packaging and the tails required for the formation of the satellite infective particles were depending on the helper phage, the host tropism of these elements was already defined after induction of the satellites. The situation with the cf-PICIs is completely different. First, they produce packaged cf-PICIs even in the absence of an inducing phage. Since most of the clinical strains are poly-lysogenic, this strategy ensures that they will be released after prophage induction or phage infection. Second, since the cf-PICIs capsids can interact with phage tails from different species, this strategy allows these elements an extremely intra- and inter-species transfer. Note that bacteria have multiple barriers to transduction, including the ability they have to change the phage receptor,^30^ or the possibility to produce barriers to infection (such as capsule or biofilms).^31^ Since the cf-PICIs can interact with different tails, from different species, with different specificities for the surface receptors, this strategy allows them to bypass most of the barriers that limit their dissemination in nature. Third, in addition to their promiscuity in terms of host range, the frequencies of transfer observed with these elements exceed those previously seen with other satellites. This is important because, as demonstrated here, the transfer of these elements occurs in the absence of prophage induction in bacterial populations formed by different bacterial species. It is likely that all these exclusive characteristics have allowed these elements to be the most prevalent satellites in nature described so far.^2^

The phage tail serves as an intricate nano-machinery, specifically designed to identify bacterial host cells, breach the cell wall and/or membrane, and inject the phage genome into the host cytosol, initiating the production of new viral particles.^15^ Positioned at the distal end of the tail, the tail fibers (or spikes) facilitate the binding of the phage to specific receptors on the bacterial host surface, such as lipopolysaccharide (LPS), transmembrane proteins, teichoic acids, or even cellular organelles (e.g., pili or flagella).^32,33^ These tail fibers (or spikes) predominantly dictate host specificity (or range) and influence the process of phage infection.^15^ With a diverse array of phage tail fibers (or spikes),^34^ phages effectively recognize and adhere to a wide spectrum of bacterial hosts. Notably, owing to their capacity to lyse bacterial cells, phages have emerged as potential therapeutic alternatives to antibiotics. However, their therapeutic potential is constrained by the fact that most phages infect only a limited range of strains due to the specific interaction between phage tail fibers and host receptors. Moreover, bacteria can swiftly develop resistance by reducing the expression of the phage receptors, or by blocking or masking these receptors through spontaneous mutation or phenotypic variation.^35^ To overcome these limitations, there is growing interest in engineering characterized phages (primarily by manipulating tail fibers or spikes) to expand or reprogram their host range,^36^ obviating the necessity for isolating new phages and adjusting phage cocktail compositions. Remarkably, cf-PICIs have effectively addressed all the aforementioned issues by generating cf-PICI capsids capable of interacting with various phage tails, thereby infecting different species and strains. Once again, our findings underscore PICIs as intriguing subcellular entities that have evolved unprecedented strategies to facilitate their dissemination in nature.

In the satellite world, helpers have been classically defined as those phages that are able to induce the satellites and provide the components required for their packaging and transfer. In this report, we expand this idea by demonstrating that the induction, the packaging, and the formation of the infective particles are three processes that do not necessarily depend on the same helper phage. Thus, while in the original report describing the cf-PICIs we utilized helper phages that induced the island and produced the tails required for the formation of the infective particles,^12^ the results presented here demonstrate that a more common scenario seems to be that the helper phages induce the island but do not provide the correct tails for the completion of the particle. To that end, these islands require an additional phage, which will broaden the host specificity, depending on the tails. This is a new scenario that highlights the intricate and fascinating biology of the cf-PICIs.

In this study, we present the first detailed structural analysis of a capsid-forming pathogenicity island (cf-PICI), specifically EcCIEDL933. Despite its low sequence homology to the well-studied phage HK97, EcCIEDL933 assembles a nearly identical MCP, sharing many structural characteristics, which underscores their evolutionary relationship. The assembly of the EcCIEDL933 capsid involves a proteolytic maturation of the MCP, akin to the maturation process observed in HK97, further suggesting that cf-PICIs retain fundamental phage-derived assembly mechanisms.

A key evolutionary adaptation observed in EcCIEDL933 is its exclusive formation of a smaller capsid with T=4 icosahedral symmetry, in contrast to the larger T=7 capsid of its helper phage, HK106.^12^ In the absence of other factors that can also been involved in the specific packaging of the cf-PICI DNA into the small sized cf-PICI capsids (i.e. terminase proteins), this size reduction is a critical adaptation that prevents the accidental incorporation of the helper phage genome, allowing EcCIEDL933 to selectively package its own DNA.^37,12^ This phenomenon is not unique to cf-PICIs; a similar strategy is observed in non-capsid-forming PICIs like SaPI1, where capsid morphogenesis proteins CpmA and CpmB, encoded within the island induce a T=7 to T=4 capsid size reduction in the co-residing 80α phage (PDB ID: 6C21 and PDB ID:6C22).^23^ Similar strategies have been also observed in other satellites, such P4^38^ or PLE^39^.

Structurally, the EcCIEDL933 capsid is notable for its lack of additional decorating proteins, with the MCP alone forming the T=4 symmetry. While pinpointing the exact structural modifications driving the smaller capsid size in EcCIEDL933 is challenging, a unique connector loop between residues 227-248 differentiates its MCP from HK97. This loop may function similarly to the S-loop in bacteriophage P22, altering capsid size and symmetry, suggesting a homologous evolutionary adaptation for optimal packaging efficiency. These observations underscore the broader evolutionary trend among all PICIs to modify capsid for selective packaging and efficient gene transfer. One could also envision this as a resource-minimisation strategy that could offer long-term benefits, particularly under stress conditions when resources are sparse, that is because generating a T=7 capsid requires 420 MCP and a T=4 capsid requires only 240.

On the molecular level, the evolution of cf-PICIs like EcCIEDL933 is marked by their ability to exploit the structural versatility of the HK97 fold—a widely conserved organization regarded as “nature’s favorite building block” due to its capacity to form secure, shell-like entities of various sizes and symmetries.^40,22^ This includes not only phage capsids (PDB ID: 1OHG),^24^ but also bacterial and archaeal encapsulins (PDB ID: 3DKT and 2E0Z),^41,42^ and now also cf-PICI. By forming smaller, tail-less capsids, cf-PICIs have adapted this fold to create a unique packaging system that expands their host range and facilitates efficient gene transfer. This structural versatility is key to cf-PICIs evolution, enabling them to maintain distinct identities separate from their helper phages, and avoid detrimental interactions that could hinder their propagation.

Unlike HK97, which uses covalent crosslinks between MCP subunits to stabilize its capsid, the EcCIEDL933 capsid relies on a complex network of non-covalent interactions for stability while simultaneously maintaining the conserved fold needed for attachment of the tail. This highlights the complexity of structural trade-offs cf-PICIs must navigate, balancing mutations that enhance capsid stability while ensuring compatibility with the helper phage tail machinery. At the same time, cf-PICIs must remain distinct enough to avoid interference with the helper phage’s propagation. Overall, our structural data reveal the sophistication of cf-PICI as mobile genetic elements, shedding light on their evolutionary trajectory as specialized molecular pirates that play a significant role in shaping bacterial evolution and pathogenicity.

The formation of hybrid viral particles to cross species barriers is not exclusive to cf-PICIs. Recently, it has been shown that after the infection of human lung cells with influenza A and respiratory syncytial viruses, hybrid viral particles carrying the structural proteins and genomes of both parental viruses were formed.^43^ Remarkably, these filamentous structures were infectious and expanded the cellular tropism of influenza A virus. This suggests that by combining a diverse repertoire of proteins capable of interacting with eukaryotic cells, these previously unknown hybrid structures may have the potential to expand the tropism (and potentially the host range) of the viruses they carry. However, whether these structures form during natural infections and play a relevant role in nature remains to be determined.

In summary, our findings uncover a new subcellular entity that packages specific DNA for delivery into initially non-infective particles. This ingenious strategy is pivotal, enabling these entities to complete the formation of infective particles by attaching tails from phages of different species. Ultimately, this packaged DNA can be mobilized across multiple species, representing an exceptionally powerful system crucial for the evolution of bacterial pathogens.

### Limitations of the study

While cf-PICIs are widespread in nature, all sharing common structural components required to form the small-sized cf-PICI capsids, our study focused on cf-PICIs from *Proteobacteria*, mainly from *E. coli* and *K. pneumoniae*. Moreover, while the formation of the small-sized cf-PICI capsids is an essential process in the formation of this novel biological entity, how these particles specifically interact with the cf-PICI-encoded portal and not with the phage-encoded one remains to be determined. Another interesting area for future research involves understanding the promiscuity of some cf-PICIs in hijacking tails from different phages, while others seem to be less promiscuous. Here, not only the molecular basis of this process but also the ecological consequences of these different strategies require further investigation to fully understand the impact of this novel mechanism of gene transfer in nature.

## ACKNOWLEDGEMENTS

We would like to acknowledge Avinash Shenoy, Thomas Clarke, Patrice Nordmann, Vincent Perreten, Pilar Domingo-Calap, and Olaya Rendueles-Garcia for sharing their strains, phages and/ or plasmids. We would like to acknowledge Aravindan Illangovan for insights with the capsid model building, Joe Barritt for sharing his expertise of cryo-EM of phage capsids, Paul Simpson for the technical support and Ambre Bexter for various insights on the first draft of the manuscript. Electron microscopy data acquisition was performed at the Centre of Structural Biology, Imperial College London.

This work was supported by grants MR/X020223/1, MR/M003876/1, MR/V000772/1 and MR/S00940X/1 from the Medical Research Council (UK), BB/V002376/1 and BB/V009583/1 from the Biotechnology and Biological Sciences Research Council (BBSRC, UK), EP/X026671/1 from the Engineering and Physical Sciences Research Council (EPSRC, UK), and ERC-2023-SyG Project 101118890 – TalkingPhages to J.R.P, and 215164/Z/18/Z/WT to T.R.D.C.

## AUTHOR CONTRIBUTIONS

L.H., J.B.P., T.R.D.C. and J.R.P. conceived the study; L.H., J.B.P., L.M.-R., C.H.S.A and A.F.-S. conducted the experiments; L.H., J.B.P., L.M.-R., C.H.S.A, A.F.-S., T.R.D.C. and J.R.P. analyzed the data; T.R.D.C. and J.R.P. wrote the manuscript, with inputs from L.H. and J.B.P.

## STAR Methods RESOURCE AVAILABILITY

### Lead contact

Further information and requests for resources and reagents should be directed to and will be fulfilled by the Lead Contact, José R Penadés.

### Materials availability

Strains, phages and plasmids generated in this study are available upon request and without restrictions from the lead contact upon request.

### Data and code availability

All data reported in this paper will be shared by the lead contact upon request. This paper does not report original code. Any additional information required to reanalyze the data reported in this paper is available from the lead contact upon request.

## EXPERIMENTAL MODEL AND SUBJECT DETAILS

### Bacterial strains and growth conditions

Phages and bacterial strains used in this study are listed in table S7. *E. coli*, *K. pneumoniae, E. hormaechei*, *Enterobacter cloacae, Citrobacter koseri, Citrobacter freundii* and *Salmonella enterica* strains were grown at 37°C or 30°C on Luria-Bertani (LB) agar or in LB broth with shaking (120 r.p.m.). When appropriate, Ampicillin (100 μg/mL), Streptomycin (100 μg/mL) Kanamycin (30 μg/mL), Tetracycline (20 μg/mL), or Chloramphenicol (20 μg/mL) were added. For auxotrophic *E. coli* strain S17 MFD*pir*, 0.3 mM DAP (2,6-Diaminopimelic acid, Sigma-Aldrich) was added when required.

## METHOD DETAILS

### Plasmids and oligonucleotides

The plasmids and oligonucleotides used in this study are listed in S8. The plasmid pWRG99-*kmR* for mutagenesis was constructed using Gibson assembly,^44^ using the oligonucleotides referenced in Table S8. Additional plasmids were generated by cloning PCR products (amplified with oligonucleotides from Table S8) into the pBAD18 or pKNG101 vector via restriction-ligation. All plasmids were verified by Sanger sequencing in Eurofins Genomics.

### Phage and PICI induction

Overnight cultures of lysogenic strains were diluted in 1:50 in LB and then sub-cultured until OD_600_ = 0.15. Mitomycin C was added at the final concentration of 2 μg/mL. The sub-cultures were supplemented with 0.02% L-arabinose only if the strains were containing pBAD18 derivative plasmids to express corresponding proteins. Sub-cultures were subsequently incubated at 80 r.p.m. at 30 °C for 4 hours followed by incubating at 25 °C without shaking overnight. Following induction, lysates were collected, centrifuged and then filtered using sterile 0.2 μm filters. The number of phage particles in the lysates was quantified.

### Capsid precipitation and DNA extraction

Following phage and PICI induction, 35 mL lysates were filtered through 0.2 μm sterile filters and treated with DNase and RNase. Capsids were precipitated using PEG 8000 (Polyethylene Glycol 8000, Sigma-Aldrich) after centrifugation at 11,000 g, and then resuspended in 200 μL Phage Buffer (PHB, 50 mM Tris pH 8.0, 1 mM NaCl, 1mM MgSO_4_, 4mM CaCl_2_). The precipitated lysates were lysed in 200 μL Lysis Mix (9.5 µL SDS 20%, 4.5 µL 20 mg/mL proteinase K, 90 µL H_2_O) at 55 °C for 1 hour. DNA was extracted with Phenol:Chloroform:Isoamyl Alcohol method and resuspended in 50 µL TE buffer (pH 8.0). 1 µL DNA was used for electrophoresis analysis, 0.4 µL DNA was used for Southern blotting analysis.

### Southern blotting

After phage and PICI induction, 1.5 mL samples were taken at defined time points and pelleted. DNA was extracted using the bacterial genomic DNA kit (GenElute Bacterial Genomic DNA Kit, Sigma-Aldrich). 5 µL genomic DNA was separated in a 0.7% agarose gel at 25V overnight, treated with 0.25 M HCl and 0.4 M NaOH, and transferred to a nylon membrane (Nylon Membranes positively charged, Roche) by inverse capillarity. The membrane was hybridized with a DIG-labeled probe (Digoxigenin-11-dUTP, alkali-stable; Roche), washed, and detected using an anti-DIG antibody (Anti-Digoxigenin-AP; Roche) and CSPD (Roche). Primers used for DIG-labeled probes are listed in Table S8.

### PICI transduction with exogenous tails

50 μL PICI lysate and 50 μL phage WT or mutant lysate were mixed and incubated at 25 °C for 30 mins, followed by a 10-fold serial dilution in PHB. 100 μL of diluted lysate was added to 1 mL recipient strain (OD_600_ = 1.4 for *E. coli*, OD_600_ = 0.6 for *K. pneumoniae, E. hormaechei, Enterobacter cloacae, Citrobacter koseri, Citrobacter freundii* and *Salmonella enterica*) supplemented with 4.4 mM CaCl_2_, following by incubation at 37 °C for 30 mins. 3 mL LTA (LB top agar) were then added to the samples and plated on the LBA plate with selective antibiotics. The plates were incubated in 37 °C for 48 hours. The number of forming colonies was counted and represented as CFU/mL of PICI lysate.

To maximize the detection of PICI transfer and measure the transfer of the same PICI with different WT *E. coli* phages comparably, these *E. coli* phages (HK022 and HK106) were quantified to PFU/mL = 10^10^. The same procedure was followed as above. Colony counts were represented as CFU/mL of PICI lysate.

### PICI transduction with endogenous tails

A 10-fold serial dilution of PICI lysate using PHB was performed. 100 μL of diluted lysate was added to 1 mL recipient strain (OD_600_ = 1.4 for *E. coli*, and OD_600_ = 0.6 for *E. hormaechei*) supplemented with 4.4 mM CaCl_2_, and incubated at 37 °C for 30 mins. 3 mL LTA were then added to the sample and plated on the LBA plate with selective antibiotics. The plates were incubated in 37 °C for 48 hours. The number of forming colonies was counted and represented as CFU/mL of PICI lysate.

### Phage titration

Recipient strains were sub-cultured until OD_600_ = 0.34. Bacterial lawns were prepared by mixing 300 μL of cells with phage top agar (PTA, 25 g of Nutrient Broth No. 2, Oxoid; 4g agar) and poured onto square plates with phage bottom agar (PBA; 25 g of Nutrient Broth No. 2, Oxoid; 7g agar) supplemented with 10 mM CaCl₂. Serially diluted phage lysates were spotted on the agar. Plates were incubated at 37°C for 24 hours. Plaques were counted and quantified as plaque-forming units (PFU/mL).

For HK97 capsid mutagenesis verification experiments, PBA was supplemented with 0.02% L-arabinose to induce the expression of WT HK97 capsid in recipient strains.

### DNA methods

For phage and PICI scarless deletion, an allelic replacement method was performed. Empty attachment sites of bacteria (*attB*) along with their flanking regions were amplified by PCR using primers listed in Table S8 and then cloned to the allelic-exchange vector pKNG101.^45^ The constructed pKNG101 derivatives were trans-conjugated from the auxotrophic doner strain S17 MFD*pir* to the recipient strain.^46^ Overnight cultures of donor and recipient strains were washed with PBS, mixed at a 1:1 ratio, and spotted on LBA plates with DAP for conjugation at 37°C. Transconjugants were selected on LBA plates with 200 μg/mL streptomycin. Sucrose-sensitive transconjugants were cultured in LB without NaCl at 30 °C overnight. The second recombination event was selected on NaCL-free LBA plates complemented with 20% sucrose at 37°C. Mutants obtained were verified by PCR and Sanger sequencing in Eurofins Genomics.

The HK022 tail deletion mutant was constructed based on the allelic-exchange vector pKNG101. The flanking regions of the HK022 major tail gene were amplified using primers from Table S8 and cloned into pKNG101. The remaining gene mutagenesis process was performed as above described. Mutants were verified by PCR and Sanger sequencing in Eurofins Genomics.

Gene insertions or deletions in *E. coli* and *K. pneumoniae* were performed using a λ Red recombinase-mediated method.^47^ The kanamycin resistance cassette (*kmR*) or tetracycline resistance cassette (*tetA*) was amplified from pKD4 or pRW224 by PCR using primers listed in Table S8. In brief, PCR products were electroporated into the recipient strains harboring pWRG99 or pWRG99-*kmR*, allowing λ Red recombinase-mediated insertion of antibiotic cassettes into the bacterial genome. To remove *kmR* cassette flanked by flippase recognition target (*FRT*) sites, a thermal sensitive plasmid pCP20 was transformed into the corresponding strain. The strains carrying pCP20 were cultured at 30 °C overnight with ampicillin to permit FLP recombination. Overnight culture was diluted in fresh ampicillin-free LB at a 1:50 ratio and then grown at 42 °C for 5 h to promote plasmid loss. Kanamycin-sensitive colonies were selected and verified by PCR and Sanger sequencing in Eurofins Genomics.

For deletion of β-Lactamase gene *shv-1* from DSM30104, the *tetA* cassette was first inserted to replace *shv-1* using pWRG99-*kmR*. The *tetA* cassette was then replaced by a *kmR* cassette flanked by *FRT* sites, followed by the deletion of *kmR* using pCP20. Mutants were verified by PCR and Sanger sequencing in Eurofins Genomics.

Site-directed scarless mutagenesis of EcCIEDL933 capsid mutants was performed as previously described.^37^ The *kmR* cassette flanked by I-SceI recognition sites was PCR-amplified from pWRG717 using primers listed in Table S8. PCR products were then electroporated into the recipient strains harbouring pWRG99 for λ Red recombinase-mediated insertion. Mutants with inserted cassettes were verified by PCR. After that, a second PCR product containing the desired mutations were electroporated into the strain with cassette. The second recombination event was selected on LBA plates with anhydrotetracycline (AHT)-induced I-SceI meganuclease at 30 °C. Mutants were verified by PCR and Sanger sequencing in Eurofins Genomics.

### Mimicking gut environment to study cf-PICIs transfer

Non-lysogenic *K. pneumoniae* JP24871 carrying KpCIDSM30104 was sub-cultured 1:50 in fresh LB without antibiotics until OD600 = 0.2. The strain was then treated overnight with no treatment (control), lytic phage K68 QB (MOI = 0.01), fresh lysozyme (500 μg/mL), or ampicillin (50 μg/mL). Filtered lysates were tested for transduction in *E. coli* C1a, using exogenous phage HK022. The number of forming colonies was counted and represented as CFU/mL of PICI lysate.

### Co-cultivation and selection of KpCIDSM30104 transfer

Single colony of *E. coli* recipient strain, *E. coli* lysogen strain and *K. pneumoniae* strain (containing KpCIDSM30104 Δ*orf17*::*tetA*) was inoculated in LB with appropriate antibiotics for overnight, respectively. The overnight culture was centrifuged at 4200 rpm and then the supernatant was discarded to eliminate the antibiotics. The pellet was resuspended by LB without antibiotics. This resuspended overnight culture was diluted in 1:50 in LB without antibiotics and then sub-cultured at 120 r.p.m. at 37 °C until OD_600_ = 0.2. The sub-cultures of *E. coli* and *K. pneumoniae* were mixed at the ratio 1:1:1 followed by co-cultured at 80 r.p.m. at 30 °C for 5 hours. After that, 1 mL co-culture was added 3 mL LTA and plated on LBA plated containing both 30 μg/mL kanamycin and 20 μg/mL tetracycline to select the transferred KpCIDSM30104 from *K. pneumoniae* to *E. coli*. Number of forming colonies was counted and represented as colony forming units (CFU/mL of co-culture).

### Identification of cf-PICI candidates in virome samples

We searched for cf-PICIs in metagenome studies, focusing on their common features as previously described.^48,4912^ Specifically, we targeted cf-PICIs with: (i) a size of approximately 10–18 kb; (ii) conserved gene organization, including an integrase, site-specific recombinase, or GIY-YIG endonuclease, along with a replication module, regulation module, and exclusive packaging modules (capsid, protease, portal, adaptor, connector, HNH endonuclease, small terminase, large terminase); and (iii) the absence of lytic genes and a complete tail module.

### Bioinformatics analysis

Prophages in natural isolated were predicted by PHASTEST.^50^ Comparison of phage or cf-PICI sequences was based on BLASTn, BLASTp, and PRALINE alignments.^51^

### Quantification and statistical analysis

Experiments were repeated four times, with sample sizes indicated in the figure legends. Data are presented as mean ± SD. Statistical analyses were performed with GraphPad Prism v.10. One-way ANOVA with Dunnett’s multiple comparisons test or Two-way ANOVA with Sidak’s multiple comparisons test was performed to compare three or more groups. An unpaired t-test was applied to compare two groups. Adjusted P values as: ns: P > 0.05; ∗: P ≤ 0.05; ∗∗: P ≤ 0.01; ∗∗∗: P ≤ 0.001; ∗∗∗∗: P ≤ 0.0001.

### EcCIEDL933 purification

The *E. coli* strain JP24598, which overexpresses *alpA* to promote EcCIEDL933 reproduction,^12^ was grown overnight at 37°C in LB medium supplemented with 100 µg/mL of ampicillin with 180 rpm shaking. On the next day, 1 mL of the overnight culture was added into each of 3x1L of fresh LB supplemented with 100 µg/mL ampicillin and grown at 37°C, 180 rpm until OD600=0.05. 0.02% L-arabinose was added, and growth was continued at 37°C, 120 rpm until OD600=0.1-0.2. Phage was induced by addition of 2 µg/mL MC and incubated for 4 h at 32°C, 80 rpm. The cultures were further incubated overnight at room temperature (RT) for bacterial complete lysis. On the next day, 0.2 mg/mL lysozyme, 1 µg/mL RNAse and 1 µg/mL DNAse I was added to the lysates and incubated for 1h at RT. The cultures were centrifuged 10,000 x G for 20 min at 4°C to remove cell debris, and the supernatant was collected and spun down again at 10,000 x G for 20 min 4°C. The supernatant was transferred into a beaker and 1M NaCl was added, followed by 1h incubation at 4°C with gentle stirring. After the lysate was centrifuged at 10,000 x G for 20 min at 4°C, the supernatant was supplemented with 10% PEG 8,000 and stirred until dissolved. The solution was then incubated without stirring overnight at 4°C. On the next day, the precipitate was collected by centrifugation at 10000 x G for 30 min at 4°C and the remaining liquid was removed by placing the tube upside down for 30 min. All pellets were resuspended by addition of 3 mL phage buffer (50mM Tris, pH=8, 100 mM NaCl, 1 mM MgSO_4_) and gently pipetting up and down. The resulting resuspension was applied on top of a pre-layered 1.3, 1.4 and 1.6 g/cm^3^ CsCl density gradient. After 2h ultracentrifugation at 80,000 x G at 4°C, the visible EcCIEDL933 containing layer was collected and dialysed overnight against dialysis buffer (50mM Tris, pH 8.4, 100 mM NaAc). Following dialysis, the EcCIEDL933 sample was slowly diluted with 50 mL of buffer A (50 mM Tris pH 8.4 10 mM NaAc) and loaded into a 1 mL Q Column XL (Cytvia) pre-equilibrated with the buffer A. The column was extensively washed with buffer A, followed by a gradient elution with buffer B (50 mM Tris pH 8.4 2M NaAc) over 80 mL. Free helper phage tails elute at 300-500 mM NaAc, and the assembled EcCIEDL933 particles at 800 mM to 1M NaAc. The eluted particles were dialysed against buffer A and gently concentrated using a centrifugal concentrator filtered through a 0.2 µm filter and stored at 4°C. The determined EcCIEDL933 titer in the sample was approximately 3x10^10^ particles.

### Grid preparation and cryo-EM data collection and processing

3 µl of freshly prepared EcCIEDL933 sample was applied on glow discharged (70s) holey carbon QuantiFoil R2/2 200 mesh grids (Agar Scientific) using Vitrobot (Thermo) operating at 95% humidity at 4°C and vitrified in liquid ethane. The cryo-EM data was acquired using a Glacios Microscope (Thermo Scientific), operating at 200kV and equipped with Falcon 4 direct electron detector. 4337 movies were recorded using EPU software at a magnification of x100000, corresponding to 1.1 Å/px pixel size,50 e−/Å^2^ electron dose and a defocus range of -0.8 to -2.8 μm. All processing was performed with in CryoSparc v4.4.1 software.^52^ After all movies were motion- and CTF-corrected, the micrographs were curated to select those with ideal ice thickness, CTF estimation and resolution. From a total of 3525 selected micrographs, 200 capsid particles were manually picked and 2D classified to generate templates for automated particle picking in the entire dataset. A total of 4658 particles were automatically picked and extracted after binning to a box size of 1000 px. Low quality particles were removed after several rounds of 2D classification, followed by a final extraction with a larger box size of 1300 px and subsequent downsampling by Fourier cropping to 500 px. Following a final round of 2D classification, 1933 particles were selected to generate an ab initio model that was non-uniform refined to an estimated 3.24 Å resolution when an icosahedral symmetry was imposed. The local resolution of the final map was estimated in CryoSparc and visualised in ChimeraX.^53^ Details of the data collection and 3D refinement are provided in Table S4.

### Atomic model building and refinement

The initial atomic model of the ASU was built using ModelAngelo,^54^ followed by manual correction of chain placements and real-space refinement in Coot.^55^ The resulting model was refined iteratively with Phenix^56^ until Ramachandran outliers and poor rotamers were in the favored regions. The final atomic model was validated with Molprobity^57^ and Phenix^56^ and visualized in ChimeraX.^53^ Contacts between chains were detected using ChimeraX^53^ and PDBe PISA software (https://www.ebi.ac.uk/pdbe/pisa/picite.html). The final capsid map and model was deposited under EMD-51699 and PDB ID 9GYI.

### Negative stain electron microscopy

10μl of purified phage sample was applied onto glow-discharged carbon coated copper grids (300 mesh, Agar Scientific) and incubated for 2 min at room temperature. Grids were washed three times in 10 μl of sterile water, excess was blotted away with filter paper and the grids stained for 1 min with 2% uranyl acetate. Excess was removed with blotting paper and grids were allowed to dry for 5 min. Negative-stain electron microscopy images were acquired using FEI Tecnai 12 Spirit operating at 120 kV equipped with a LaB6 filament and a CCD TVIPS fast frame camera.

