## Supplementary Figures for "Chimeric infective particles expand species boundaries in phage inducible chromosomal island mobilization"

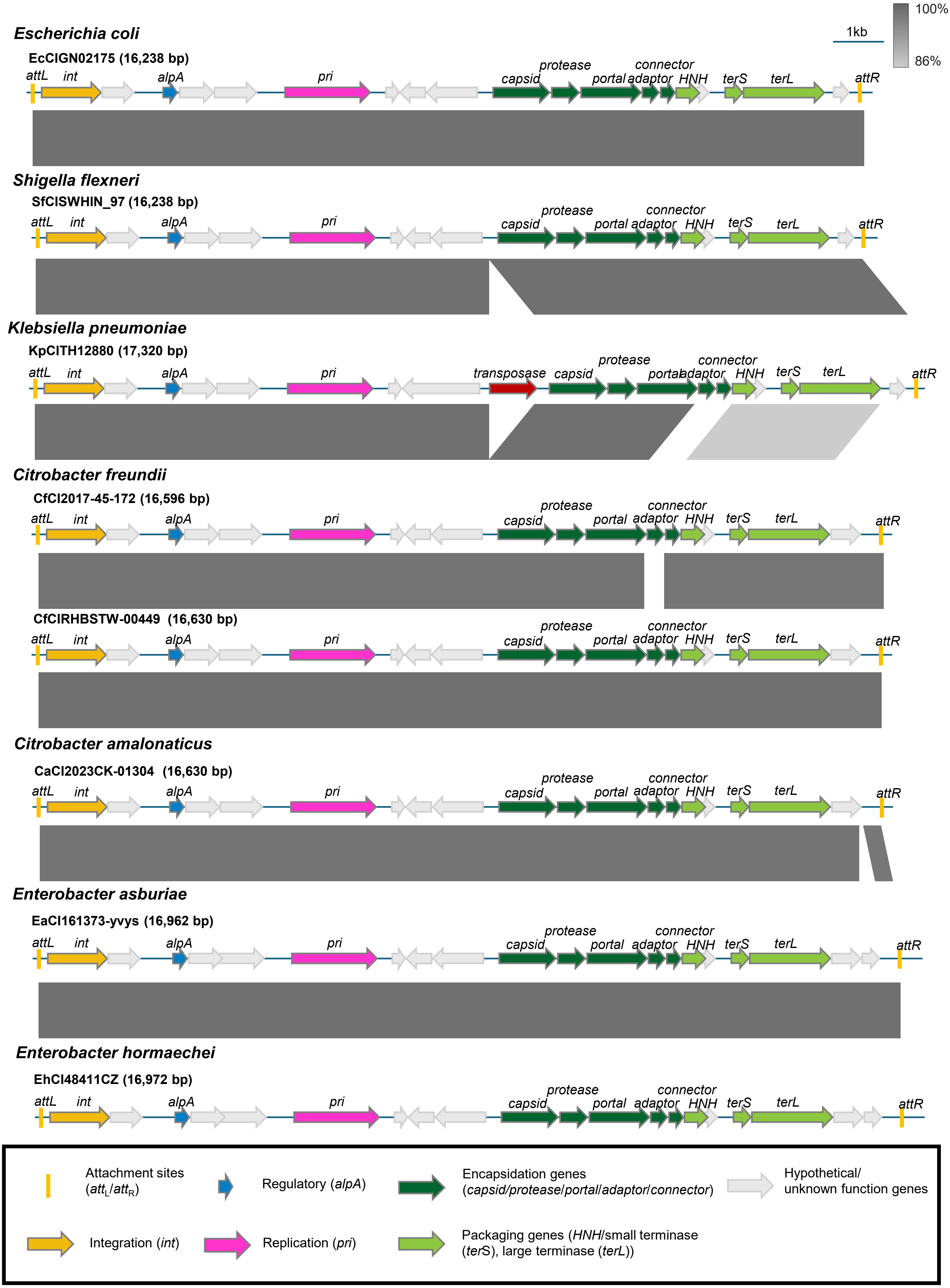


**Figure S1. EcCIGN02175 is present in five different bacterial genera and seven different bacterial species.** Genes are colored according to their sequence and function (see box for more details). Grey arrows indicate hypothetical proteins. Grey scales between cf-PICI sequences indicate that the region share similarity based on BLASTn. See also Table S1.

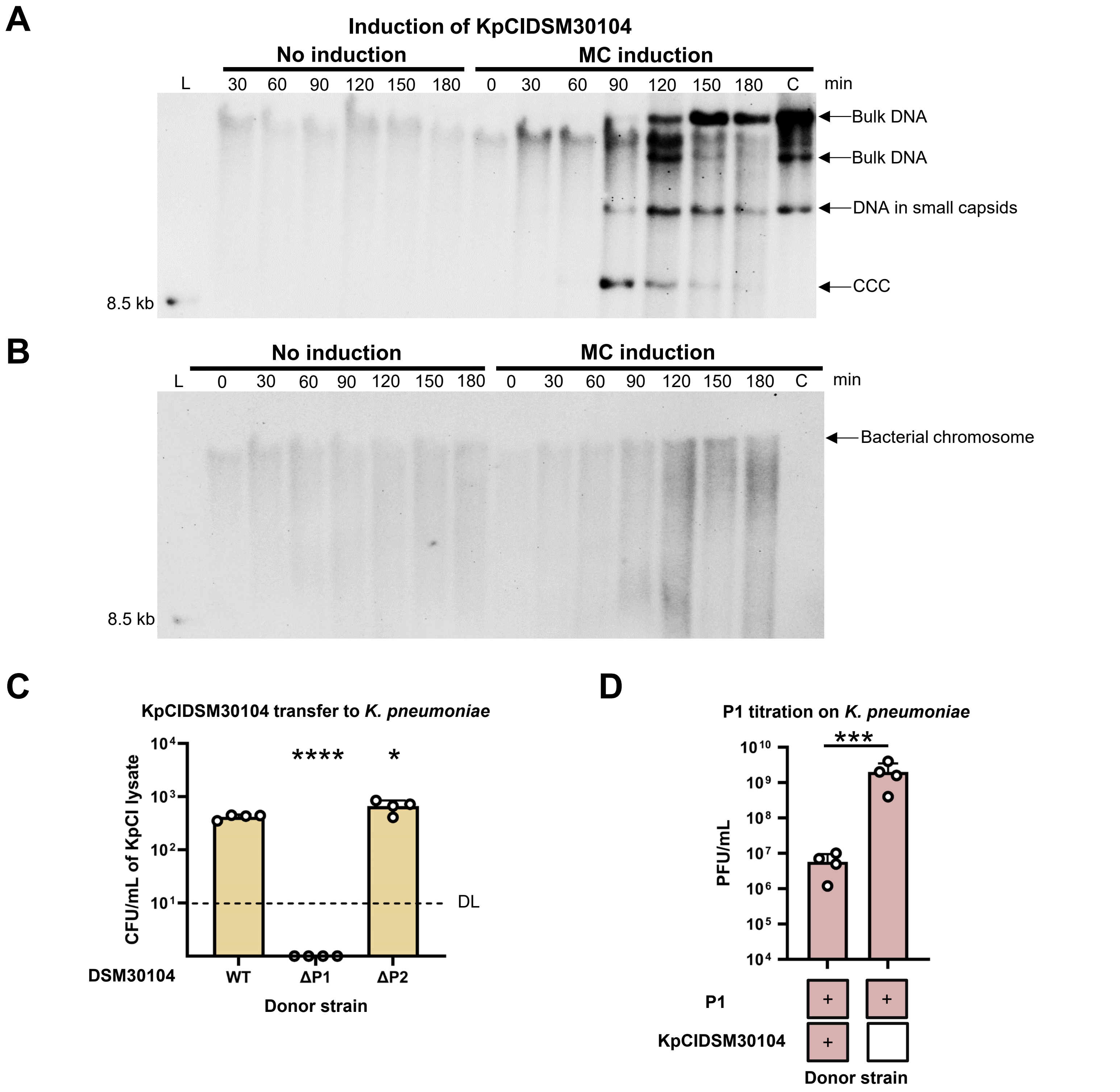


**Figure S2. Phage P1 induces KpCIDSM30104.**

*K. pneumoniae* DSM30104 strains mutant in P2 **(A)** or in P1 **(B)** were MC induced, and samples were taken at the indicated time points for DNA analysis. DNA was separated on a 0.7% agarose gel, followed by Southern blotting analysis using a specific KpCIDSM30104 probe. L: Southern blot molecular marker (DNA molecular weight marker VII; Roche). C: DNA extracted from capsid. CCC: Covalently closed circular.

**(C)** Deletion of ΔP1 abolished intra-species KpCIDSM30104 transfer. The *K. pneumoniae* DSM30104 strains mutant in P1 or P2 were MC induced, and the resulting lysates were tested for KpCIDSM30104 transfer, using *K. pneumoniae* strain JP24460 as the recipient. Values are presented as means of colony-forming units (CFU) per milliliter of cf-PICI donor lysates. Error bars represent the standard deviation. After log10 transformation, a one-way ANOVA was conducted followed by Dunnett's multiple comparisons test to compare the WT sample to other samples. n = 4 independent samples. *: P ≤ 0.05. ****: P ≤ 0.0001. WT: wild-type DSM30104 carrying both prophages P1 and P2. ΔP1: DSM30104 ΔP1. ΔP2: DSM30104 ΔP2. KpCI: KpCIDSM30104.

**(D)** KpCIDSM30104 interferes with P1 reproduction. DSM30104 derivative strains carrying P1, in the presence or absence of KpCIDSM30104, were MC induced, and the P1 titer was quantified. '+' indicates the presence of P1 or KpCIDSM30104 in the donor strains. The recipient strain was *K. pneumoniae* strain JP24460, a derivative of DSM30104 mutant in P1, P2, and KpCIDSM30104. Values are presented as means of plaque-forming units (PFU) per milliliter of lysates. Error bars indicate the standard deviation. A t-test was used to compare the data after log10 transformation. ***: P ≤ 0.001. n = 4.

A

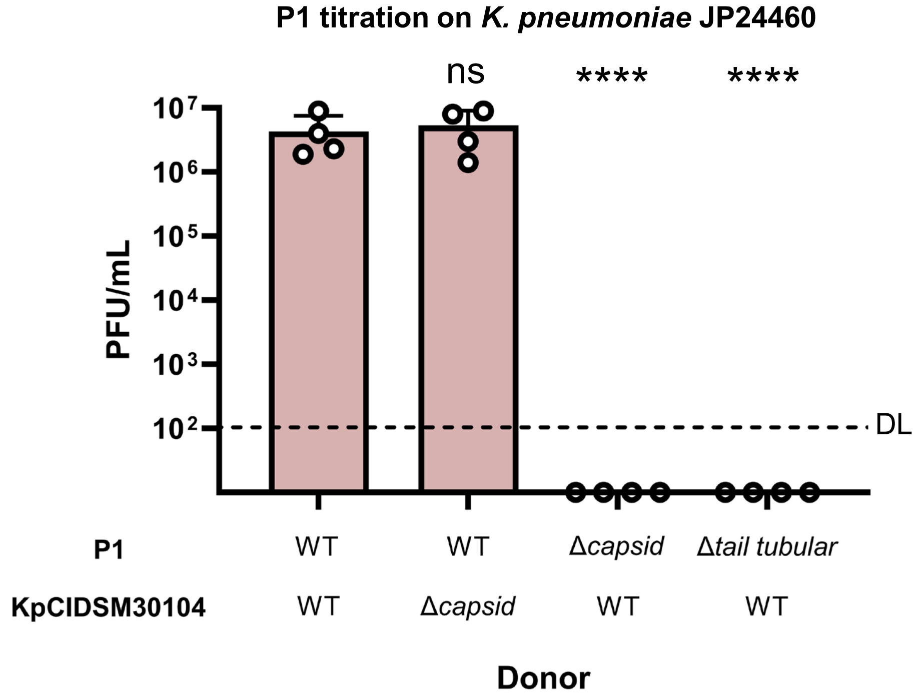


B

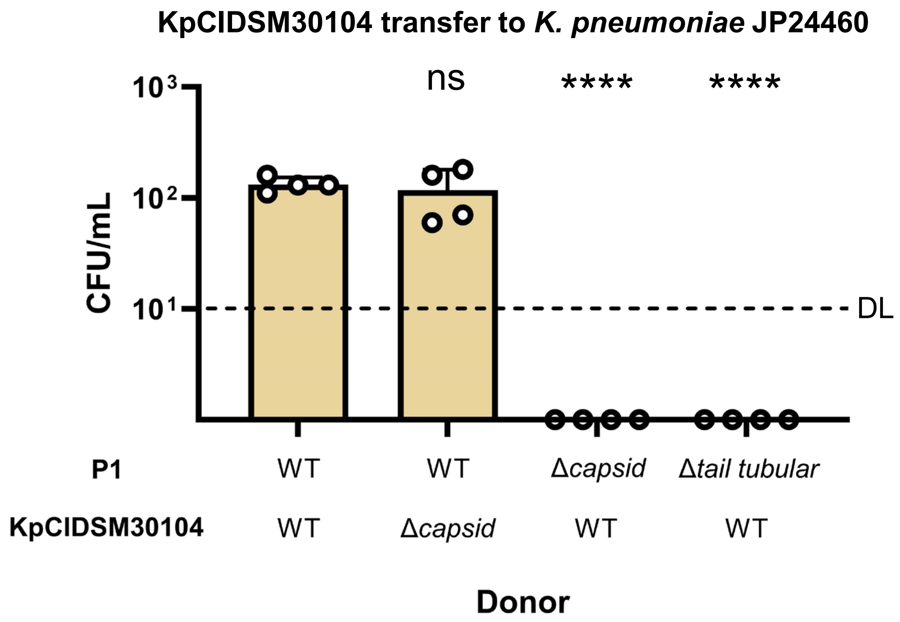


**Figure S3. The low intra-species transfer of KpCIDSM30104 was due to P1-mediated generalized transduction.**

Derivative *K. pneumoniae* DSM30104 strains carrying either WT P1, P1 mutant in the capsid gene, or P1 mutant in the tail protein, and carrying either WT KpCIDSM30104 or its capsid mutant derivative, were MC induced, and the formation of P1 **(A)** or KpCIDSM30104 **(B)** infective particles was measured, using *K. pneumoniae* strain JP24460, a derivative of DSM30104 mutant in P1, P2, and KpCIDSM30104, as the recipient. After log10 transformation, a one-way ANOVA was conducted followed by Dunnett's multiple comparisons test to compare the wild-type DSM30104 sample to other samples. n = 4 independent samples. ns: $P > 0.05$. ****: $P \leq 0.0001$. WT: wild-type element of P1 or KpCIDSM30104 in DSM30104 derivative strains. DL: Detection limits.

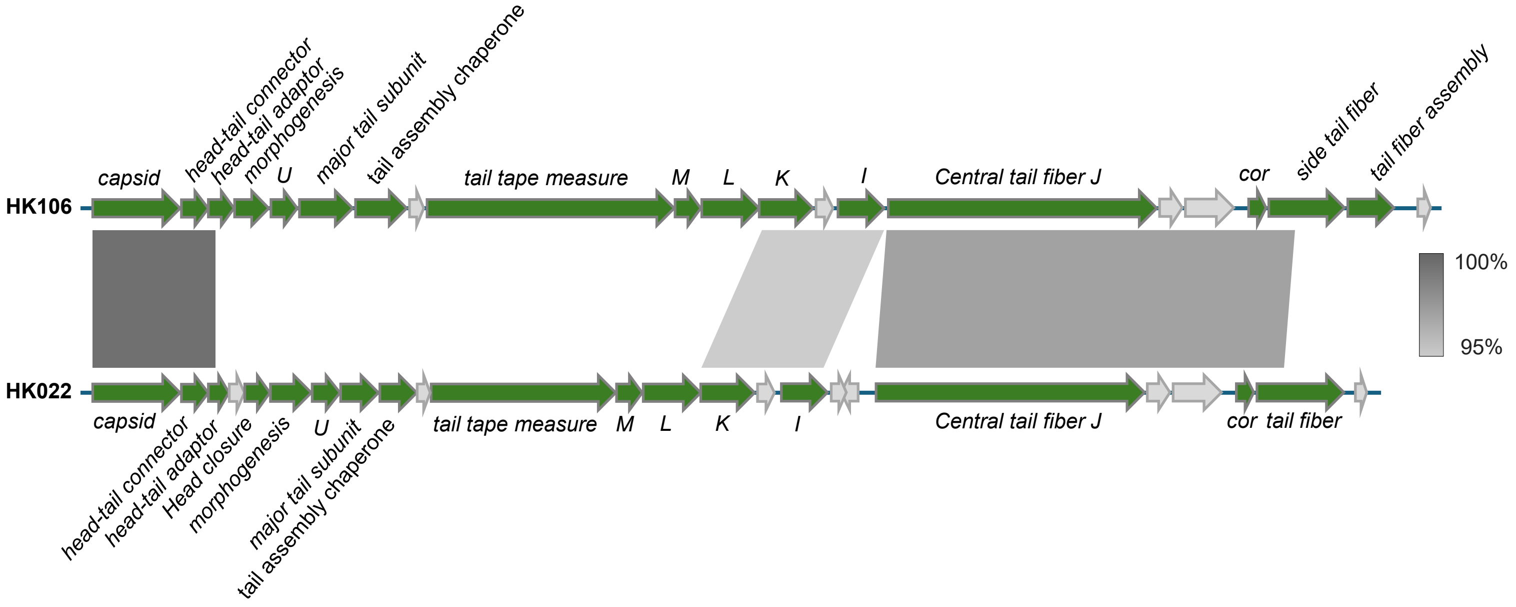


**Figure S4. Comparative maps of the tail region encoded by the HK022 and HK106 prophages.** Green arrows indicate phage structural proteins. Grey arrows indicate hypothetical proteins. Grey scales between phage sequences indicate regions that share similarity based on BLASTn.

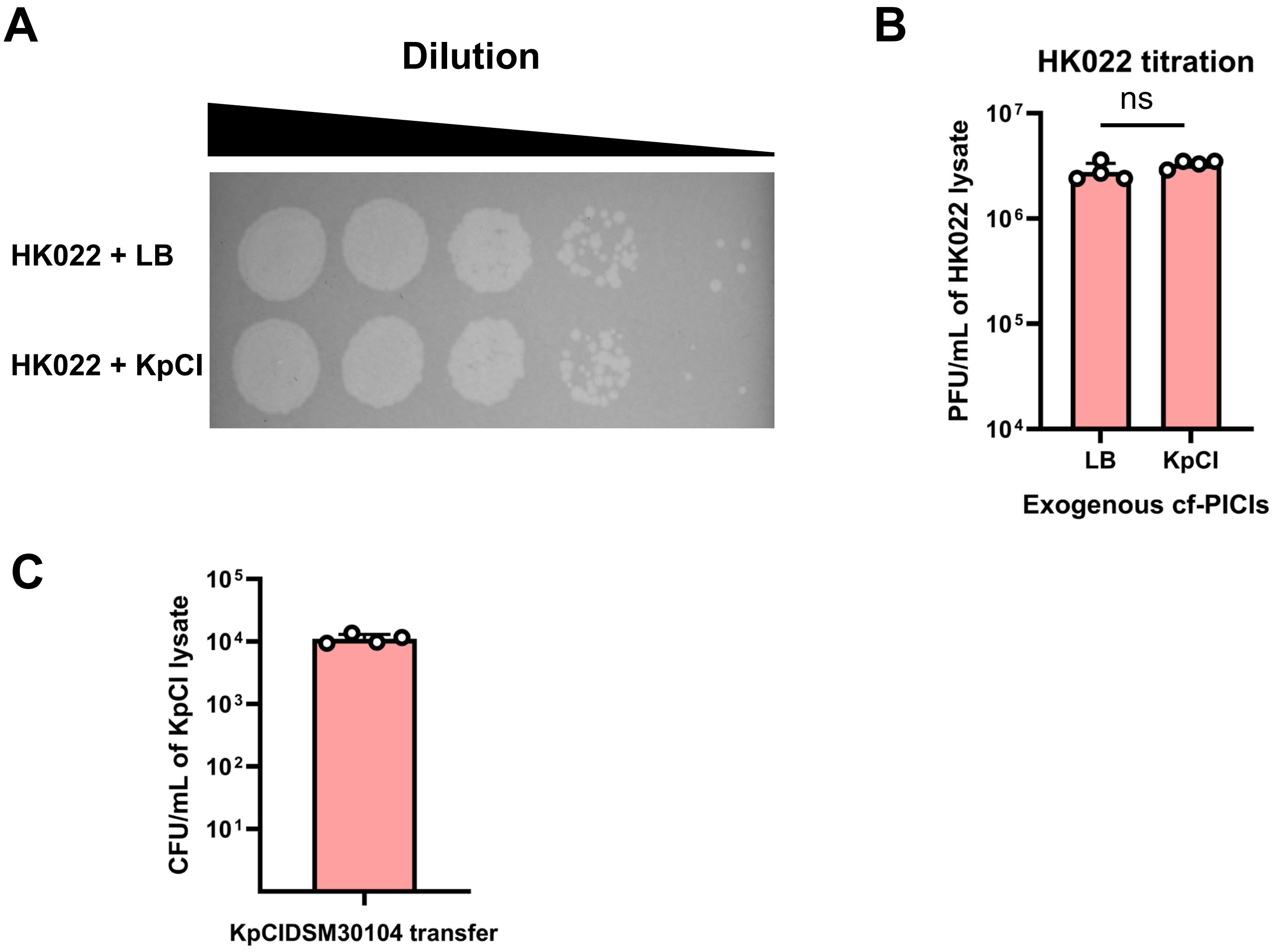


**Figure S5. KpCIDSM30104 uses the excess of tails produced after induction of the HK022 prophage.** Tenfold dilutions of the HK022 lysate (maximum concentration: $10^6$ PFU/mL) were mixed with KpCIDSM30104 ($10^8$ particles/mL) or LB (as a control). The mixed lysates were then used to infect *E. coli* **(A)**, and the number of plaques obtained was quantified **(B)**. A t-test was used to compare the data after log10 transformation. ns: $P > 0.05$.

**(C)** KpCIDSM30104 transfer obtained with the mix that contained the highest concentration of phage lysate ($10^6$ PFU/mL).

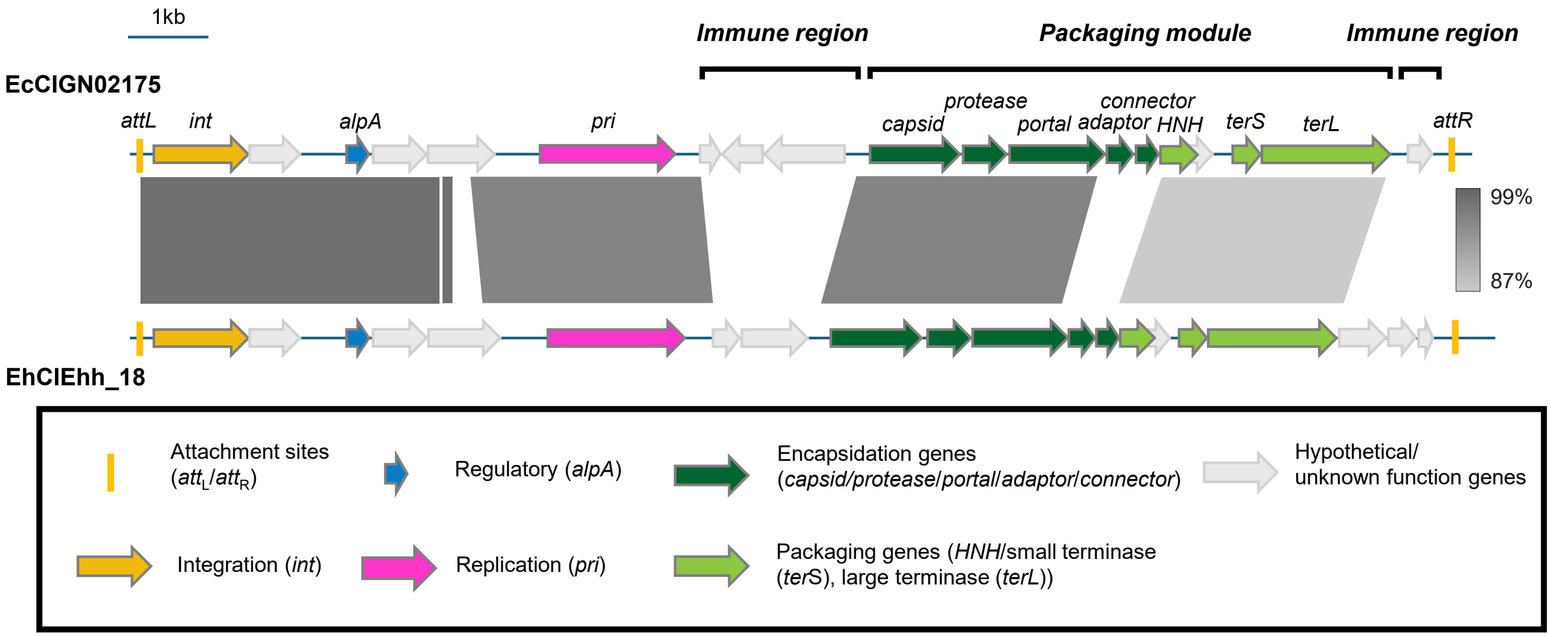


**Figure S6. Comparison between EhCIEhh_18 and EcCIGN02175.** A comparative map between cf-PICIs EcCIGN02175 and EhCIEhh_18. Genes are colored based on their function. Grey scales between cf-PICI sequences indicate regions that share similarity identified by BLASTn.

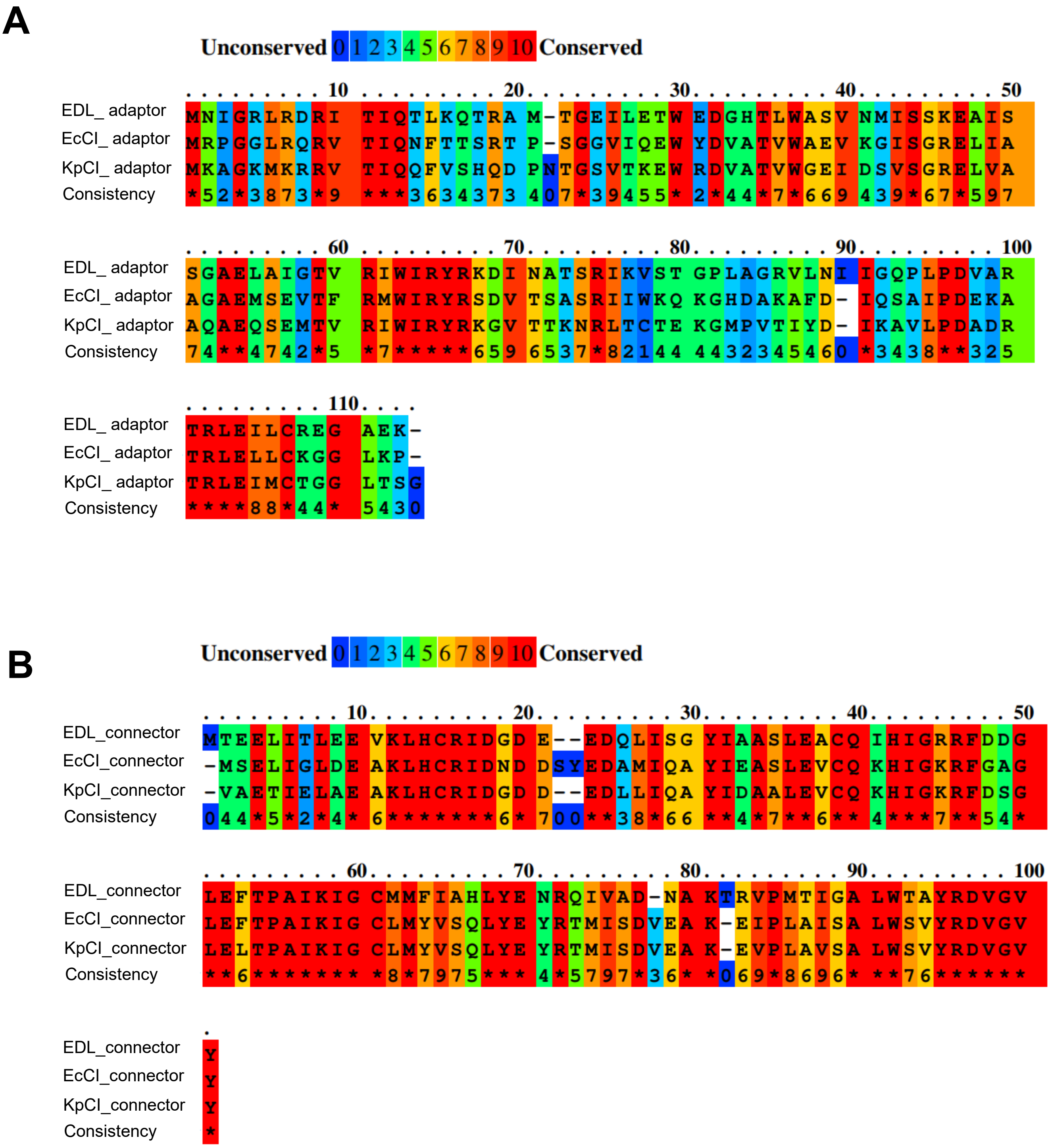


**Figure S7. EcCIEDL933, EcCIGN02175, and KpCIDSM30104 encode different adaptor and connector proteins.**

(A) PRALINE alignment for cf-PICI adaptors.

(B) PRALINE alignment for cf-PICI connectors.

(C) Colors represent the conservation between amino acids. EDL: EcCIEDL933, KpCI: KpCIDSM30104, EcCI: EcCIGN02175.

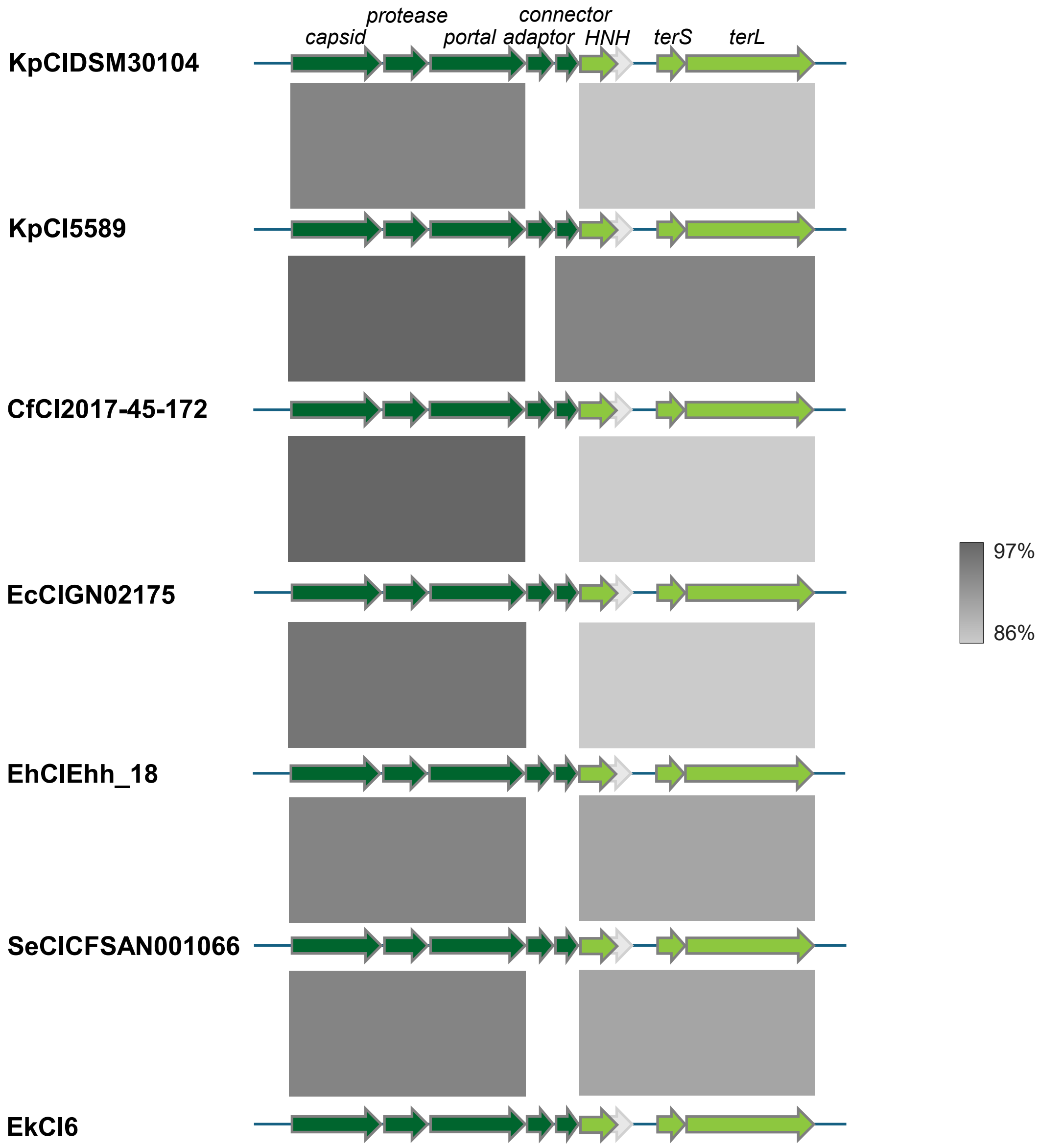


**Figure S8. Comparative maps of the packaging modules present in different cf-PICIs.** cf-PICIs carry almost identical packaging modules but different adaptor and connector genes. Green arrows indicate phage structural proteins. Grey arrows indicate hypothetical proteins. Grey scales between phage sequences indicate regions that share similarity based on BLASTn.

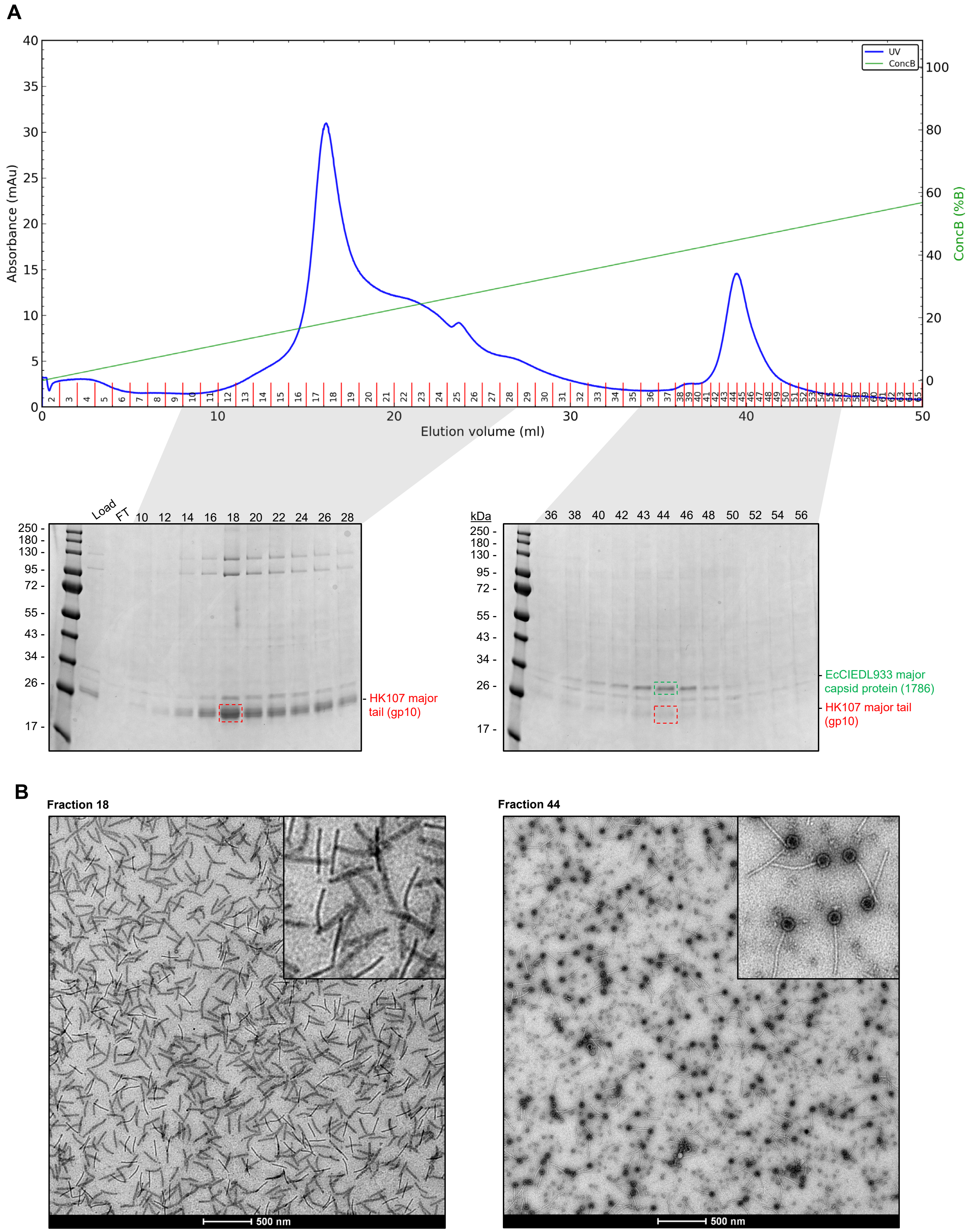


**Figure S9. Purification of EcCIEDL933 via ion-exchange chromatography.**
**(A)** After initial purification by CsCl density gradient centrifugation, EcCIEDL933 virions were subjected to anionic exchange chromatography on HiTrap Q HP Cytvia column to remove the excess unbound tails of the helper phage HK107. The chromatography was conducted at pH 8.4 in a gradient of buffer B containing 2M NaAc relative to buffer A devoid of NaAc, resulting in two distinct peaks. **(B)** Left: The first peak corresponding to low salt elution (fractions 10 to 28) was subjected to SDS-PAGE, revealing the presence of major tail protein (gp10) of phage HK107. Right: The second peak corresponding to high salt elution (fractions 36 to 56) revealed the presence of both, the EcCIEDL933 PICI capsid as well as the major tail protein. **(C)** Fractions 18 and 44 were subjected to negative stain transmission electron microscopy, visualising the HK107 tails, and assembled EcCIEDL933 PICI particles, respectively.

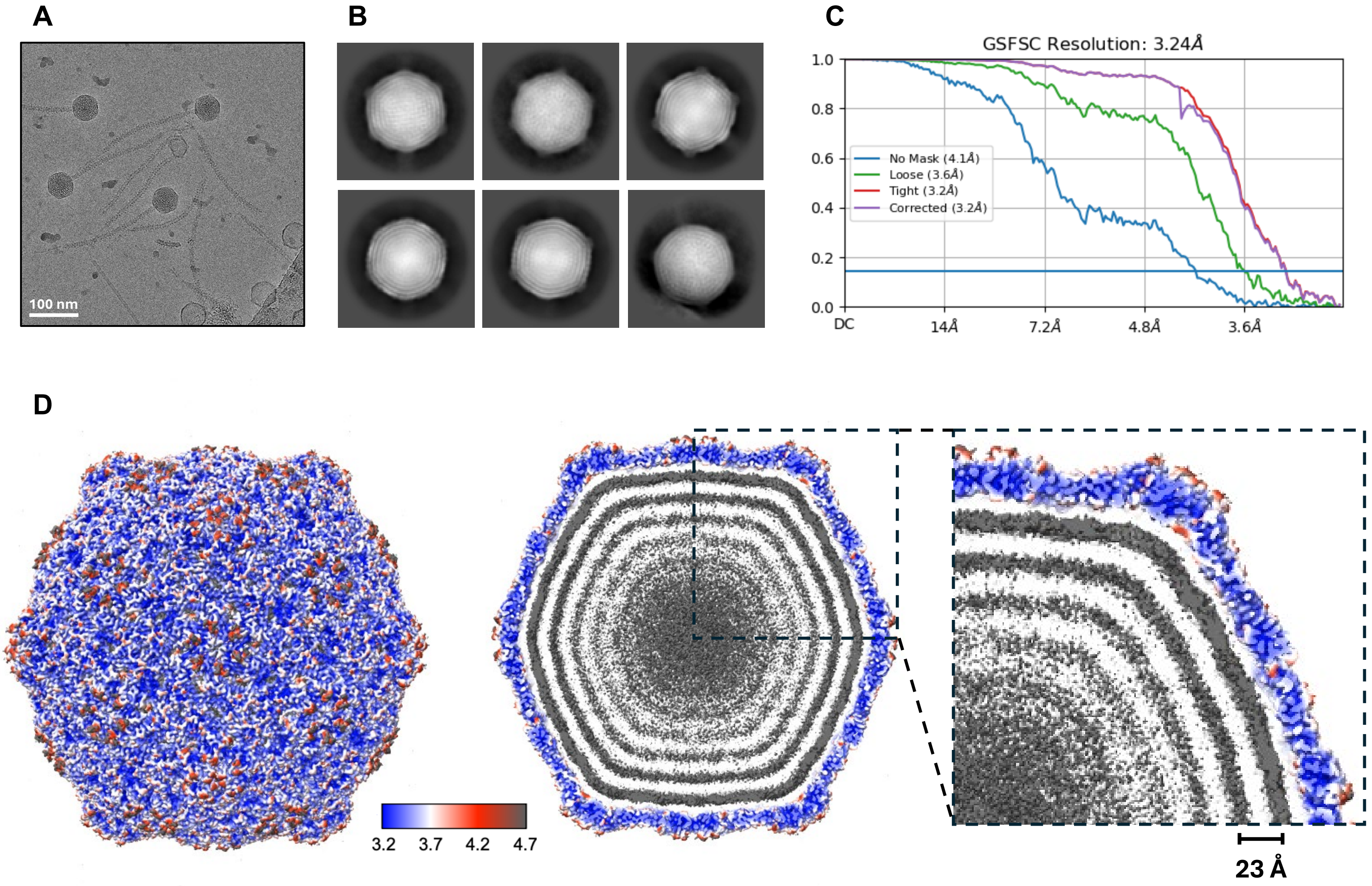


**Figure S10. Reconstruction of a 3D map of EcCIEDL933 capsid**
**(A)** Representative cryo-EM micrograph generated during data collection. **(B)** 2D classes of the capsid used for 3D reconstruction. **(C)** Fourier shell correlation (FSC) curves of the 3D reconstruction. **(D)** Local resolution of the capsid map, with the genome layers distance labelled.

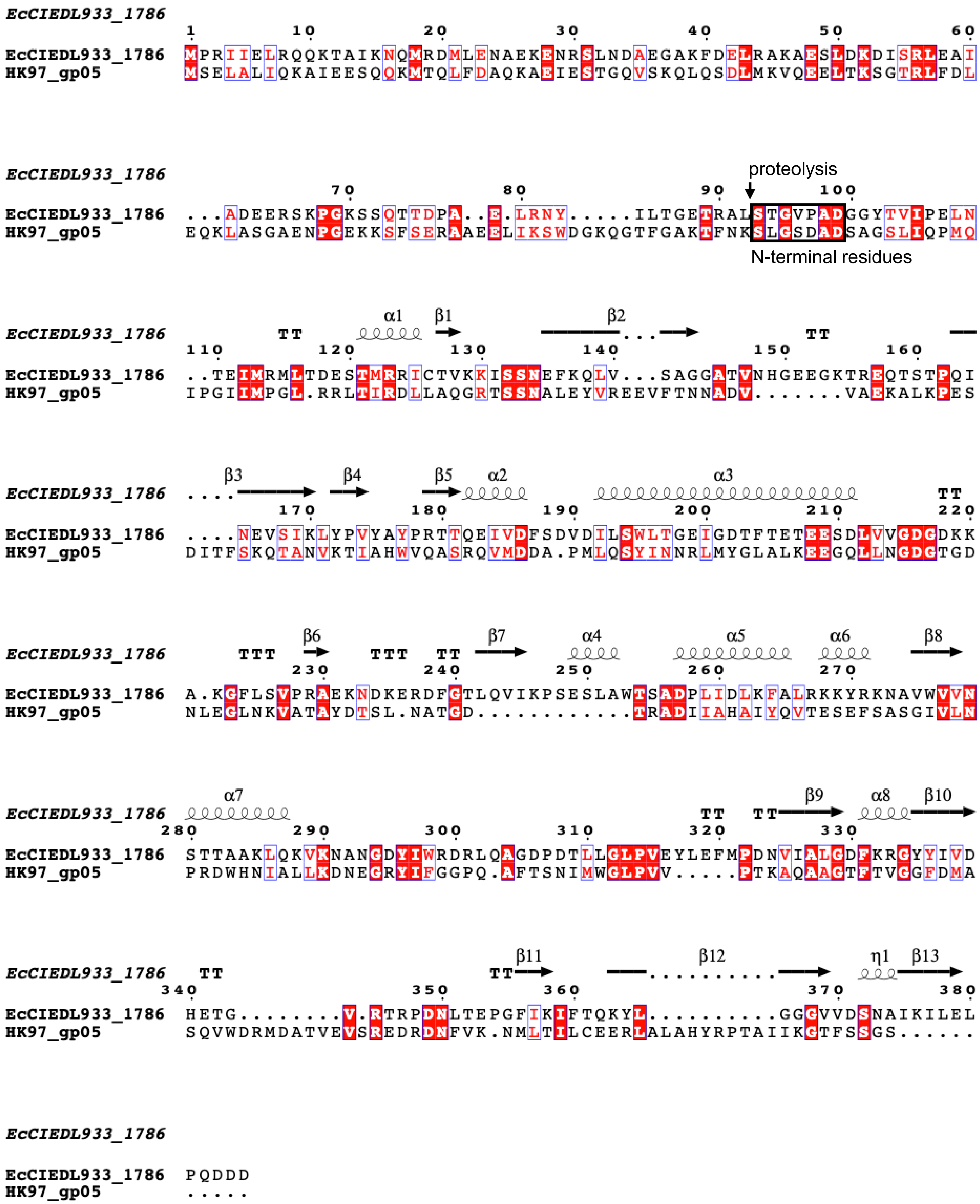


**Figure S11. Amino acid alignment of MCP between EcCIEDL933 and HK97.**
The N-termini of both MCP start with a conserved sequence [SxGxxAD], with x denoting any residue, which suggests a shared proteolysis mechanism.

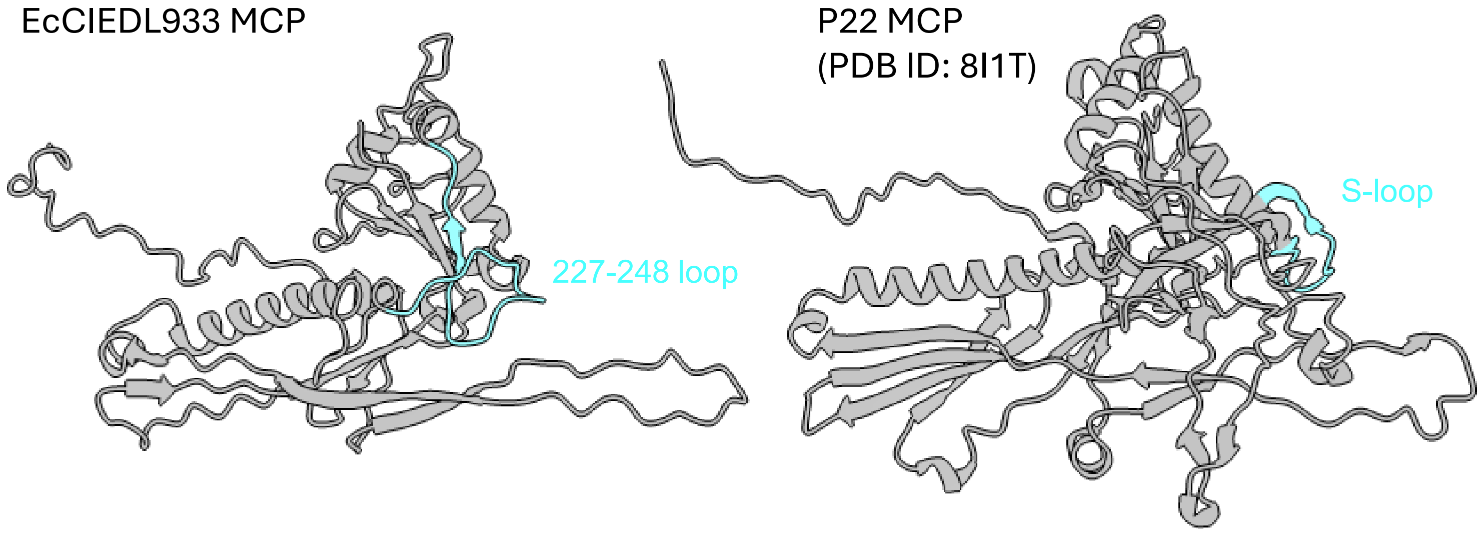


**Figure S12. Localization of the 227-248 loop within MCP of EcCIEDL933 and the S-loop within the P22 bacteriophage.**
The 227-248 loop (cyan) present in EcCIEDL933 MCP localises similarly to the capsid size and symmetry determination loop (S-loop) of the P22 bacteriophage (PDB ID: 8I1T).

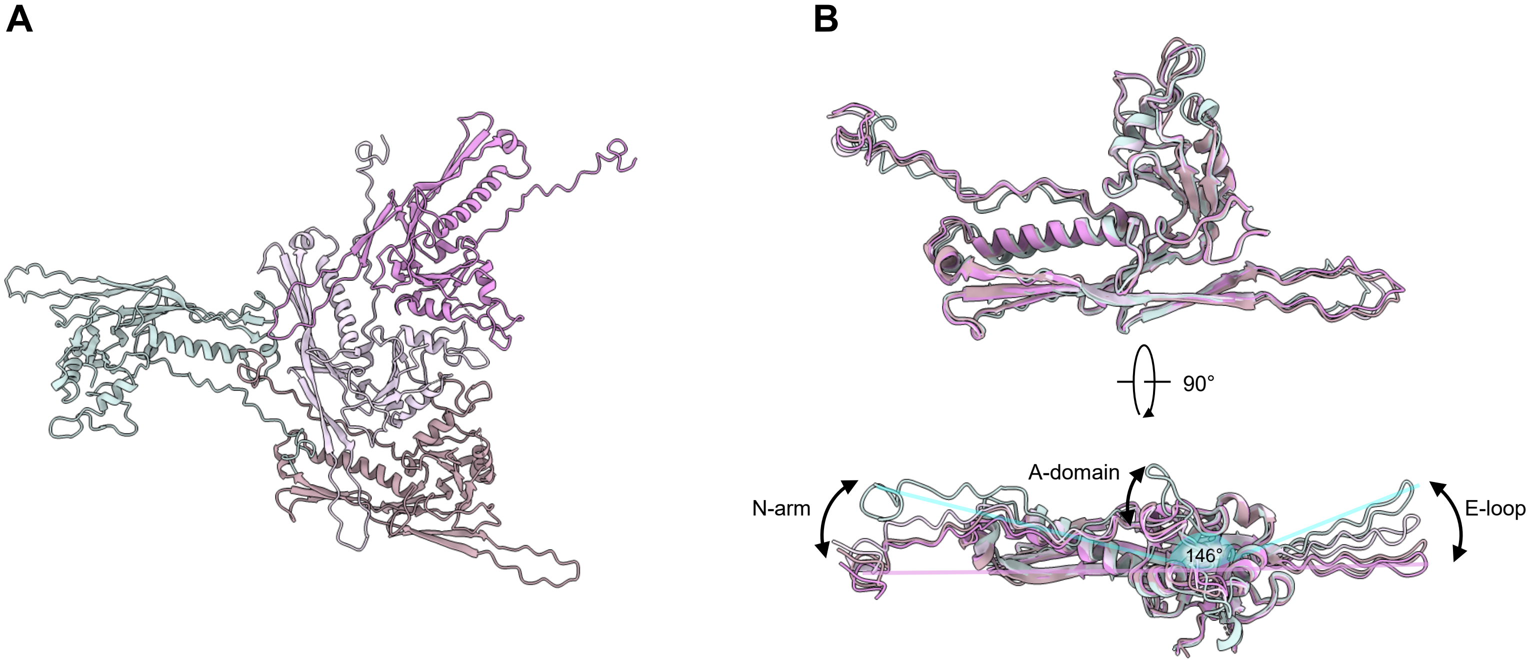


**Figure S13. Variance between MCP from the hexon and penton.**
**(A)** Ribbon representation of the ASU, showing the four MCP. **(B)** Overlay of all the MCP, showing variation primarily in three sites: the N-arm, A-domain and E-loop. It results in different angle between the E-loop and N-arm, causing the penton MCP (mint green) to be the narrowest at an angle of 146° between the E-loop and N-arm, whereas the MCP from the hexon (shades of pink) being flatter, nearing 180° angle between the E-loop and N-arm.

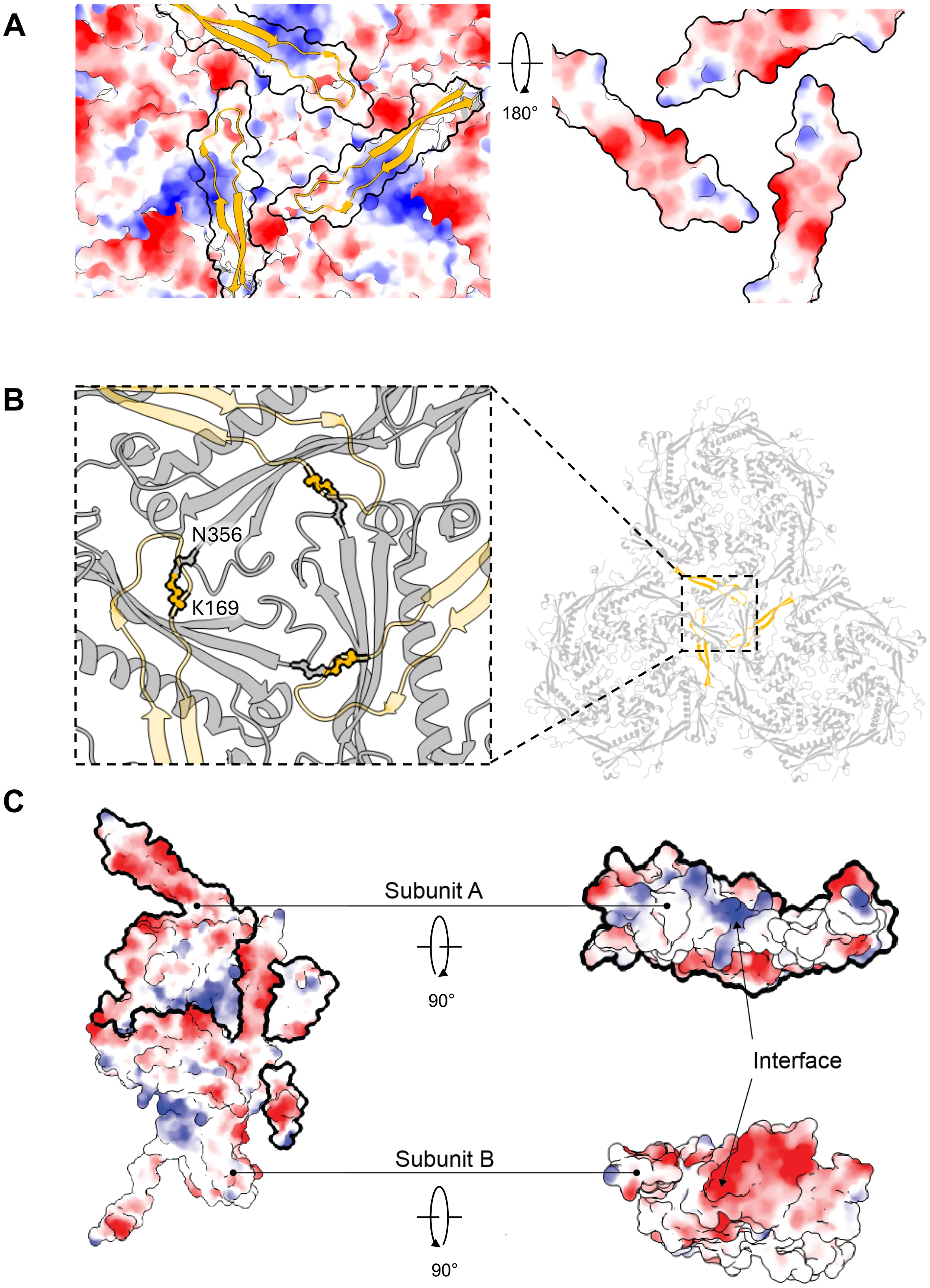


**Figure S14. Comparison of capsomer interactions within the threefold axis in HK97 (PDB ID: 1OHG)**
**(A)** Electrostatic surface potential at the threefold axis except for E-loops (orange ribbon), for which only outlines are shown. The E-loops are shown on the right but rotated 180°. **(B)** In HK97, the interaction between three hexons includes formation of a crosslink between the N-arm and P-domain of the neighboring hexon between residues K169 and N356. **(C)** Within a capsomer, interaction between two HK97 subunits are established by complementary patches at the two faces of the MCP.

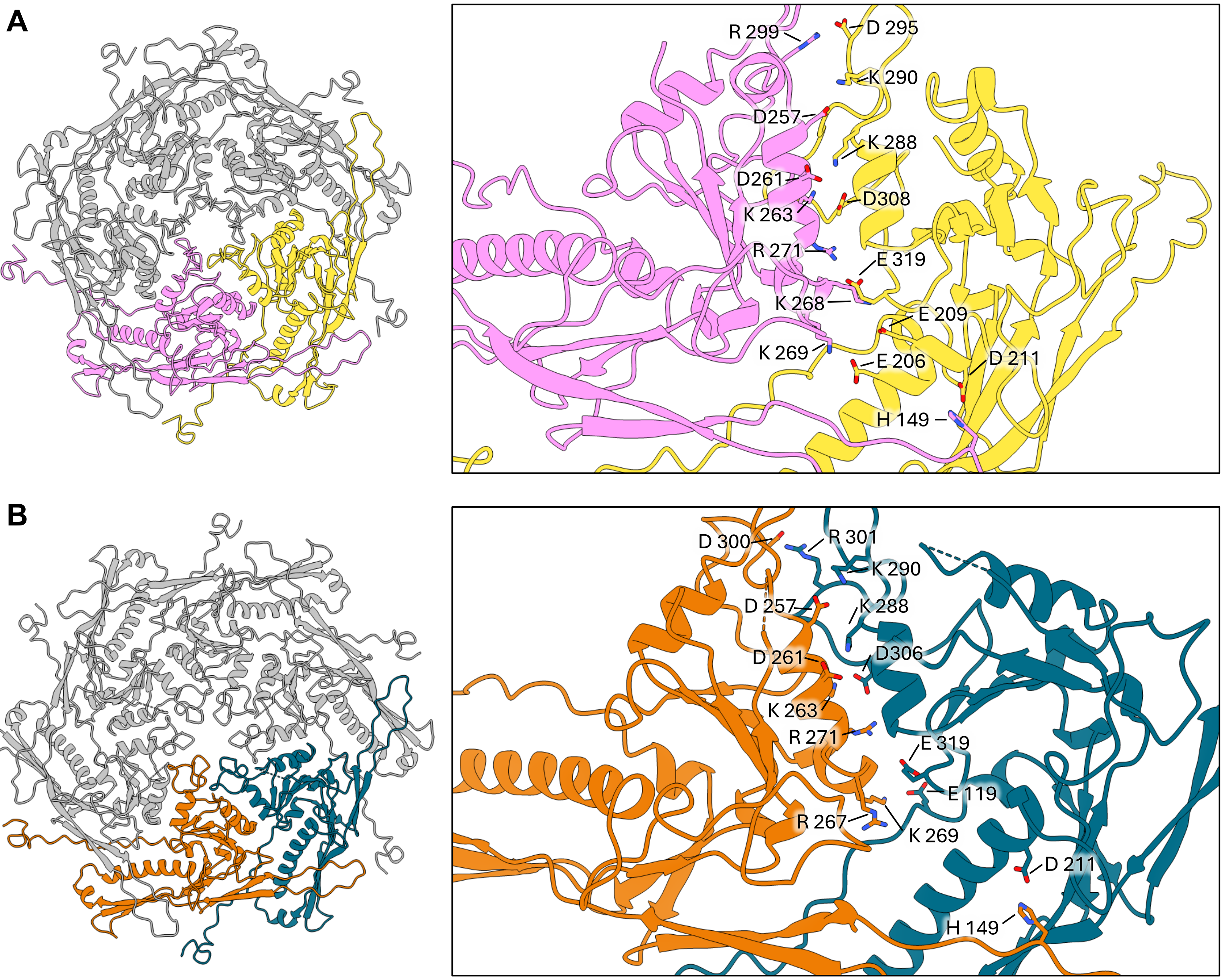


**Figure S15. A network of salt bridges maintains the capsomer integrity**
Salt bridges formed between two MCP within a penton **(A)** and hexon **(B)** are shown, with the specific interacting residues highlighted.

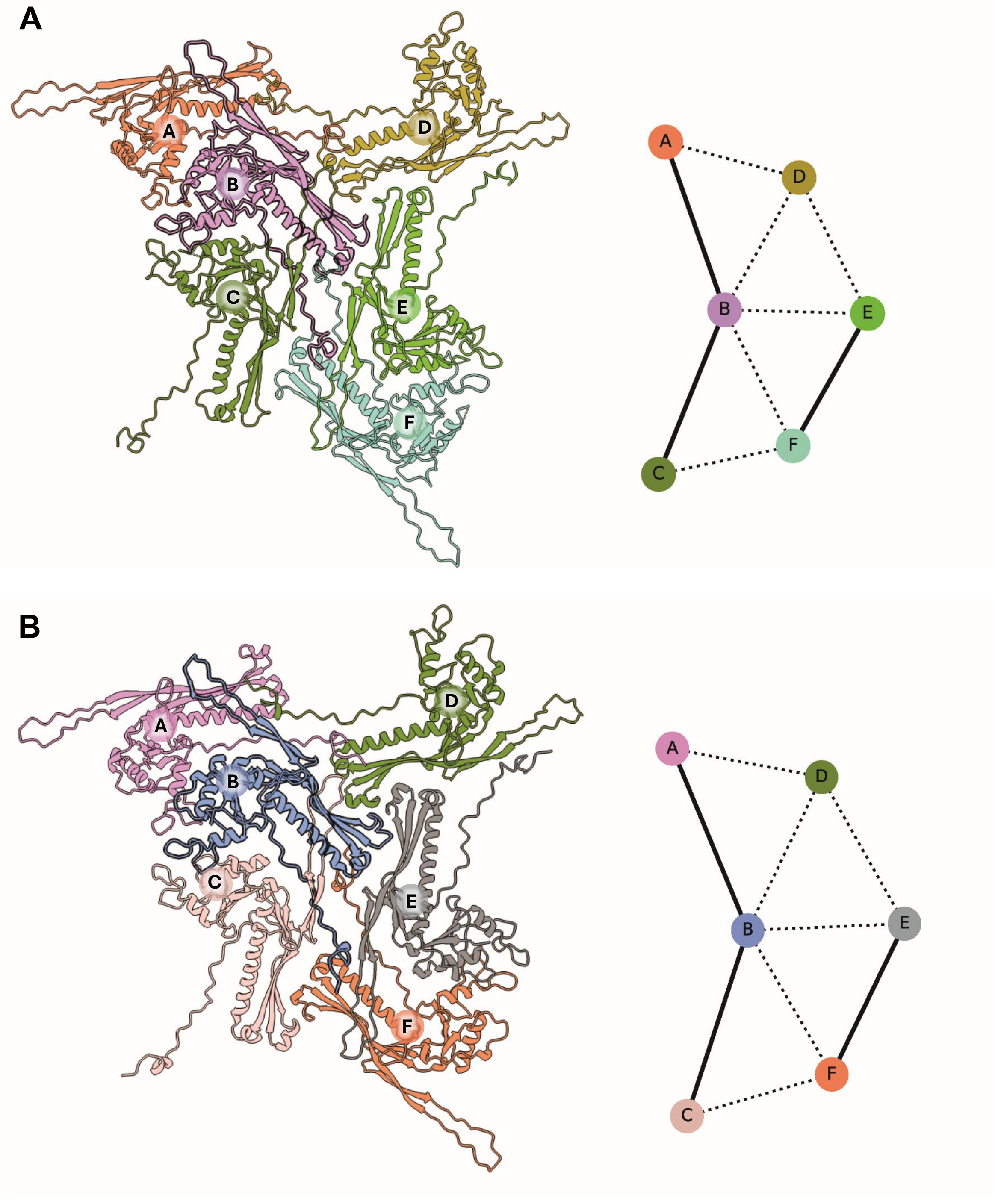


**Figure S16. Representation of unique interactions of a single MCP**

Any single MCP (blue) interacts with five neighbouring MCP. Shown are the unique interactions between EcCIEDL933 MCP **(A)** and HK97 MCP (PDB ID: 1OHG) **(B)**. Detailed interactions are listed in table S5 (EcCIEDL933) and table S6 (HK97).
