## Supplementary Tables for "Chimeric infective particles expand species boundaries in phage inducible chromosomal island mobilization"

**Table S1. EcCIGN02175 is present in presented in five different bacterial genera and seven different bacterial species**. See also Figure S1.

| **Strain** | **Species** | **Accession number (genomic location)** | **Size** | **% identity with EcCIGN2175** |
|---|---|---|---|---|
| GN02175 | *Escherichia coli* | CP041550 (1,227,235-1,243,472) | 16.238 | 100% |
| GN06667 | *Escherichia coli* | CP147587 (472,705-488,942) | 16.238 | 100% |
| GN05476 | *Escherichia coli* | CP147578 (472,705-488,942) | 16.238 | 100% |
| CU37RT-2 | *Escherichia coli* | CP082774 (4,487,602-4,503,839) | 16.238 | 100% |
| RHB36-C16 | *Escherichia coli* | CP057126 (4,126,048-4,142,285) | 16.238 | 99,94% |
| OW1E2 | *Escherichia coli* | CP067245 (3,551,394-3,567,650) | 16.257 | 99,75% |
| TUM9754 | *Escherichia coli* | AP026487 (526,112-543,126) | 17.015 | 99,94% |
| E706 | *Escherichia coli* | CP029687 (465,358-482,372) | 17.015 | 99,94% |
| DETEC-P61 | *Escherichia coli* | CP116152 (463,811-480,825) | 17.015 | 99,94% |
| SWHIN_97 | *Shigella flexneri* | CP055070 (471,659-487,896) | 16.238 | 99,93% |
| TH12880 | *Klebsiella pneumoniae* | CP098771 (478,853-496,172) | 17.320 | 99,68% |
| 2017-45-172 | *Citrobacter freundii* | CP109711 (2,048,803-2,065,398) | 16.596 | 98,97% |
| RHBSTW-00449 | *Citrobacter freundii* | CP056505 (439,836-456,465) | 16.630 | 98,10% |
| 2023CK-01304 | *Citrobacter amalonaticus* | CP140489 (501,064-517,693) | 16.630 | 98,10% |
| 161373-yvys | *Enterobacter asburiae* | CP083815 (484,601-501,562) | 16.962 | 97,74% |
| 48411CZ | *Enterobacter hormaechei* | CP085780 (481,399-498,370) | 16.972 | 97,74% |

**Table S2. KpCIDSM30104 is present in different *K. pneumoniae* strains.**

| Strain | Species | Accession number (genomic location) | Size | % identity with KpCIDSM30104 |
|---|---|---|---|---|
| DSM30104 | *Klebsiella pneumoniae* | AJJI01000015 (416,261-432,888) | 16.628 | 100% |
| FDAARGOS_775 | *Klebsiella pneumoniae* | CP040993 (758,208-774,835) | 16.628 | 100% |
| PartO-Kpneumoniae-RM8376 | *Klebsiella pneumoniae* | CP064368 (3,630,781-3,647,408) | 16.628 | 100% |
| 64EVAM | *Klebsiella pneumoniae* | CP138431 (1,140,465-1,157,201) | 16.737 | 97% |
| RHBSTW-00062 | *Klebsiella pneumoniae* | CP056883 (467,470-484,206) | 16.737 | 96,85% |
| RHBSTW-00066 | *Klebsiella pneumoniae* | CP056878 (467,636-485,686) | 18.051 | 96,79% |

**Table S3. Presence of cf-PICIs in different metagenome studies.**

| Name | Accession number |
|---|---|
| uncultured phage genome assembly | OY731419.1 |
| uncultured phage genome assembly | OY731324.1 |
| ctqb520 | BK035691.1 |
| ctBIT13 | BK056511.1 |
| ctWBm6 | BK040613.1 |
| ct8dt12 | BK055826.1 |
| ctIYk15 | BK036617.1 |
| ctCqx10 | BK042697.1 |
| ctI0B6 | BK016321.1 |
| ctqTp37 | BK016502.1 |
| ctV4X6 | BK029089.1 |
| ctDoD23 | BK057057.1 |
| ctNmW2 | BK014912.1 |
| ctskP2 | BK047809.1 |
| ctgME2 | BK053201.1 |
| ctAwx12 | BK054842.1 |
| ctXsn5 | BK051976.1 |
| ctjOi9 | BK039775.1 |
| ctA3K9 | BK030312.1 |
| ctLqq2 | BK036110.1 |
| ctCod3 | BK046431.1 |
| ct5xD5 | BK039715.1 |
| ctXUn4 | BK050260.1 |
| ctBtt10 | BK036015.1 |
| ctDTK2 | BK034082.1 |
| cth3P7 | BK037448.1 |
| ctr0H5 | BK050963.1 |
| ctWUP3 | BK038032.1 |
| ctvf76 | BK022514.1 |
| ctGTD11 | BK035713.1 |
| ctPLn4 | BK042919.1 |
| ctW8q3 | BK041039.1 |
| ctBHz3 | BK038767.1 |
| ctP4E8 | BK058904.1 |
| ctOoJ1 | BK040942.1 |
| ctMT91 | BK049712.1 |
| ct0iY8 | BK043230.1 |
| cthPE4 | BK058346.1 |
| ct8gU3 | BK022567.1 |
| ctCQc2 | BK022045.1 |
| ct7I44 | BK034268.1 |
| ctRiz9 | BK044283.1 |
| ctsvX4 | BK033226.1 |
| ctvmC7 | BK049961.1 |
| ctZhr26 | BK032765.1 |
| ctoYp11 | BK059021.1 |

| | |
|---|---|
| ct2Ak3 | BK027566.1 |
| ctYnq1 | BK024041.1 |
| 2AX_6 | MF417957.1 |
| 7S_14 | MF417956.1 |
| 3F_1 | MF417951.1 |
| ctXus3 | BK016482.1 |
| ctNdu3 | BK031488.1 |
| ctiiV5 | BK029820.1 |
| ctrvD4 | BK054608.1 |
| ctJVj2 | BK058184.1 |
| ctF9y1 | BK019987.1 |
| ctiN128 | BK043660.1 |
| ctdJ013 | BK020212.1 |
| ctQZ210 | BK032445.1 |
| ctQZN6 | BK047365.1 |
| ctWWs5 | BK037892.1 |
| ctOXf3 | BK052879.1 |
| ctzCX1 | BK053308.1 |
| ctLYN8 | BK036917.1 |
| ctRtg9 | BK044109.1 |
| ctO2w5 | BK059028.1 |
| ctrqs2 | BK024841.1 |
| ctGoc3 | BK052708.1 |
| ctxN63 | BK016376.1 |
| ctXFH9 | BK037637.1 |
| ctLCL3 | BK052714.1 |
| ctCFC3 | BK026343.1 |
| ct34I1 | BK021781.1 |
| ctQMp1 | BK020760.1 |
| 2905_15832 | OP072448.1 |
| 2718_76661 | OP074827.1 |
| 1519_97203 | OP073998.1 |
| ctJsG2 | BK031250.1 |

**Table S4. Data collection, refinement parameters, and model validation statistics of the EcCIEDL933 capsid structure.**

| | |
|---|---|
| **Processing and Refinement Parameters** | |
| Voltage (kV) | 200 |
| Total number of micrographs | 4377 |
| Reconstruction software | CryoSPARC |
| Defocus range (µm) | -0.8 to -2.8 |
| Electron dose (e−/Å$^2$) | 50 |
| Frames/micrograph | 40 |
| Pixel size (Å/px) | 1.1 |
| Number of particles used for final map | 1993 |
| Reconstruction symmetry | I |
| Resolution of final map (Å) | 3.24 |
| **Deposition** | |
| PDB ID | 9GYI |
| Residue range | 93-327 |
| Map corelation coefficient | 0.8 |
| **Bonds (RMSD)** | |
| Length (Å) | 0.003 |
| Angles (°) | 0.638 |
| **Clash score** | 7.13 |
| **MolProbity score** | 1.73 |
| **Ramachandran plot** | |
| Outliers (%) | 0 |
| Allowed (%) | 4.86 |
| Favored (%) | 95.14 |
| Rotamer outliers (%) | 0 |
| CaBLAM outliers (%) | 1.36 |

**Table S5. Interactions within EcCIEDL933.**

| **B vs A** | | | |
|---|---|---|---|
| *Hydrogen Bonds* | | | |
| 1 | A:GLU 107[ N ] | 3.04 | B:GLU 135[ O ] |
| 2 | A:ASN 109[ N ] | 3.56 | B:LYS 137[ O ] |
| 3 | A:MET 113[ N ] | 3.59 | B:LEU 139[ O ] |
| 4 | A:TYR 177[ N ] | 3.49 | B:THR 159[ OG1] |
| 5 | A:ARG 179[ N ] | 3.48 | B:GLN 158[ OE1] |
| 6 | A:TRP 195[ NE1] | 3.21 | B:SER 141[ OG ] |
| 7 | A:LYS 219[ NZ ] | 3.21 | B:GLU 151[ OE1] |
| 8 | A:SER 280[ OG ] | 3.12 | B:PHE 264[ O ] |
| 9 | A:LYS 288[ NZ ] | 2.79 | B:ASP 261[ OD2] |
| 10 | A:LYS 290[ NZ ] | 3.14 | B:ASP 257[ OD2] |
| 11 | A:TYR 296[ OH ] | 3.72 | B:ASP 257[ OD1] |
| 12 | A:ARG 301[ NH1] | 3.12 | B:ASP 300[ OD1] |
| 13 | A:ARG 301[ NH2] | 3.31 | B:ASP 300[ OD1] |
| 14 | A:GLN 303[ NE2] | 2.43 | B:TRP 298[ O ] |
| 15 | A:GLU 107[ O ] | 3.40 | B:LYS 137[ N ] |
| 16 | A:GLU 107[ OE1] | 2.88 | B:SER 132[ N ] |
| 17 | A:GLU 107[ OE1] | 3.88 | B:SER 133[ OG ] |
| 18 | A:ASN 109[ OD1] | 3.78 | B:GLN 138[ NE2] |
| 19 | A:MET 113[ O ] | 3.18 | B:SER 141[ OG ] |
| 20 | A:MET 113[ O ] | 3.48 | B:SER 141[ N ] |
| 21 | A:ARG 114[ O ] | 3.16 | B:ARG 333[ NH1] |
| 22 | A:ARG 114[ O ] | 3.12 | B:ARG 333[ NH2] |
| 23 | A:MET 115[ SD ] | 3.39 | B:GLY 143[ N ] |
| 24 | A:LEU 116[ O ] | 3.64 | B:ARG 267[ NH1] |
| 25 | A:THR 117[ O ] | 3.57 | B:LYS 269[ NZ ] |
| 26 | A:ASP 118[ OD1] | 2.58 | B:ARG 267[ NH1] |
| 27 | A:ASP 118[ OD2] | 3.20 | B:ARG 267[ NH2] |
| 28 | A:TYR 175[ OH ] | 3.02 | B:GLU 157[ N ] |
| 29 | A:GLU 199[ OE2] | 3.14 | B:SER 141[ OG ] |
| 30 | A:SER 280[ OG ] | 2.77 | B:ARG 271[ NH2] |
| 31 | A:ASN 293[ O ] | 3.03 | B:ALA 292[ N ] |
| 32 | A:ASP 295[ OD1] | 2.68 | B:ASN 291[ ND2] |
| 33 | A:GLN 303[ O ] | 3.12 | B:THR 309[ OG1] |
| 34 | A:GLN 303[ O ] | 2.83 | B:GLY 312[ N ] |
| 35 | A:ASP 306[ OD1] | 2.55 | B:LYS 263[ NZ ] |

| | | | |
|---|---|---|---|
| 36 | A:GLN 362[ OE1] | 3.49 | B:ARG 156[ NH1] |

*Salt Bridges*

| | | | |
|---|---|---|---|
| 1 | A:LYS 219[ NZ ] | 3.21 | B:GLU 151[ OE1] |
| 2 | A:LYS 219[ NZ ] | 3.50 | B:GLU 151[ OE2] |
| 3 | A:LYS 288[ NZ ] | 3.88 | B:ASP 261[ OD1] |
| 4 | A:LYS 288[ NZ ] | 2.79 | B:ASP 261[ OD2] |
| 5 | A:LYS 290[ NZ ] | 3.01 | B:ASP 257[ OD1] |
| 6 | A:LYS 290[ NZ ] | 3.14 | B:ASP 257[ OD2] |
| 7 | A:ARG 301[ NH1] | 3.12 | B:ASP 300[ OD1] |
| 8 | A:ARG 301[ NH2] | 3.31 | B:ASP 300[ OD1] |
| 9 | A:ASP 118[ OD1] | 2.58 | B:ARG 267[ NH1] |
| 10 | A:ASP 118[ OD1] | 3.67 | B:ARG 267[ NH2] |
| 11 | A:ASP 118[ OD2] | 3.51 | B:ARG 267[ NH1] |
| 12 | A:ASP 118[ OD2] | 3.20 | B:ARG 267[ NH2] |
| 13 | A:ASP 306[ OD1] | 2.55 | B:LYS 263[ NZ ] |
| 14 | A:ASP 306[ OD2] | 3.74 | B:LYS 263[ NZ ] |

**B vs C**

*Hydrogen Bonds*

| | | | |
|---|---|---|---|
| 1 | B:ASN 109[ N ] | 3.27 | C:LYS 137[ O ] |
| 2 | B:MET 113[ N ] | 3.01 | C:LEU 139[ O ] |
| 3 | B:TYR 175[ OH ] | 2.99 | C:GLU 157[ OE1] |
| 4 | B:TYR 177[ N ] | 3.19 | C:THR 159[ OG1] |
| 5 | B:PRO 178[ N ] | 3.65 | C:THR 159[ OG1] |
| 6 | B:ARG 179[ NE ] | 3.53 | C:GLN 158[ OE1] |
| 7 | B:TRP 195[ NE1] | 3.04 | C:SER 141[ OG ] |
| 8 | B:LYS 288[ NZ ] | 3.10 | C:ASP 261[ OD2] |
| 9 | B:LYS 290[ NZ ] | 3.16 | C:ASP 257[ OD1] |
| 10 | B:LYS 290[ NZ ] | 2.89 | C:ASP 257[ OD2] |
| 11 | B:ASP 295[ N ] | 2.74 | C:ASN 291[ OD1] |
| 12 | B:TYR 296[ OH ] | 3.48 | C:ASP 257[ OD1] |
| 13 | B:ARG 301[ NH1] | 3.33 | C:ARG 299[ O ] |
| 14 | B:ARG 301[ NH1] | 3.06 | C:ASP 300[ O ] |
| 15 | B:ARG 301[ NH1] | 3.64 | C:ASP 300[ OD1] |
| 16 | B:ARG 301[ NH2] | 2.52 | C:TYR 296[ O ] |
| 17 | B:GLN 303[ NE2] | 2.85 | C:TRP 298[ O ] |
| 18 | B:PRO 106[ O ] | 3.60 | C:LYS 137[ N ] |
| 19 | B:GLU 107[ OE2] | 2.71 | C:SER 132[ OG ] |

| | | | |
|---|---|---|---|
| 20 | B:GLU 107[ OE2] | 3.31 | C:SER 132[ N ] |
| 21 | B:ASN 109[ OD1] | 3.68 | C:GLN 138[ NE2] |
| 22 | B:MET 113[ O ] | 3.85 | C:SER 141[ OG ] |
| 23 | B:MET 113[ O ] | 3.53 | C:SER 141[ N ] |
| 24 | B:MET 113[ SD ] | 3.04 | C:GLN 138[ NE2] |
| 25 | B:ARG 114[ O ] | 2.61 | C:ARG 333[ NH2] |
| 26 | B:TYR 175[ O ] | 3.76 | C:ASN 148[ N ] |
| 27 | B:TYR 175[ OH ] | 3.70 | C:LYS 154[ NZ ] |
| 28 | B:TYR 175[ OH ] | 3.63 | C:GLU 157[ N ] |
| 29 | B:TYR 175[ OH ] | 3.03 | C:ASN 148[ ND2] |
| 30 | B:TYR 177[ O ] | 3.82 | C:THR 159[ N ] |
| 31 | B:SER 280[ OG ] | 3.82 | C:ARG 271[ NH2] |
| 32 | B:ASN 293[ O ] | 3.38 | C:ASN 293[ N ] |
| 33 | B:ASN 293[ O ] | 3.47 | C:ASN 293[ ND2] |
| 34 | B:ASN 293[ OD1] | 2.54 | C:ASN 293[ ND2] |
| 35 | B:TYR 296[ OH ] | 3.47 | C:ASP 257[ N ] |
| 36 | B:LEU 302[ O ] | 2.52 | C:LEU 311[ N ] |
| 37 | B:GLN 303[ O ] | 2.93 | C:THR 309[ OG1] |
| 38 | B:GLU 319[ OE1] | 3.89 | C:ARG 267[ NE ] |

*Salt Bridges*

| | | | |
|---|---|---|---|
| 1 | B:LYS 288[ NZ ] | 3.42 | C:ASP 261[ OD1] |
| 2 | B:LYS 288[ NZ ] | 3.10 | C:ASP 261[ OD2] |
| 3 | B:LYS 290[ NZ ] | 3.16 | C:ASP 257[ OD1] |
| 4 | B:LYS 290[ NZ ] | 2.89 | C:ASP 257[ OD2] |
| 5 | B:ARG 301[ NH1] | 3.64 | C:ASP 300[ OD1] |
| 6 | B:GLU 119[ OE1] | 3.23 | C:LYS 269[ NZ ] |
| 7 | B:ASP 211[ OD1] | 3.36 | C:HIS 149[ ND1] |
| 8 | B:ASP 211[ OD2] | 3.75 | C:HIS 149[ ND1] |
| 9 | B:ASP 306[ OD1] | 2.84 | C:LYS 263[ NZ ] |
| 10 | B:ASP 306[ OD2] | 3.71 | C:LYS 263[ NZ ] |
| 11 | B:ASP 306[ OD2] | 3.95 | C:ARG 271[ NH1] |
| 12 | B:GLU 319[ OE1] | 3.89 | C:ARG 267[ NE ] |
| 13 | B:GLU 319[ OE2] | 3.99 | C:ARG 267[ NH2] |

**B vs D**

*Hydrogen Bonds*

| | | | |
|---|---|---|---|
| 1 | B:ILE 164[ N ] | 3.09 | D:ALA 98[ O ] |
| 2 | B:GLN 163[ NE2] | 2.63 | D:ASP 99[ OD1] |

| | | | |
|---|---|---|---|
| 3 | B:LYS 219[ NZ ] | 3.29 | D:GLU 183[ OE1] |
| 4 | B:HIS 340[ NE2] | 3.32 | D:ASP 186[ O ] |
| 5 | B:ARG 345[ NH2] | 3.36 | D:ASP 186[ OD1] |
| 6 | B:ARG 345[ NH1] | 3.36 | D:ASP 186[ OD1] |
| 7 | B:ARG 347[ NH1] | 3.31 | D:ASN 350[ O ] |

*Salt Bridges*

| | | | |
|---|---|---|---|
| 1 | B:LYS 219[ NZ ] | 3.29 | D:GLU 183[ OE1] |
| 2 | B:ARG 345[ NH2] | 3.36 | D:ASP 186[ OD1] |
| 3 | B:ARG 345[ NH1] | 3.36 | D:ASP 186[ OD1] |
| 4 | B:ARG 345[ NH2] | 3.71 | D:ASP 186[ OD2] |
| 5 | B:ARG 345[ NE ] | 3.84 | D:ASP 186[ OD2] |
| 6 | B:ARG 345[ NH1] | 3.02 | D:ASP 186[ OD2] |

**B vs E**

*Hydrogen Bonds*

| | | | |
|---|---|---|---|
| 1 | B:ALA 98[ O ] | 2.40 | E:ILE 164[ N ] |
| 2 | B:ASP 99[ OD1] | 3.32 | E:THR 161[ OG1] |
| 3 | B:ASP 186[ OD2] | 2.63 | E:ARG 345[ NH2] |
| 4 | B:ASN 350[ O ] | 2.65 | E:ARG 347[ NE ] |

*Salt bridges*

| | | | |
|---|---|---|---|
| 1 | B:ASP 186[ OD1] | 3.99 | E:ARG 345[ NE ] |
| 2 | B:ASP 186[ OD1] | 3.81 | E:ARG 345[ NH2] |
| 3 | B:ASP 186[ OD2] | 3.77 | E:ARG 345[ NH1] |
| 4 | B:ASP 186[ OD2] | 2.63 | E:ARG 345[ NH2] |

**B vs F**

*Hydrogen Bonds*

| | | | |
|---|---|---|---|
| 1 | B:SER 93[ N ] | 3.66 | F:ARG 179[ O ] |
| 2 | B:THR 103[ OG1] | 2.71 | F:SER 188[ OG ] |
| 3 | B:THR 181[ N ] | 3.03 | F:SER 93[ O ] |
| 4 | B:SER 93[ O ] | 3.29 | F:THR 181[ N ] |
| 5 | B:SER 93[ O ] | 3.59 | F:THR 181[ OG1] |
| 6 | B:THR 103[ O ] | 3.04 | F:ASP 189[ N ] |

**Table S6. Interactions within HK97.**

| **B vs A** | | | |
|---|---|---|---|
| *Hydrogen Bonds* | | | |
| 1 | A:MET 119[ N ] | 3.20 | B:ALA 147[ O ] |
| 2 | A:ILE 121[ N ] | 2.87 | B:GLU 149[ O ] |
| 3 | A:ILE 125[ N ] | 2.84 | B:VAL 151[ O ] |
| 4 | A:ALA 187[ N ] | 2.93 | B:ASP 161[ O ] |
| 5 | A:HIS 188[ ND1] | 3.30 | B:SER 172[ OG ] |
| 6 | A:TRP 189[ N ] | 2.99 | B:PRO 170[ O ] |
| 7 | A:TRP 189[ NE1] | 3.19 | B:LYS 169[ O ] |
| 8 | A:TYR 206[ OH ] | 2.54 | B:GLU 153[ OE1] |
| 9 | A:ARG 210[ NE ] | 2.90 | B:GLU 153[ OE1] |
| 10 | A:ARG 210[ NH2] | 3.05 | B:GLU 153[ OE2] |
| 11 | A:ARG 280[ NH2] | 3.04 | B:SER 246[ OG ] |
| 12 | A:ARG 294[ NE ] | 3.44 | B:ASP 290[ OD2] |
| 13 | A:ARG 294[ NH2] | 3.29 | B:ASP 290[ OD2] |
| 14 | A:TYR 295[ OH ] | 2.54 | B:ASP 256[ OD2] |
| 15 | A:GLN 301[ NE2] | 3.90 | B:THR 304[ OG1] |
| 16 | A:GLN 301[ NE2] | 3.75 | B:PHE 297[ O ] |
| 17 | A:LYS 317[ NZ ] | 3.07 | B:GLU 267[ O ] |
| 18 | A:GLN 117[ O ] | 3.47 | B:SER 145[ OG ] |
| 19 | A:MET 119[ O ] | 3.04 | B:GLU 149[ N ] |
| 20 | A:MET 119[ SD ] | 3.32 | B:THR 143[ OG1] |
| 21 | A:ILE 125[ O ] | 3.53 | B:GLU 153[ N ] |
| 22 | A:MET 126[ O ] | 3.06 | B:ARG 372[ NH2] |
| 23 | A:THR 185[ O ] | 2.65 | B:VAL 163[ N ] |
| 24 | A:ALA 187[ O ] | 3.26 | B:ASP 161[ N ] |
| 25 | A:TRP 189[ O ] | 2.86 | B:SER 172[ N ] |
| 26 | A:GLY 214[ O ] | 3.45 | B:ASN 158[ ND2] |
| 27 | A:ASN 291[ O ] | 3.15 | B:ASN 291[ ND2] |
| 28 | A:GLU 292[ O ] | 3.08 | B:ASN 291[ N ] |
| 29 | A:PRO 300[ O ] | 2.70 | B:TRP 309[ N ] |
| 30 | A:GLN 301[ O ] | 2.73 | B:GLY 310[ N ] |
| *Salt Bridges* | | | |
| 1 | A:ARG 210[ NE ] | 2.90 | B:GLU 153[ OE1] |
| 2 | A:ARG 210[ NH2] | 3.34 | B:GLU 153[ OE1] |

| | | | |
|---|---|---|---|
| 3 | A:ARG 210[ NH2] | 3.05 | B:GLU 153[ OE2] |
| 4 | A:LYS 289[ NZ ] | 2.88 | B:ASP 256[ OD2] |
| 5 | A:ARG 294[ NE ] | 3.44 | B:ASP 290[ OD2] |
| 6 | A:ARG 294[ NH2] | 3.29 | B:ASP 290[ OD2] |
| 7 | A:LYS 317[ NZ ] | 3.11 | B:GLU 269[ OE1] |
| 8 | A:GLU 363[ OE1] | 3.98 | B:LYS 169[ NZ ] |

**B vs C**

*Hydrogen bonds*

| | | | |
|---|---|---|---|
| 1 | B:MET 119[ N ] | 3.06 | C:ALA 147[ O ] |
| 2 | B:GLN 120[ NE2] | 3.00 | C:GLU 149[ OE1] |
| 3 | B:ILE 121[ N ] | 2.88 | C:GLU 149[ O ] |
| 4 | B:ILE 125[ N ] | 2.72 | C:VAL 151[ O ] |
| 5 | B:ALA 187[ N ] | 2.89 | C:ASP 161[ O ] |
| 6 | B:HIS 188[ ND1] | 2.93 | C:SER 172[ OG ] |
| 7 | B:TRP 189[ N ] | 3.05 | C:PRO 170[ O ] |
| 8 | B:TRP 189[ NE1] | 2.98 | C:LYS 169[ O ] |
| 9 | B:TYR 206[ OH ] | 2.61 | C:GLU 153[ OE1] |
| 10 | B:ARG 210[ NE ] | 2.57 | C:GLU 153[ OE1] |
| 11 | B:ARG 210[ NH2] | 3.41 | C:GLU 153[ OE2] |
| 12 | B:ARG 280[ N ] | 3.62 | C:TYR 263[ OH ] |
| 13 | B:ARG 280[ NE ] | 3.00 | C:GLU 267[ OE2] |
| 14 | B:ARG 280[ NH1] | 3.05 | C:SER 246[ OG ] |
| 15 | B:ARG 280[ NH2] | 3.81 | C:GLU 267[ OE1] |
| 16 | B:ARG 280[ NH2] | 2.89 | C:ASP 244[ OD2] |
| 17 | B:TYR 295[ OH ] | 2.75 | C:ASP 256[ OD1] |
| 18 | B:TYR 295[ OH ] | 3.50 | C:THR 253[ OG1] |
| 19 | B:GLN 301[ NE2] | 3.88 | C:ILE 296[ O ] |
| 20 | B:LYS 317[ NZ ] | 3.08 | C:GLU 267[ O ] |
| 21 | B:GLN 117[ O ] | 3.35 | C:SER 145[ OG ] |
| 22 | B:MET 119[ O ] | 3.05 | C:GLU 149[ N ] |
| 23 | B:ILE 125[ O ] | 3.15 | C:GLU 153[ N ] |
| 24 | B:MET 126[ O ] | 3.78 | C:ARG 372[ NE ] |
| 25 | B:THR 185[ O ] | 2.76 | C:VAL 163[ N ] |
| 26 | B:ALA 187[ O ] | 3.47 | C:ASP 161[ N ] |
| 27 | B:TRP 189[ O ] | 2.88 | C:SER 172[ N ] |
| 28 | B:GLY 214[ O ] | 3.76 | C:ASN 158[ ND2] |
| 29 | B:ASN 291[ O ] | 3.03 | C:ASN 291[ ND2] |

| | | | |
|---|---|---|---|
| 30 | B:GLU 292[ O ] | 2.88 | C:ASN 291[ N ] |
| 31 | B:PRO 300[ O ] | 2.87 | C:TRP 309[ N ] |
| 32 | B:GLN 301[ O ] | 2.88 | C:GLY 310[ N ] |

*Salt Bridges*

| | | | |
|---|---|---|---|
| 1 | B:ARG 210[ NE ] | 2.57 | C:GLU 153[ OE1] |
| 2 | B:ARG 210[ NE ] | 3.95 | C:GLU 153[ OE2] |
| 3 | B:ARG 210[ NH2] | 3.42 | C:GLU 153[ OE1] |
| 4 | B:ARG 210[ NH2] | 3.41 | C:GLU 153[ OE2] |
| 5 | B:ARG 280[ NE ] | 3.00 | C:GLU 267[ OE2] |
| 6 | B:ARG 280[ NH2] | 3.81 | C:GLU 267[ OE1] |
| 7 | B:ARG 280[ NH2] | 3.67 | C:GLU 267[ OE2] |
| 8 | B:ARG 280[ NH2] | 2.89 | C:ASP 244[ OD2] |
| 9 | B:LYS 289[ NZ ] | 3.01 | C:ASP 256[ OD2] |
| 10 | B:LYS 317[ NZ ] | 3.10 | C:GLU 269[ OE1] |
| 11 | B:GLU 363[ OE1] | 3.90 | C:LYS 169[ NZ ] |

**B vs D**

*Hydrogen Bonds*

| | | | |
|---|---|---|---|
| 1 | B:PHE 176[ N ] | 3.15 | D:SER 111[ O ] |
| 2 | B:THR 185[ OG1] | 3.43 | D:GLN 195[ OE1] |
| 3 | B:ARG 338[ NH2] | 3.27 | D:ASP 198[ O ] |
| 4 | B:ARG 338[ NH2] | 3.34 | D:ASP 199[ OD1] |
| 5 | B:ARG 365[ NH1] | 3.04 | D:ASP 199[ OD1] |
| 6 | B:ARG 365[ NH2] | 3.06 | D:ASP 199[ OD2] |
| 7 | B:SER 346[ OG ] | 2.52 | D:GLU 348[ OE1] |
| 8 | B:GLU 348[ N ] | 3.79 | D:GLU 348[ OE1] |
| 9 | B:ARG 350[ N ] | 3.07 | D:ARG 350[ O ] |
| 10 | B:ASP 173[ OD2] | 3.08 | D:SER 104[ N ] |
| 11 | B:GLU 171[ OE2] | 3.37 | D:SER 104[ N ] |
| 12 | B:PHE 176[ O ] | 3.34 | D:SER 114[ OG ] |
| 13 | B:GLU 149[ OE2] | 3.79 | D:GLN 117[ NE2] |
| 14 | B:GLU 344[ OE2] | 2.86 | D:ARG 194[ NH2] |
| 15 | B:GLU 344[ OE1] | 3.55 | D:ARG 347[ NH1] |

*Salt Bridges*

| | | | |
|---|---|---|---|
| 1 | B:ARG 338[ NH2] | 3.34 | D:ASP 199[ OD1] |
| 2 | B:ARG 338[ NE ] | 3.36 | D:ASP 199[ OD1] |
| 3 | B:ARG 365[ NH1] | 3.04 | D:ASP 199[ OD1] |
| 4 | B:ARG 365[ NH2] | 3.69 | D:ASP 199[ OD1] |

| | | | |
|---|---|---|---|
| 5 | B:ARG 365[ NH1] | 3.66 | D:ASP 199[ OD2] |
| 6 | B:ARG 365[ NH2] | 3.06 | D:ASP 199[ OD2] |
| 7 | B:ASP 173[ OD2] | 3.08 | D:SER 104[ N ] |
| 8 | B:GLU 171[ OE2] | 3.37 | D:SER 104[ N ] |
| 9 | B:GLU 344[ OE2] | 3.75 | D:ARG 194[ NH1] |
| 10 | B:GLU 363[ OE1] | 3.94 | D:ARG 194[ NH1] |
| 11 | B:GLU 363[ OE2] | 3.57 | D:ARG 194[ NH1] |
| 12 | B:GLU 344[ OE2] | 2.86 | D:ARG 194[ NH2] |
| 13 | B:GLU 363[ OE2] | 3.46 | D:ARG 194[ NH2] |
| 14 | B:GLU 344[ OE1] | 3.55 | D:ARG 347[ NH1] |
| 15 | B:GLU 344[ OE2] | 3.91 | D:ARG 347[ NH2] |

**B vs E**

*Hydrogen Bonds*

| | | | |
|---|---|---|---|
| 1 | B:SER 104[ N ] | 3.41 | E:GLU 171[ OE2] |
| 2 | B:SER 104[ N ] | 3.06 | E:ASP 173[ OD2] |
| 3 | B:SER 114[ OG ] | 3.64 | E:PHE 176[ O ] |
| 4 | B:ARG 194[ NH2] | 2.54 | E:GLU 344[ OE1] |
| 5 | B:GLN 195[ NE2] | 3.27 | E:THR 185[ OG1] |
| 6 | B:ARG 347[ NH1] | 3.62 | E:GLU 344[ OE1] |
| 7 | B:ARG 347[ NH2] | 2.66 | E:GLU 344[ OE2] |
| 8 | B:SER 111[ O ] | 3.01 | E:PHE 176[ N ] |
| 9 | B:ASP 198[ O ] | 2.95 | E:ARG 338[ NH2] |
| 10 | B:ASP 199[ OD1] | 3.12 | E:ARG 338[ NE ] |
| 11 | B:ASP 199[ OD1] | 3.05 | E:ARG 338[ NH2] |
| 12 | B:ASP 199[ OD1] | 3.01 | E:ARG 365[ NH1] |
| 13 | B:ASP 199[ OD2] | 3.06 | E:ARG 365[ NH2] |
| 14 | B:GLU 348[ OE2] | 3.32 | E:ARG 347[ N ] |
| 15 | B:GLU 348[ OE2] | 2.83 | E:SER 346[ OG ] |
| 16 | B:GLU 348[ OE2] | 3.55 | E:GLU 348[ N ] |
| 17 | B:ARG 350[ O ] | 3.12 | E:ARG 350[ N ] |

*Salt Bridges*

| | | | |
|---|---|---|---|
| 1 | B:SER 104[ N ] | 3.41 | E:GLU 171[ OE2] |
| 2 | B:SER 104[ N ] | 3.06 | E:ASP 173[ OD2] |
| 3 | B:ARG 194[ NE ] | 3.69 | E:GLU 363[ OE2] |
| 4 | B:ARG 194[ NH1] | 3.29 | E:GLU 344[ OE1] |
| 5 | B:ARG 194[ NH1] | 3.80 | E:GLU 363[ OE2] |
| 6 | B:ARG 194[ NH2] | 3.95 | E:GLU 344[ OE2] |

| | | | |
|---|---|---|---|
| 7 | B:ARG 194[ NH2] | 2.54 | E:GLU 344[ OE1] |
| 8 | B:ARG 194[ NH2] | 3.73 | E:GLU 363[ OE2] |
| 9 | B:ARG 347[ NH1] | 3.62 | E:GLU 344[ OE1] |
| 10 | B:ARG 347[ NH2] | 2.66 | E:GLU 344[ OE2] |
| 11 | B:ARG 347[ NH2] | 3.40 | E:GLU 344[ OE1] |
| 12 | B:ASP 199[ OD1] | 3.12 | E:ARG 338[ NE ] |
| 13 | B:ASP 199[ OD1] | 3.05 | E:ARG 338[ NH2] |
| 14 | B:ASP 199[ OD1] | 3.01 | E:ARG 365[ NH1] |
| 15 | B:ASP 199[ OD1] | 3.80 | E:ARG 365[ NH2] |
| 16 | B:ASP 199[ OD2] | 3.48 | E:ARG 365[ NH1] |
| 17 | B:ASP 199[ OD2] | 3.06 | E:ARG 365[ NH2] |

**B vs F**

*Hydrogen Bonds*

| | | | |
|---|---|---|---|
| 1 | B:LEU 105[ N ] | 2.81 | F:GLN 191[ O ] |
| 2 | B:GLN 117[ NE2] | 3.37 | F:GLN 120[ OE1] |
| 3 | B:SER 193[ N ] | 3.06 | F:LEU 105[ O ] |
| 4 | B:GLN 191[ O ] | 2.66 | F:LEU 105[ N ] |
| 5 | B:LEU 105[ O ] | 3.20 | F:SER 193[ N ] |

**Table S7. Strains used in this study.**

| Strain | Description | Reference |
|---|---|---|
| 594 | *E. coli.* Laboratory strain | |
| DH5α | *E. coli.* Laboratory strain | |
| C1a | *E. coli.* Laboratory strain | |
| DH5α λ*pir* | *E. coli.* Laboratory strain | (Hanahan, 1983) |
| S17 MFD*pir* | *E. coli.* Laboratory strain | (Ferrières et al., 2010) |
| EDL933 | *E. coli.* Natural isolate | (Wells et al., 1983) |
| JP23851 | *E. coli.* Natrual isolate RHBSTW-00139 | (AbuOun et al., 2021) |
| JP23772 | *E. coli.* Natural isolate GN02175 | (Bethke et al., 2020) |
| JP22739 | *K. pneumoniae.* Natrual isolate DSM30104 | |
| JP24901 | *Enterobacter cloacae subsp. Cloacae.* Natrual isolate ATCC 13047 | A gift from Dr. Avinash Shenoy |
| JP18938 | *Salmonella enterica subsp. enterica serovar Typhimurium.* LT2 ΔFels-1 ΔGifsy-2 ΔGifsy-1 ΔFels-2 | (Fillol-Salom et al., 2021) |
| JP25005 | *Enterobacter hormaechei.* Natrual isolate Ehh_4 | (Donà et al., 2023) |
| JP25008 | *Enterobacter hormaechei.* Natrual isolate Ehh_18 | (Donà et al., 2023) |
| JP24915 | *Citrobacter koseri.* Natrual isolate 2F8 | A gift from Dr. Thomas Clarke |
| JP24916 | *Citrobacter freundii.* Natrual isolate 2H5 | A gift from Dr. Thomas Clarke |
| JP24943 | *E. coli* EDL933 EcCIEDL933 1795-1796::cat | (Alqurainy et al., 2023) |
| JP23877 | Lytic phage for K. pneumoniae. Natural isolate K68 QB | A gift from Dr. Pilar Domingo-Calap |
| JP24850 | C1a HK022 lysogen | This work |
| JP12509 | 594 HK106 lysogen | (Alqurainy et al., 2023) |
| JP23887 | DSM30104 Δ*shv-1* | This work |
| JP23778 | DSM30104 KpCIDSM30104 Δ*orf17*::*tetA* | This work |
| JP25198 | DSM30104 ΔP1 KpCIDSM30104 Δ*orf17*::*tetA* | This work |
| JP24853 | DSM30104 Δ*shv-1* ΔP2 KpCIDSM30104 *Δorf17*::*tetA* | This work |
| JP24871 | DSM30104 Δ*shv-1* ΔP1 ΔP2 KpCIDSM30104 *Δorf17*::*tetA* | This work |
| JP24467 | DSM30104 Δ*shv-1* ΔP2 ΔKpCIDSM30104 | This work |
| JP24177 | DSM30104 ΔP1 ΔKpCIDSM30104 | This work |
| JP24460 | DSM30104 Δ*shv-1* ΔP1 ΔP2 ΔKpCIDSM30104 | This work |
| JP24891 | DSM30104 Δ*shv-1* ΔP2 P1 Δ*capsid* | This work |
| JP24929 | DSM30104 Δ*shv-1* ΔP2 KpCIDSM30104 Δ*orf17*::*tetA* Δ*capsid* | This work |
| JP24918 | DSM30104 Δ*shv-1* ΔP2 P1 Δ*capsid* KpCIDSM30104 Δ*orf17*::*tetA* | This work |
| JP24946 | DSM30104 Δ*shv-1* ΔP2 P1 Δ*tail tubular* KpCIDSM30104 Δorf17::*tetA* | This work |
| JP24699 | GN02175 ΔEcCIGN02175 | This work |
| JP25235 | JP24699 KpCIDSM30104 Δorf17::tetA | This work |
| JP24927 | GN02175 EcCIGN02175::*tetA* | This work |
| JP25170 | *Enterobacter hormaechei* Ehh_18 ΔEhCIEhh_18 | This work |
| JP25149 | JP25170 EcCIGN02175::*tetA* | This work |
| JP25021 | DH5α pBAD18 EcCIEDL933 *adaptor* and *connector* | This work |
| JP24949 | DH5α pBAD18 EcCIGN02175 *adaptor* and *connector* | This work |
| JP24950 | DH5α pBAD18 KpCIDSM30104 *adaptor* and *connector* | This work |
| JP24930 | DSM30104 Δ*shv-1* ΔP2 KpCIDSM30104 Δ*adaptor* and | This work |

| | | |
|---|---|---|
| | *connector* Δ*orf17*::*tetA* | |
| JP24953 | JP24930 pBAD18 | This work |
| JP24954 | JP24930 pBAD18 EcCIGN02175 *adaptor* and *connector* | This work |
| JP24955 | JP24930 pBAD18 KpCIDSM30104 *adaptor* and *connector* | This work |
| JP25022 | JP24930 pBAD18 EcCIEDL933 *adaptor* and *connector* | This work |
| JP25009 | C1a HK022 *gp05** (*capsid**) | This work |
| JP25010 | C1a HK022 Δ *major tail* | This work |
| JP24888 | C1a pBAD18-*kmR* | This work |
| JP24970 | C1a HK106 pBAD18-*kmR* | This work |
| JP24971 | C1a HK022 pBAD18-*kmR* | This work |
| JP22479 | 594 HK106 EcCIEDL933 1795-1796::*cat* | (Alqurainy et al., 2023) |
| JP25155 | 594 HK106 EcCIEDL933 1795-1796::cat *capsid* D257K | This work |
| JP25150 | 594 HK106 EcCIEDL933 1795-1796::cat *capsid* K290D | This work |
| JP25153 | 594 HK106 EcCIEDL933 1795-1796::cat *capsid* R345D | This work |
| JP25156 | 594 HK106 EcCIEDL933 1795-1796::cat *capsid* ΔLoop (Δ232-247) | This work |
| JP25152 | 594 HK106 EcCIEDL933 1795-1796::cat *capsid* D295K | This work |
| JP25151 | 594 HK106 EcCIEDL933 1795-1796::cat *capsid* E319K | This work |
| JP19758 | 594 HK106 gp05* EcCIEDL933 1795-1796::cat | (Alqurainy et al., 2023) |
| JP24598 | JP19758 pBAD18 EcCIIHE3034 *alpA* | This work |
| JP25146 | 594 HK106 gp05* EcCIEDL933 1795-1796::cat *capsid* ΔLoop (Δ232-247) | This work |
| JP25172 | JP25146 pBAD18 EcCIIHE3034 *alpA* | This work |
| JP24957 | DH5α pBAD18 HK97 *capsid* | This work |
| JP24958 | DH5α pBAD18 HK97 *capsid* K289D | This work |
| JP24959 | DH5α pBAD18 HK97 *capsid* K317D | This work |
| JP24960 | DH5α pBAD18 HK97 *capsid* D256K | This work |
| JP24961 | DH5α pBAD18 HK97 *capsid* R294D | This work |
| JP22372 | 594 HK106 *gp05** | (Alqurainy et al., 2023) |
| JP19807 | JP22372 pBAD18 | (Alqurainy et al., 2023) |
| JP24982 | JP22372 pBAD18 HK97 *capsid* | This work |
| JP24983 | JP22372 pBAD18 HK97 *capsid* K289D | This work |
| JP24984 | JP22372 pBAD18 HK97 *capsid* K317D | This work |
| JP24985 | JP22372 pBAD18 HK97 *capsid* D256K | This work |
| JP24986 | JP22372 pBAD18 HK97 *capsid* R294D | This work |
| JP24987 | 594 pBAD18 HK97 *capsid* | This work |
| JP25236 | DH5α λ*pir* pKNG101 ΔKpCIDSM30104 | This work |
| JP25233 | S17 MFD*pir* pKNG101 ΔKpCIDSM30104 | This work |
| JP25237 | DH5α λ*pir* pKNG101 ΔP1 | This work |
| JP25234 | S17 MFD*pir* pKNG101 ΔP1 | This work |
| JP24180 | DH5α λ*pir* pKNG101 ΔP2 | This work |
| JP23910 | S17 MFD*pir* pKNG101 ΔP2 | This work |
| JP24694 | DH5α λ*pir* pKNG101 ΔEcCIGN02175 | This work |
| JP24696 | S17 MFD*pir* pKNG101 ΔEcCIGN02175 | This work |
| JP25113 | DH5α λ*pir* pKNG101 ΔEhCIEhh_18 | This work |
| JP25114 | S17 MFDpir pKNG101 ΔEhCIEhh_18 | This work |
| JP25000 | DH5α λ*pir* pKNG101 HK022 Δ*major tail* | This work |
| JP25001 | S17 MFD*pir* pKNG101 HK022 Δ*major tail* | This work |
| JP23898 | DH5α pWRG99-*kmR* | This work |

**Table S8. Plasmids and oligonucleotides used in this study.**

| Plasmid | Description | Reference |
|---|---|---|
| **pBAD18** | $Amp^R$. Expression vector under the control of L-arabinose. | (Guzman et al., 1995) |
| **pBAD18-*kmR*** | $Km^R$. Expression vector under the control of L-arabinose. | (Fillol-Salom et al., 2022) |
| **pWRG99** | $Amp^R$. Thermosensitive plasmid with Red system of lambda phage and I-*Sce*I endonuclease under control of tetracycline-inducible promoter ($P_{tetA}$) | (Blank et al., 2011) |
| **pWRG99-*kmR*** | $Km^R$. Thermosensitive plasmid with Red system of lambda phage and I-*Sce*I endonuclease under control of tetracycline-inducible promoter ($P_{tetA}$) | This work |
| **pWRG717** | $Amp^R$, $Km^R$. pBluescript II SK+ derivative, aph resistance cassette and I-*Sce*I cleavage site | (Hoffmann et al., 2017) |
| **pKD4** | Plasmid with $Km^R$ flanked by *FRT* sites. | (Datsenko and Wanner, 2000) |
| **pCP20** | $Amp^R$, $Cm^R$. Thermosensitive plasmid with thermal induction of FLP recombinase synthesis. | (Cherepanov and Wackernagel, 1995) |
| **pKNG101** | $Strep^R$. A transmissible plasmid for site-directed mutagenesis of Gram-negative bacteria, replicating only in strains expressing Pir protein. Its *sacB* gene makes it lethal in the presence of sucrose for counterselection. | (Kaniga, Delor and Cornelis, 1991) |
| **pJP2267** | pBAD18 EcCIIHE3034 *alpA* | This work |
| **pJP3018** | pBAD18 HK97 *capsid* | This work |
| **pJP3019** | pBAD18 HK97 *capsid* K289D | This work |
| **pJP3020** | pBAD18 HK97 *capsid* K317D | This work |
| **pJP3021** | pBAD18 HK97 *capsid* D256K | This work |
| **pJP3022** | pBAD18 HK97 *capsid* R294D | This work |
| **pJP3023** | pBAD18 EcCIEDL933 *adaptor* and *connector* | This work |
| **pJP3024** | pBAD18 EcCIGN02175 *adaptor* and *connector* | This work |
| **pJP3025** | pBAD18 KpCIDSM30104 *adaptor* and *connector* | This work |
| **pJP3026** | pKNG101 ΔKpCIDSM30104 | This work |
| **pJP3027** | pKNG101 ΔP1 | This work |
| **pJP3028** | pKNG101 ΔP2 | This work |
| **pJP3029** | pKNG101 ΔEcCIGN02175 | This work |
| **pJP3030** | pKNG101 ΔEhCIEhh_18 | This work |
| **pJP3031** | pKNG101 HK022 Δ*major tail* | This work |

| Mutagenesis | Primers | Sequence (5'-3') |
|---|---|---|
| **DSM30104 Δ*shv-1*** | **DSM30104-SHV1::tetA-1m** | GCGTGCTTAGGCAGGGCTAGATATTGATTATTCGAAATAAAAGATGACAACGCTGTTAATCACTTTACTTTTATC |
| | **DSM30104-SHV1::tetA-2c** | CACGCCTGCGAGGTGCTACGAGCCGGATAACGCGCGCGGCCACCGCCGGGGGTTATCAAGAGGGTCATTATATTTC |
| | **DSM30104-AtetA-1m** | GCGTGCTTAGGCAGGGCTAGATATTGATTATTCGAAATAAAAGATGACAATGTGTAGGCTGGAGCTGCTTCG |
| | **DSM30104-AtetA-2c** | CACGCCTGCGAGGTGCTACGAGCCGGATAACGCGCGCGGCCACCGCCGGGGGTCCATATGAATATCCTCCTTAG |
| **KpCIDSM30104 Δorf17::*tetA*** | **KpCIDSM30104-Aorf17::tet-1m** | GTCCTGAACAAGTCAGGTGAGGCTGCTCAGGCTCTGGATACTCTCGGGCTTCGCTGTTAATCACTTTACTTTTATC |
| | **KpCIDSM30104-Aorf17::tet-2c** | CTGCTGGATAGCGATAACAGACTGAGCAATAGCCGCGGCTTTCTGAACTGCGGTTATCAAGAGGGTCATTATATTTC |
| **KpCIDSM30104 Δ*adaptor* and *connector*** | **KpCIDSM-Aadaptor-FRT-1m** | GGAAGTCACTGTGAGCAAGGGCAATAAGGATGGTGACGAATGAAAGCCGGTGTGTAGGCTGGAGCTGCTTCG |
| | **KpCIDSM-Aconnector-FRT-2c** | GTACACCCCCACGTCACGATAGACAGACCACAGCGCTGAGACAGCAAGAGGGGTCCATATGAATATCCTCCTTAG |
| **KpCIDSM30104 Δ*capsid*** | **KpCIDSM-Acapsid-FRT-1m** | CTCCAAGCGCTGGTTTCACGTCTCTACATTAATTGATACGGAAACCACTCCTGTGTAGGCTGGAGCTGCTTCG |
| | **KpCIDSM-Acapsid-FRT-2c** | GACTCCACAGAGGGCAGAAAGGGGCCGAAGCCCCTCACACGTCGAATCAGGGTCCATATGAATATCCTCCTTAG |
| **P1 Δ*capsid*** | **31290-Acapsid-FRT-1m** | GAAAAACGATCTGGTCGAAGCGTTTTATCCAAGCAAAAAATAGTGAGGTGTAATCTGTGTAGGCTGGAGCTGCTTCG |
| | **31290-Acapsid-FRT-2c** | GCCTGAGCTGCGGAAAAAAGCGCTTGCTGGTCGAGCAGCATGATCCCCCCCTCCATATGAATATCCTCCTTAG |
| **P1 Δ*tail tubular*** | **31290-Atail-tubular-1m** | GAGGATCTGCTGATGGCTTATTCACTGGTGCAGCCGTCGCTTGCCGGCGGTGTGTAGGCTGGAGCTGCTTCG |
| | **31290-Atail-tubular-2c** | GAATTGCGCTGATATGCTCGTCAGTAACGCTGACTATCTCAACTTTTCGCATAGGTCCATATGAATATCCTCCTTAG |
| **EcCIGN02175::*tetA*** | **Ec144-tetA-End-1m** | ATATGAGCGACAGTCACAAAGAGCGACTGGCTTTATTTAATGGGATTTGACGCTGTTAATCACTTTACTTTTATC |
| | **Ec144-tetA-End-2c** | GCGCCTTTTTTATTATTGTTTATACAGATGTTTATACAGCTTTGGCTGAACAAACAAAAAAGGTTATCAAGAGGGTCATTATATTTC |
| **EcCIEDL933 *capsid*::*kmR* first recombination** | **EcCIEDL-capsid-717-1m** | GATGTGGACATCCTTTCATGGCTGACGGGTGAGATTGGCGACACCTTCACGAGGGTTTCCCAGTCACGAC |
| | **EcCIEDL-capsid-717-2c** | CCTGTGGCAGTTCCAGAATCTTGATCGCGTTCGAATCCACCACGCCACCGTGCTTCCGGCTCGTATGTTG |
| **EcCIEDL933 *capsid* D257K** | **EcCIEDL933-capsid-11m** | CCGGTCTATGCGTACCCG |
| | **EcCIEDL933-capsid-12c** | GACCGGATTTCCATACTCTTC |
| | **EDL-capsid-D257K-1m** | CTCTGGCGTGGACATCTGCGAAACCGCTGATCGACCTGAAATTTGCATTACG |

| | | |
|---|---|---|
| | **EDL-capsid-D257K-2c** | CGTAATGCAAATTTCAGGTCGATCAGCGGTTTCGCAGATGTCCACGCCAGAG |
| **EcCIEDL933 *capsid* K290D** | **EcCIEDL933-capsid-11m** | CCGGTCTATGCGTACCCG |
| | **EcCIEDL933-capsid-12c** | GACCGGATTTCCATACTCTTC |
| | **EDL-capsid-K290D-1m** | CTCCACGACGGCGGCAAAACTTCAGAAGGTGGATAACGCGAACGGTGATTAC |
| | **EDL-capsid-K290D-2c** | GTAATCACCGTTCGCGTTATCCACCTTCTGAAGTTTTGCCGCCGTCGTGGAG |
| **EcCIEDL933 *capsid* R345D** | **EcCIEDL933-capsid-11m** | CCGGTCTATGCGTACCCG |
| | **EcCIEDL933-capsid-12c** | GACCGGATTTCCATACTCTTC |
| | **EDL-capsid-R345D-1m** | GTTGATCACGAAACAGGTGTTGATACCAGACCGGACAACCTC |
| | **EDL-capsid-R345D-2c** | GAGGTTGTCCGGTCTGGTATCAACACCTGTTTCGTGATCAAC |
| **EcCIEDL933 *capsid* ΔLoop (Δ232-247)** | **EcCIEDL933-capsid-11m** | CCGGTCTATGCGTACCCG |
| | **EcCIEDL933-capsid-12c** | GACCGGATTTCCATACTCTTC |
| | **EDL-A232-247-1m** | CCGTACCCCGTGCAGAGTCCGAATCTCTGGCGTGGAC |
| | **EDL-A232-247-2c** | GTCCACGCCAGAGATTCGGACTCTGCACGGGGTACGG |
| **EcCIEDL933 *capsid* D295K** | **EcCIEDL933-capsid-11m** | CCGGTCTATGCGTACCCG |
| | **EcCIEDL933-capsid-12c** | GACCGGATTTCCATACTCTTC |
| | **EDL-capsid-D295K-1m** | GGTGAAGAACGCGAACGGTAAATACATCTGGCGCGACCG |
| | **EDL-capsid-D295K-2c** | CGGTCGCGCCAGATGTATTTACCGTTCGCGTTCTTCACC |
| **EcCIEDL933 *capsid* E319K** | **EcCIEDL933-capsid-11m** | CCGGTCTATGCGTACCCG |
| | **EcCIEDL933-capsid-12c** | GACCGGATTTCCATACTCTTC |
| | **EDL-capsid-E319K-1m** | GCCTTCCGGTCGAATATCTGAAATTTATGCCTGATAACGTTATTGCCC |
| | **EDL-capsid-E319K-2c** | GGGCAATAACGTTATCAGGCATAAATTTCAGATATTCGACCGGAAGGC |

| Plasmid | Primers | Sequence (5'-3') |
|---|---|---|
| pJP3023 | Sakai-ECs1593-1mS | ACGCGTCGACTCAAATCCGGCGTTATGTGCC |
| | Sakai-ECs1594-2cH | CCCAAGCTTTTCTGTGCATCGTCTTAATGG |
| pJP3024 | EcCI-addaptor-1mS | ACGCGTCGACGCAGGAAGTGAAGGTCAGTAC |
| | EcCI-connector-2cH | CCCAAGCTTGCACCGGCGCATTGGTTG |
| pJP3025 | KpCI-adaptor-1mS | ACGCGTCGACGGCATGGAAGCAGGAAGT |
| | KpCI-connector-2cH | CCCAAGCTTCTCAGTGCAACGGCGTAAG |
| pJP3018 | HK97-capsid-1mS | ACGCGTCGACCTCTGCTTCAGAGCATCAAATC |
| | HK97-capsid-2cH | CCCAAGCTTCCATCATGAGCCAGAAGAG |
| pJP3021 | HK97-capsid-1mS | ACGCGTCGACCTCTGCTTCAGAGCATCAAATC |
| | HK97-capsid-2cH | CCCAAGCTTCCATCATGAGCCAGAAGAG |
| | HK97-capsid-D256K-1m | CCACCGGCGACACCCGCGCTAAAATTATCGCTCACGCTATTTATC |
| | HK97-capsid-D256K-2c | GATAAATAGCGTGAGCGATAATTTTAGCGCGGGTGTCGCCGGTGG |
| pJP3019 | HK97-capsid-1mS | ACGCGTCGACCTCTGCTTCAGAGCATCAAATC |
| | HK97-capsid-2cH | CCCAAGCTTCCATCATGAGCCAGAAGAG |
| | HK97-capsid-K289D-1m | GACTGGCACAACATCGCGTTGCTGGACGACAATGAAGGCCGCTATATC |
| | HK97-capsid-K289D-2c | GATATAGCGGCCTTCATTGTCGTCCAGCAACGCGATGTTGTGCCAGTC |
| pJP3022 | HK97-capsid-1mS | ACGCGTCGACCTCTGCTTCAGAGCATCAAATC |
| | HK97-capsid-2cH | CCCAAGCTTCCATCATGAGCCAGAAGAG |
| | HK97-capsid-R294D-1m | GTTGCTGAAAGACAATGAAGGCGACTATATCTTCGGTGGTCCTCAGGC |
| | HK97-capsid-R294D-2c | GCCTGAGGACCACCGAAGATATAGTCGCCTTCATTGTCTTTCAGCAAC |
| pJP3020 | HK97-capsid-1mS | ACGCGTCGACCTCTGCTTCAGAGCATCAAATC |
| | HK97-capsid-2cH | CCCAAGCTTCCATCATGAGCCAGAAGAG |
| | HK97-capsid-K317D-1m | GCTTGCCAGTGGTTCCGACTGACGCGCAGGCCGCCGGCACC |
| | HK97-capsid-K317D-2c | GGTGCCGGCGGCCTGCGCGTCAGTCGGAACCACTGGCAAGC |
| pJP3031 | Amajortail-1mB | CGCGGATCCACGGCGTAGATAATGACGTG |
| | HK022-Amajortail-2cNot | AAGGAAAAAAGCGGCCGCCCGGTACTTTGCCCAGAG |
| | HK022-Amajortail-1m | CCGGCCTTGAGCCGGTTTTTTTATGACCGGAGATAACAAATCAGGGCGGCAACGCCCTTATTAATC |
| | HK022-Amajortail-2c | GATTAATAAGGGCGTTGCCGCCCTGATTTGTTATCTCCGGTCATAAAAAAACCGGCTCAAGGCCGG |
| pJP3029 | EcCI144-FlankingL-1mS | ACGCGTCGACCACCGTTGAACGTGTAGTTC |
| | EcCI144-FlankingR-2cB | CGCGGATCCTGCCTTTGGCAGAGGTTAAG |

| | | |
|---|---|---|
| **pJP3030** | **AEhh18-3m** | CCGTGGTACCCTGCGTAAGAAACTGAAAAAATACGGCATGAACTAATTTCGATTAGCTAACTGCTTG |
| | **AEhh18-4c** | CAAGCAGTTAGCTAATCGAAATTAGTTCATGCCGTATTTTTTCAGTTTCTTACGCAGGGTACCACGG |
| | **AEhh18-1mS** | ACGCGTCGACGACTATACGGGAGCTGATGCTC |
| | **AEhh18-2cSpe** | GGACTAGTCAGCCATATTCAGTGCGGTATAG |
| **pJP3029** | **KpClDSM30104-FlankingL-3mS** | ACGCGTCGACGACGGAGCTGTTTAATGAAATG |
| | **KpClDSM30104-FlankingR-4cB** | CGCGGATCCAGAGCTGTGCTCGACTATAC |
| **pJP3027** | **DSM30104-31290-FlankL-5mS** | ACGCGTCGACAAAGCACACGTCATCAAATC |
| | **DSM30104-31290-FlankR-6cSpe** | GGACTAGTCTTTATTGTTACGGCATGATCC |
| **pJP3028** | **DSM30104-P2-FlankL-6mS** | ACGCGTCGACGCACAAAGGCAACATCATG |
| | **DSM30104-P2-FlankR-7cSpe** | GGACTAGTCGGATAGCCTGAGTTTAGGC |
| | **DSM-AP2-1m** | CTCAGATTATGTCAGAACGGTGATTTGAACAAGCAGGTTC |
| | **DSM-AP2-2c** | GAACCTGCTTGTTCAAATCACCGTTCTGACATAATCTGAG |
| **pWRG99-*kmR*** | **pWRG99-Gibson-KanR-1m** | GCACTTTTCGGGGAAATGTGGGCGGTTTTATGGACAGCAAG |
| | **pWRG99-Gibson-KanR-2c** | CTTGCTGTCCATAAAACCGCCCACATTTCCCCGAAAAGTGC |
| | **pWRG99-Gibson-KanR-3m** | GCGGGACTCTGGGGTTCGAAACTGTCAGACCAAGTTTACTC |
| | **pWRG99-Gibson-KanR-4c** | GAGTAAACTTGGTCTGACAGTTTCGAACCCCAGAGTCCCGC |
| | **pWRG99-3m** | ggtcccatgGATTCTTCGTCT |
| | **pWRG99-4c** | GTGCCAATACCAGTAGAAAC |
| **pJP2267** | **IHE3034-alpA-4mS** | ACGCGTCGACTCCGATCTCATATATCTACAGC |
| | **IHE3034-alpA-2cH** | CCCAAGCTTCGGTCGCCTTTGTTTTATATG |

| Southern blot | Primers | Sequence (5’-3’) |
|---|---|---|
| **EcClGN02175 probe** | **KpClDSM30104-3m** | CTTATATAGGGAAAATGGCGCT |
| | **KpClDSM30104-Int-4c** | GCAGTGTCCATGGTTCAGAATG |
| **KpClDSM30104 probe** | **KpClDSM30104-primase-1m** | TCAGCATGATTGCAAATAAG |
| | **KpClDSM30104-primase-2c** | CACTGAACGTCATAGCATTG |
| **P1 probe** | **31290-Int-1m** | GTATCACACTGGCTGAAGCC |
| | **31290-Int-3c** | CTGCCTTTCATCCTGTCAG |
